# Spatial transcriptomics of T and B cell receptors uncovers lymphocyte clonal dynamics in human tissue

**DOI:** 10.1101/2022.11.22.516865

**Authors:** Camilla Engblom, Kim Thrane, Qirong Lin, Alma Andersson, Hosein Toosi, Xinsong Chen, Embla Steiner, Giulia Mantovani, Michael Hagemann-Jensen, Sami Saarenpää, Mattias Jangard, Jakob Michaëlsson, Johan Hartman, Jens Lagergren, Jeff Mold, Joakim Lundeberg, Jonas Frisén

## Abstract

The spatial distribution of lymphocyte clones within tissues is critical to their development, selection, and expansion. We have developed Spatial Transcriptomics of VDJ sequences (Spatial VDJ), which maps immunoglobulin and TR antigen receptors in human tissue sections. Spatial VDJ captures lymphocyte clones matching canonical T, B, and plasma cell distributions in tissues and amplifies clonal sequences confirmed by orthogonal methods. We confirm spatial congruency between paired receptor chains, develop a computational framework to predict receptor pairs, and link the expansion of distinct B cell clones to different tumor-associated gene expression programs. Spatial VDJ delineates B cell clonal diversity, class switch recombination, and lineage trajectories within their spatial context. Taken together, Spatial VDJ captures lymphocyte spatial clonal architecture across tissues, which could have important therapeutic implications.

**One-Sentence Summary:** Spatial transcriptomics-based technology co-captures T and B cell receptors within their anatomical niche in human tissue.

## Main Text

B and T cells are critical to health; they respond to infections and transformed cells, regulate tissue homeostasis, and maintain immunological memory. T and B cells’ targeted reactivity is determined by their clonally unique antigen receptors expressed by each cell and its progeny. Single-cell technologies permit the study of antigen receptors (i.e., T and B cell receptors) at a cellular level, but lack spatial resolution. Current spatial transcriptomics (ST) methods, which locate gene expression in tissues, do not retain full-length antigen receptors. Due to fragmentation for short-read sequencing, the Complementarity Determining Region 3 (CDR3), which is needed to define lymphocyte clonality, is absent or highly reduced in the final Visium sequencing library (fig. S1A). Spatial analysis of lymphocyte clonality could link specific antigen receptors to proximal tissue elements, such as tumor-associated, self, or foreign antigens, which could help identify and harness antigen-specific clones for therapy. An area of high interest across research fields, several techniques have recently been developed to spatially resolve antigen receptors within tissues, including PCR-based amplification of T cell receptors from either ST (*1*) or Slide-seq libraries (*2*) or, through deep sequencing, capture of rare longer IG transcripts containing the CDR3 regions (that remained post-fragmentation) in ST libraries (*3*). However, methodologies to comprehensively map full-length antigen receptors and their lineage relationships within tissues in a high-throughput, user-friendly manner are currently lacking.

Here, we developed Spatial Transcriptomics for VDJ sequences (Spatial VDJ) that simultaneously maps full-length T and B cell antigen receptor genes in human tissue (Fig. 1A). Spatial VDJ is an extension of ST described by Ståhl et al. (*4*), then modified into Visium Spatial Gene Expression (10x Genomics). Spatial transcriptomics relies on 3’-targeted barcoding of polyadenylated RNAs, resulting in whole-transcriptome gene expression data from 55µm-sized spots in a tissue section (fig. S1A) (*1*). Visium gene expression (hereafter Spatial GEX) libraries from human tonsil tissue showed ample B and T cell receptor constant gene expression (fig. S1B to D). Still, they constituted a small fraction of the total Spatial GEX library (fig. S1E). Prior to the fragmentation steps, the CDR3 region could be PCR amplified from spatially barcoded full-length Visium cDNA (fig. S1F), showing that the CDR3 sequence is present before fragmentation. These results support the feasibility of sequencing the CDR3 region and the spatial barcode to determine both clonality and antigen receptor transcripts’ original location.

**Fig. 1.**
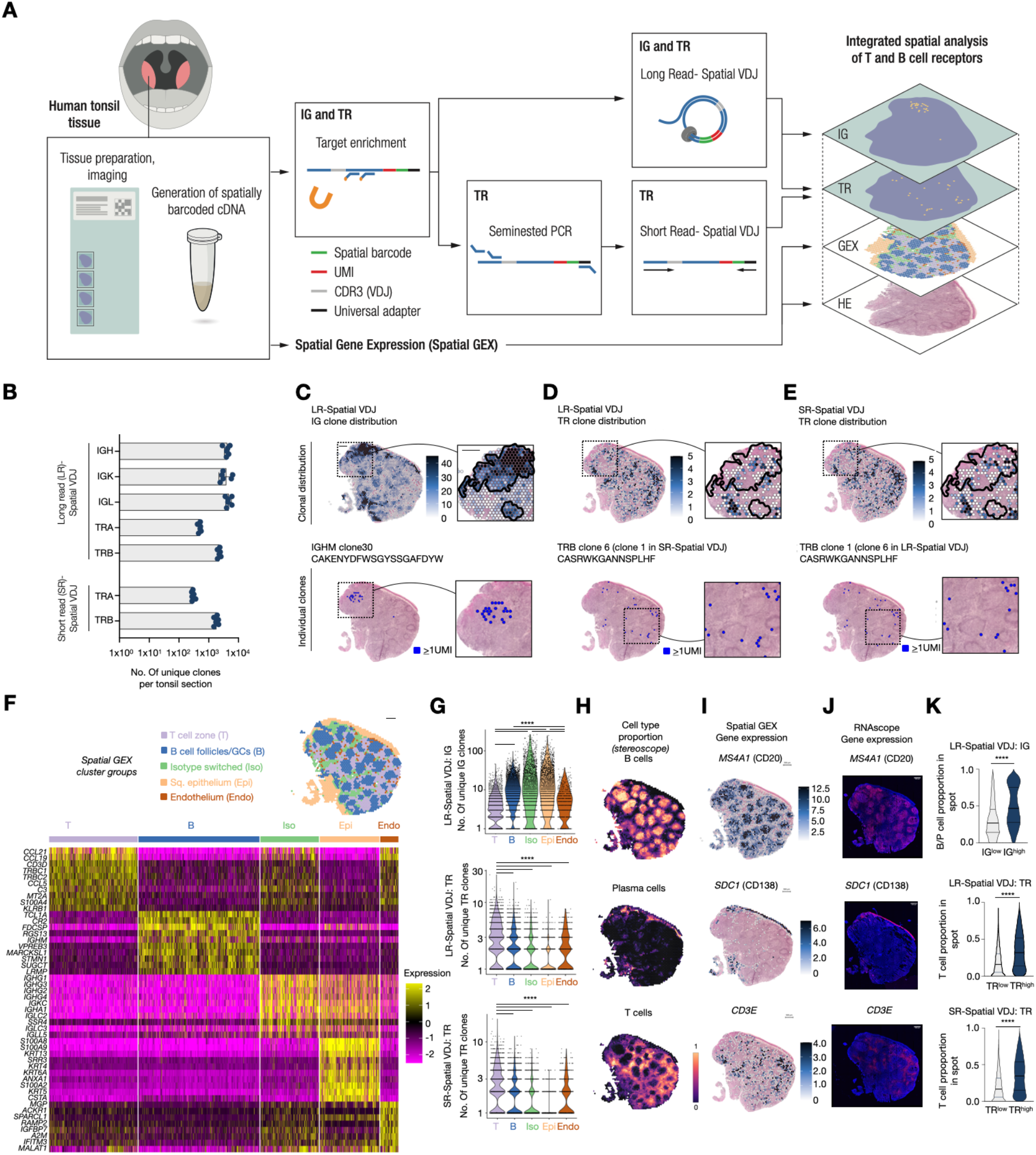
Spatial transcriptomics for VDJ sequences (Spatial VDJ) maps antigen receptors in human tonsil matching canonical T and B cell distributions. (**A**) Overview of Long read (LR) and Short read (SR) Spatial VDJ method in human tonsil tissue. (**B**) Unique IGH, IGK, IGL, TRA, and TRB clonotype sequences for each tonsil section (n=6). (**C**-**E**) Unique IG (C) and TR (D) clonal receptor sequence distribution for LR-Spatial VDJ, and unique TR (E) clonal distribution for SR-Spatial VDJ. Scale bar = 500µm. Black line represents regions with high IG clonal diversity. (**F**) Spatial GEX cluster group spatial distribution on tonsil tissue (top) and enriched genes (bottom). (**G**) IG or TR clonal diversity across Spatial GEX cluster groups. (Kruskal-Wallis test, Dunn’s multiple comparison test; **** = *p*<0.0001). (**H**) Visualization of major cell type distribution within tonsil defined by single cell deconvolution. (**I**) Spatial gene expression of canonical B (top), plasma (middle), and T (bottom) cell markers (*MS4A1, SDC1*, and *CD3E*, respectively). (**J**) RNAscope of genes shown in I. Scale bar = 500µm. (**K**) Cell type proportion for spots containing high or low clonotype counts (IG or TR) (Mann Whitney test; **** = *p*<0.0001).

We developed two Spatial VDJ strategies from antigen receptor-enriched full-length spatial cDNA (Fig. 1A): i) Long read Spatial VDJ (LR-Spatial VDJ) that generates spatially barcoded libraries of full-length immunoglobulin (IG genes) and T cell (TR genes) antigen receptors and ii) Short-Read Spatial VDJ (SR-Spatial VDJ), for TR genes only, which employs a two-step semi-nested PCR-based approach using CDR3-adjacent V primers (Methods, Table S1). For both approaches, we used target enrichment with hybridization capture probes directed at the TR and IG constant regions to enrich for antigen receptor transcripts while preserving the spatial barcode, UMI, and the full-length variable region domains, including the CDR3 (Fig. S2A) (*5, 6*). Hybridization capture significantly increased the TR and IG UMI counts compared to the non-enriched Spatial GEX library and preserved the IG and TR transcript distribution (fig. S2B and C).

### Spatial VDJ permits high-fidelity T and B cell clonal mapping

We applied LR- and SR-Spatial VDJ to the human tonsil, due to its abundance and spatial compartmentalization of T and B cell lineages (Fig. 1A to E, and fig. S3A to D). We generated Spatial GEX libraries from six tonsil tissue sections from the same individual, with consistent gene and UMI counts (fig. S3A and B). Applying LR-Spatial VDJ to the same samples, we identified 62,533 unique IG (IGH, IGK, IGL) and TR (TRB, TRA) clonal sequences (fig. S4A), hereafter referred to as ‘clonotypes’ or ‘clones’ (Methods). We detected fewer TRA versus TRB clones, likely due to TRA’s lower transcriptional abundance (fig. S1B). The TR clone counts were slightly lower in SR versus LR-Spatial VDJ libraries (Fig. 1B), but the SR method recapitulated over half of the clonal sequences (fig. S4B and C). Additionally, clonal abundance correlated well between LR- and SR-Spatial VDJ datasets (fig. S4D) and clones shared between both datasets were more expanded (fig. S4E and F). We visualized the overall antigen receptor distribution (Fig. 1C to E) and individual expanded clones, which showed high inter-clonal diversity, in the tonsil tissue (fig. S5 and S6). Notably, the same TR clonal sequence, retrieved by either LR- or SR-Spatial VDJ, exhibited overlapping spatial distribution, demonstrating that these approaches yielded comparable results (fig. S6).

Relating the antigen receptor distribution to the tonsil architecture, we harnessed the matched tonsil Spatial GEX dataset. Unsupervised clustering rendered 13 clusters (i.e., groups of spots with similar gene expression) (Fig. S7A to G), which were annotated based on differentially expressed genes and grouped according to shared features (Fig. 1F and fig. S7E to G). IG clonal diversity was the highest within the ‘Squamous Epithelium’, ‘Isotype switched cells’, and ‘B cell follicles/GCs’ cluster groups (Fig. 1G). Clones of different IGH isotypes exhibited distinct regional distributions, which were consistent between Spatial VDJ and GEX (fig. S8A to E); e.g., IGHM and IGHD clones were primarily in B cell follicle/GC regions, whereas isotype-switched clones (i.e., IGHA or IGHG) were spread more evenly across multiple tissue regions (fig. S8A). These findings are consistent with naive or primary activated IGHM/D-expressing B cells being enriched in and around B cell follicles, including germinal centers (GCs), which are specialized sites of B cell maturation (*7*). In contrast, antibody-producing cells (i.e., plasmablasts and more long-lived plasma cells), which are often isotype-switched, typically reside near the tonsil epithelium, but also appear in GCs (*8*). Memory B cells, which are also often isotype switched, localize both in intra- and extra-follicular regions (*7*). Individual IGH clones of the same isotype showed different spatial distributions (fig. S8F and G), which could reflect the representation of clones at different activation and/or maturation stages. For TR, the clonal diversity was the highest in spots belonging to T cell zone-associated clusters (Fig. 1G); in line with the T cell zone as the main site for T cell and antigen-presenting cell interactions (*9, 10*). We found concordance between TR receptor chain expression in the Spatial VDJ and GEX datasets (fig. S8H to J). Expanded TR clones (TR clonotypes in >1 unique array spot) were typically scattered throughout the tissue and often found at the border of one, or several, follicles; these may be T follicular helper cells, which activate B cells at the T-B cell interface and facilitate their GC entry (fig. S6) (*11*). In sum, Spatial VDJ delineates diverse spatial distributions for both T and B cell clones consistent with canonical secondary lymphoid tissue architecture.

We next asked how the Spatial VDJ-mapped clones related to T, B, and plasma cell distributions in the tonsil. To infer the cell type distribution, we performed deconvolution of the Spatial GEX data using *stereoscope* (*12*), harnessing a comprehensive human tonsil scRNA-seq dataset (*13*) (Fig. 1H and fig. S9A to C). The *stereoscope* results were consistent with the Spatial GEX cluster groups (fig. S9C), and matched well-established B, plasma, and T cell-associated marker gene (*MS4A1, SDC1*, and *CD3E*, respectively) expression shown by Spatial GEX (Fig. 1I) and RNAscope on sections from the same tissue (Fig. 1J and fig. S9D). For spots that contained IG high (IG^high^) versus low (IG^low^) clonal content, the B and plasma cell proportion was significantly higher (Fig. 1K and fig. S9E). Similarly, TR^high^ spots had elevated T cell proportions compared to TR^low^ spots (Fig. 1K, and fig. S9F and G). Lymphocyte distribution within lymphoid tissues is carefully orchestrated by chemokine ligands and receptors (*14*), which displayed the expected spatial patterns in our datasets, and the antigen receptor distribution was concordant with known chemokine gradients within lymphoid tissues (fig. S10) (*14, 15*). Combined, Spatial VDJ-defined antigen receptor distribution correlated with lymphocyte spatial predictions, demonstrating our ability to define clonotypic distributions within distinct tissue microenvironments, such as densely packed human lymphoid tissue.

The spatial segregation of individual clonotypes was highly reproducible and the most abundant clones were detected across sequential tonsil sections (Fig. 2A and B and fig. S11A); however, TRA clones were rarely found across all six sections, likely due to lower *TRA* capture. Most clones appeared in one tonsil section, but expanded clones were detected across multiple or all sections, corresponding to higher UMI counts per clone (fig. S11B and C). These results were expected, since many T and B cells in secondary lymphoid organs are naive, non-expanded clones expressing a unique antigen receptor and unlikely to be in multiple array spots (and by extension, sections). To validate Spatial VDJ-defined clonal sequences, we analyzed nearby or adjacent sections from the same sample by a VDJ-targeted Smart-seq3 (SS3) protocol for bulk RNA, hereafter called ‘bulk SS3 VDJ’ (fig. S3A and fig. S12A to D, Methods). Overall, the bulk SS3 VDJ analysis reproduced the Spatial VDJ results. Of the IG clonotypes identified by LR-Spatial VDJ, we found almost half by bulk SS3 VDJ (Fig. 2C and fig. S12E). Bulk SS3 VDJ reproduced approximately a third of the TR clones captured by LR- and SR-Spatial VDJ. The clones detected by multiple methods had, on average, increased UMI counts, suggesting that expanded clones were more likely to be detected across methods (fig. S12F to H). Additionally, the clone fraction of shared clonal sequences correlated well between datasets (fig. 2D to F). Taken together, Spatial VDJ reproducibly detected expanded B and T cell clones, of which a substantial fraction was confirmed by an independent method, indicating high fidelity antigen receptor retrieval.

**Fig. 2.**
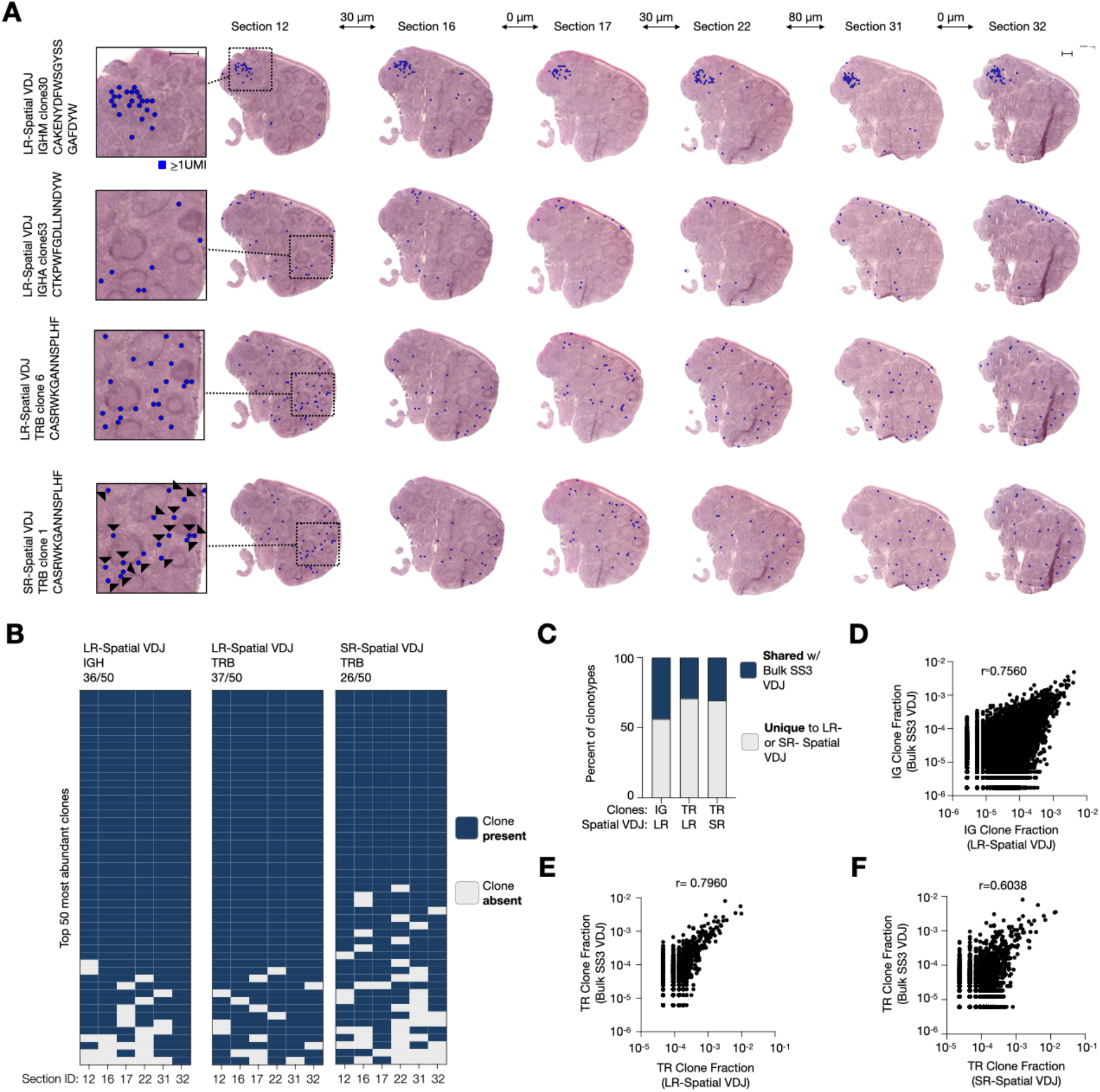
Spatial VDJ reproducibly delineates B and T cell clones across tissue sections and amplifies validated clonal sequences. (**A**) Spatial distribution of individual clones across tonsil sections. Distance between sections (orthogonal to the tissue plane) above the arrows. Matched TRB clonotype sequence (identical ntCDR3, Methods) captured by LR- and SR-Spatial VDJ is shown, concordant spots indicated with black arrow-heads (bottom row). Matched clones can have different clone IDs since numbering occurs independently within LR- and SR-Spatial VDJ datasets. Scale bar = 500µm. (**B**) Presence or absence of the top 50 IGH, IGK or TRB clonotypes (based on UMI count and sorted on number of sections) across tonsil sections. (**C**) Percent shared and not shared clonotype sequences between Spatial VDJ and Bulk SS3 VDJ analysis from separate tissue sections from the same tonsil sample (see fig. S3A). (**D** to **F**) Clone fraction of each shared (D) IG or (E and F) TR clonotypes between the datasets listed. Pearson correlation coefficient (r) is denoted.

### Spatial VDJ captures T and B cell clones in human tumor tissues

We next extended Spatial VDJ to human breast tumor tissue. Tumor infiltration by lymphocytes correlates with positive disease outcomes and treatment response across solid cancers, including breast cancer (*16, 17*). Mapping T and B cell clones within tumors could help identify and therapeutically harness the contribution of individual or groups of clones to tumor-associated immunity. We generated Spatial GEX and VDJ libraries from two untreated HER2-positive breast tumors (P1 and P2; Fig. 3A and B, fig. S13 to 17). To better capture the intra-patient diversity, each biopsy was divided into multiple regions (denoted as RegA, B etc), each of which contained tumor, tumor border, and adjacent non-tumor tissue. We identified thousands of unique IG clonal sequences and hundreds of unique clonal TR sequences per patient and section (fig. S16). Spatial GEX and VDJ libraries for adjacent tissue sections from the same biopsy region (fig. S14A and C, fig. S17) showed high concordance, whereas *separate* regions from the same patient were more heterogeneous, as revealed by clustering (Methods).

**Fig. 3.**
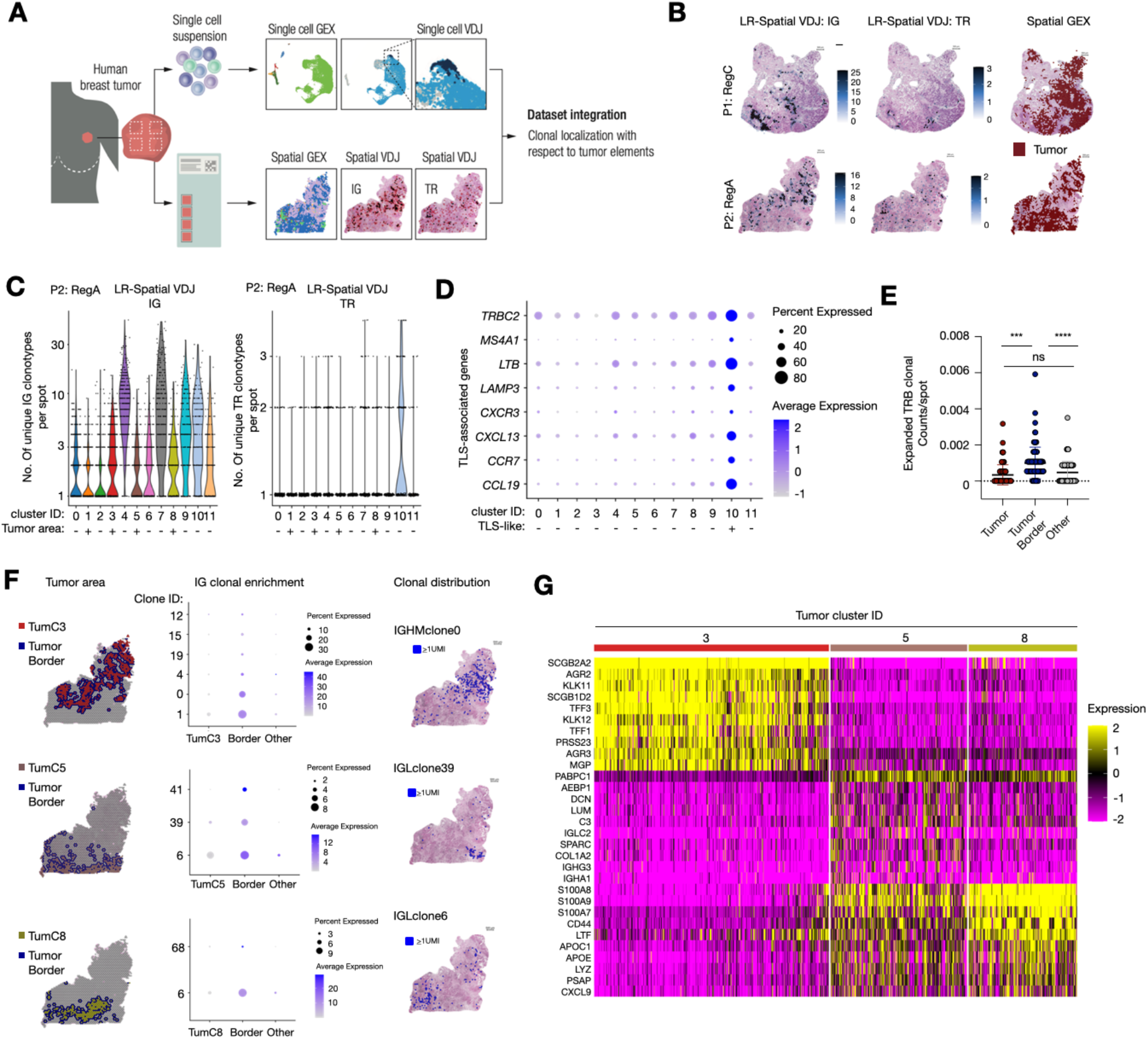
Distinct B cell lineage clones spatially segregate along different breast tumor areas. (**A**) Experimental overview. (**B**) Unique clonal distribution for LR-Spatial VDJ (IG, left, and TR, middle) and tumor area (right) for two biopsy regions from patient 1 (P1) and 2 (P2). (**C**) No. of unique IG (left) and TR (right) clonotypes per spot in different Spatial GEX clusters (‘+’ denotes tumor cluster). Data from all regions (RegA and B) from P2. (**D**) Fold enrichment of TLS-associated genes across Spatial GEX clusters. (**E**) UMI count per spot for expanded TRB clones across spots belonging to the Tumor, Tumor border, and other areas. Statistical significance was calculated using the Friedman’s test, then Dunn’s multiple comparison’s test; *** = *p*<0.001, **** = *p*<0.0001). (**F**) RegA Tumor 3, 5, and 8 clusters (Tum-c3, 5, and 8) and their surrounding borders visualized on P2 RegA (left). Dotplots showing fold change expression of significantly enriched IG clones per corresponding area (middle). Spatial distribution of representative IG clonotypes (right). See fig. S22. (**G**) Heatmap showing enriched genes for Tum-c3, 5, and 8. Abbreviations: TLS - Tertiary Lymphoid Structures; Reg - Region; Tum - Tumor

The IG and TR clonal diversity differed across spatial GEX clustering (Fig. 3C for P2 and fig. S17C for P1; Methods). IG receptor diversity was the highest in array spots classified as having an increased ‘Plasmablast’ presence in both patients as expected (c9-c12, c15 for P1; c4 and c7 for P1), based on single cell deconvolution analysis (fig. S18). These regions also showed enrichment for a ‘LAM2: APOE Myeloid’ signature in both patients (fig. S19), indicating a potential association between APOE^+^ tumor associated macrophages and tumor infiltrating B lineage cells, as observed previously in HER2 positive tumors (*18*). For TR clonotypes, the highest diversity was found in areas with signatures of immune infiltrates including both myeloid and T cell subset genes (c5 and c10 for P1; c10 for P2) (Fig. 3C, fig. S17C). These infiltrates bore hallmarks of mature Tertiary Lymphoid Structures (TLS) in P2 (c10), but not P1, including T follicular helper T cell-like CD4^+^ CXCL13-expressing cells, memory/naive B cells, and LAMP3^+^ dendritic cells (Fig. 3D and fig. S19) (*19*). Although B cells and plasmablasts tended to localize in different areas of the tissue, we detected shared IGH clones between the TLS-like area (c10), which was more enriched of B cells versus plasmablasts, and the plasmablast-rich part of the tissue (c4/c7) (fig. S17D), indicating the possibility that at least some of the tumor-associated plasmablasts were generated from TLS-activated B cells *in situ*. These findings highlight the potential for combined IG and TR profiling to uncover unique and coordinated roles for diverse populations of B and T cell clones in complex tissues like tumors.

We generated single cell GEX and VDJ clonotype datasets from CD45^+^ cells sorted from the same tumors (fig. S20 and S21) and confirmed approximately half of the Spatial VDJ-defined TR receptor chains by scRNA-seq (fig. S20G). The scVDJ TR clones that were shared with Spatial VDJ had more cells per clone, further supporting that Spatial VDJ captures expanded TR clones with high fidelity (fig. S20H). For IG clones, we confirmed only 3.0% and 2.9% of the LR-Spatial VDJ dataset by scRNA-seq (fig. S20G). These numbers are likely driven by the higher capture of *unique* IG clones by Spatial VDJ versus scRNA-seq (thousands versus hundreds) (fig. S16 and fig. S20F); it is possible that plasma cells are more sensitive to tissue digestion and sample processing for scRNA-seq, and that these cells are better captured by Spatial VDJ due to high IG receptor expression (*20*), increasing the likelihood of transcript capture by the ST array. Nonetheless, we find that Spatial VDJ successfully maps IG and TR clones in breast tumor tissue that can be confirmed by orthogonal analyses.

After encountering antigen and receiving proper activation cues, T cells proliferate and expand; using T cell clonotype expansion as a proxy for activation, we found that the UMI fraction from expanded clones (i.e., present in >1 array spot) was higher in the tumor border (i.e. tumor cluster-proximal spots), compared to either the tumor area or the spots that were neither the tumor area nor bordering the tumor (‘other’) (Fig. 3E). We sub-clustered the population of T cells in our scRNAseq dataset to assign phenotypic identities to each clonal population (fig. S21A to C). We observed that large clones tended to localize within the tumor and around the tumor border regardless of their dominant clonal phenotype, possibly reflecting a bias towards examining highly expanded clones (fig. S21D and E). Together, these findings highlight the potential to link single cell clonal phenotypes to spatial distribution in tissues.

Spatial colocalization of specific antigen receptors with discrete cancer-enriched regions may indicate tumor-antigen specificity towards certain tumor clones. For P2, we identified four distinct tumor clusters (fig. S15B and fig. S18B), three of which were detected primarily within RegA (Tum-c3, 5, and 8). Several IG clones were uniquely enriched in the tumor border surrounding Tum-c3, whereas a different set of IG clones were enriched within or adjacent to Tum-c5 and 8 (Fig. 3F and fig. S22A to C). Most clones were enriched in limited parts of the respective tumor border, indicating a localized antigen-binding response, e.g., IG clonal co-localization with distinct tumor sub-clones expressing unique antigens. In support of this idea, Tum-C3, 5, and 8 were all enriched for cancer cells (fig. S18B and fig. S19B), but the individual tumor clusters exhibited differently enriched genes (fig. 3G); Tum-c3 spots had elevated expression of genes involved in hormone-receptor signaling, including *TFF1* and *TFF3* (*21*), and expressed by breast glandular cells, e.g. *SCGB1D2* and *SCGB2A2* (*22*–*24*). In line with these findings, P2’s tumor had high estrogen and progesterone receptor positivity (fig. S13). Conversely, TumC5 and 8 shared many enriched genes, including tumorassociated *S100A7-*9, *CD44*, and *PABPC1*, which are involved in cell migration, proliferation, and cell-to-cell interactions (*25*–*28*), as well as genes, e.g., *LYZ* and *APOE* (*29*) expressed by macrophages, including those infiltrating tumors. Of note, *S100A7-9* are also highly expressed by neutrophils and could reflect neutrophil influx, which is not captured by the single cell deconvolution since this cell type is missing from the single cell GEX dataset (*30*). This intra-tumoral heterogeneity could ultimately result in differential tumor outgrowth and treatment response within the same tumor, e.g., by distinct IG clones driving antibody-dependent tumor cell killing in different locations of the tumor (*17*). These findings show the potential of combining an unsupervised methodology and Spatial VDJ to identify tumor-associated antigen receptor sequences enriched in specific tumor regions, implying potentially relevant tumor-reactive properties.

Paired B or T cell receptor sequences determine antigen binding and are needed for functional interrogation of antigen receptor specificity. Spatial VDJ captures individual antigen receptor transcripts, and thus cannot by itself define receptor pairs. However, since paired receptor chains are expressed by the same cell and therefore should co-localize in the tissue, we hypothesized that paired receptors could be linked by their spatial congruency. Using the matched single-cell VDJ dataset as the ground truth for paired receptor usage, we observed a remarkably similar spatial correlation for most IG pairs across biopsied regions (Fig. 4A and B and fig. S23A). We lacked sufficient data points for TR pairs to evaluate their spatial distribution in a statistically robust manner. Paired scVDJ data is not always available or does not capture all clones; for example, two Tum-c3-enriched clones (IGHMclone0 and IGLCclone1) listed above had paralleled spatial distributions (fig. S22A), but were missing from the scVDJ dataset. Therefore, we developed a computational framework to predict antigen receptor pairing, called *repair* (Methods), based on the Spatial VDJ data alone. We tested our model on data from P1 at different cut-offs of receptor chain abundance (minimal spot count per clone) and the ‘pairing score’ (fig. S23B and C), e.g., the confidence of the pairing. We achieved the maximum accuracy (95.8%) and number of correct pairs (n=23) when the IGH clonal chain had to be present in at least ten spots and the pairing score was at least 0.35 (Fig. 4C). At these settings, we obtained 157 *de novo* identified receptor pairs, i.e. receptor chains that were *not both* present in our ground truth data (Fig. 4D). Applying *repair* to P2, we found 25 predicted pairs (fig. S23D and E) with one out of two pairs present in the ground truth dataset verified at these settings. For the unverified pair, the ‘correct’ light chain confirmed by the scVDJ dataset (IGLclone39) only differed by one nucleotide from the light chain designated by *repair* (IGLclone45), suggesting that it may still be a ‘true’ receptor pair, albeit missing from our scVDJ dataset. The lower number of predicted pairs and overlap with the scVDJ data is due to P2’s overall lower number of IG clones and fewer analyzed tumor sections. Specifically addressing the enriched tumor-associated IG clones found in fig. S22, which for the most part were unmatched light chains, *repair* predicted a heavy partner chain for all the clones (except IGLclone39, as noted above), with a pairing score above the threshold as well as paired IGHMclone0 and IGCLclone1 (fig. S22A). These results indicate that *repair* in conjunction with Spatial VDJ could be used to predict *de novo* paired IG receptors in cryopreserved human tissues for further antibody studies.

**Figure 4:**
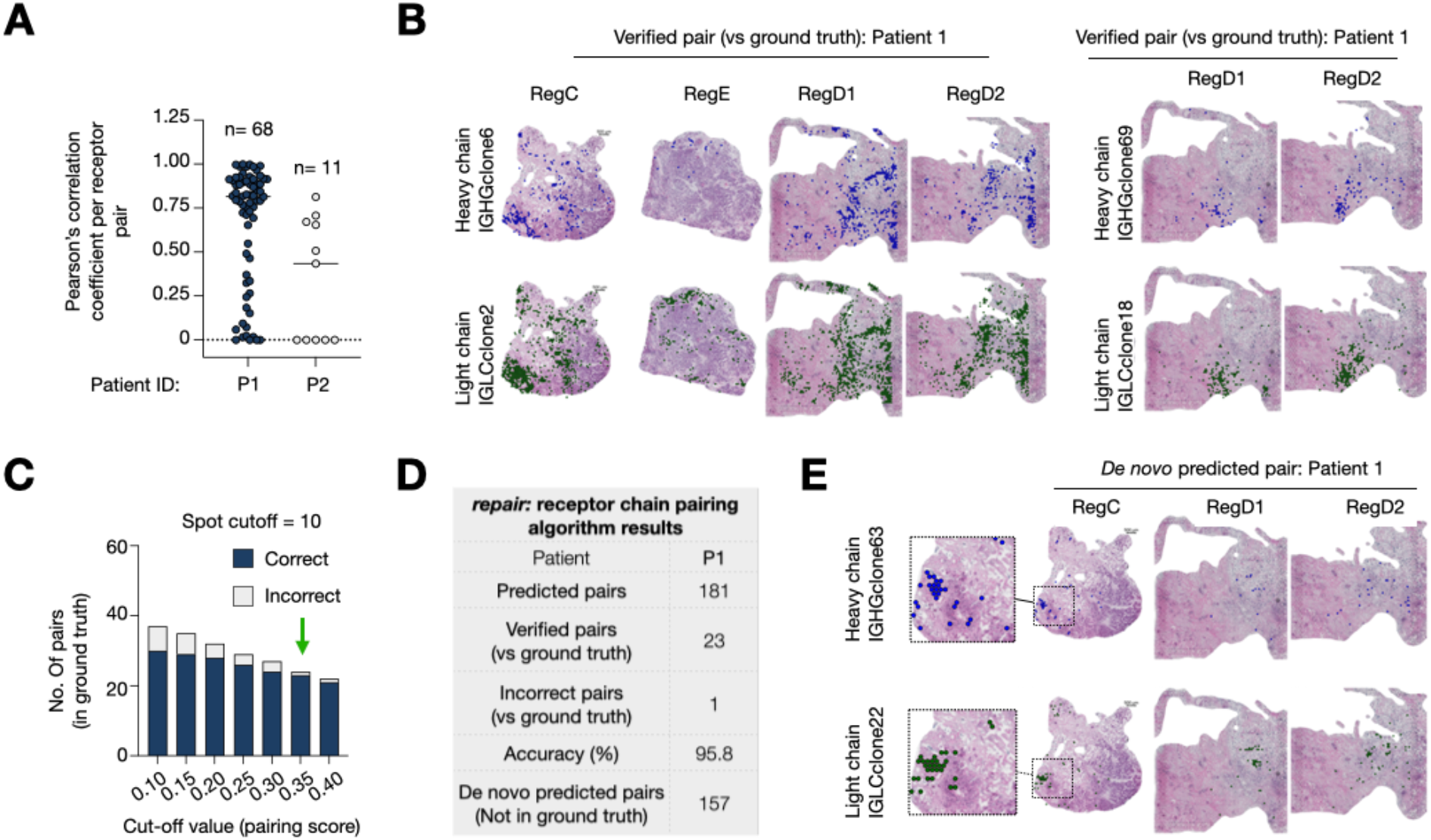
Computational framework predicts antigen receptor pairs based on their spatial distribution. (**A**) Pearson correlation coefficients (r) between the light versus heavy chain UMI count per spot for each IG receptor pair verified by single cell VDJ. (**B**) Spatial distribution of two verified IG light and heavy chain receptor pairs. (**C**) The number of correct versus incorrect pairs predicted by receptor pairing algorithm (*repair*) at different pairing score cut-offs. Each IGH chain was detected > 10 spots. Green arrow denotes selected pairing score cut-off for subsequent analyses. (**D**) Table showing results from *repair* (spot cut-off: n=10; pairing score=0.35). (**E**) Representative example of receptor pair predicted by *repair* absent from single cell VDJ data. Both pairs in (B) were predicted by *repair*. Reg=Region

### Spatial VDJ uncovers B cell spatial evolution

B cells undergo affinity-dependent selection and clonal expansion in GCs. Multiple clones can simultaneously cycle through a single GC generating a complex interplay between evolving clones, a process that dynamically changes throughout an immune response (*7*). We annotated individual B cell follicles, which included GCs, in human tonsils (Fig. 5A, fig. S7A and G, Methods). Thirty of 37 follicles (81.1%) were detected across all sections (fig. S24A). We grouped IGH clonotypes based on CDR3 similarity, resulting in 13,046 total “IGH clonal families”. We detected between approximately ten to almost one thousand IGH clonal families per follicle (Fig. 5B), and an average of 1.31 IGH clonal families (SD; +0.64) per spot per follicle (fig. S24B). The species richness (*Chao1* estimate), which partially accounts for incomplete sampling, varied between approximately thirty to almost three thousand per follicle (Fig. 5C). The largest clone within each follicle occupied between 2.38-34.0%, indicating that some follicles were more dominated by individual clonal families than others (fig. S24B). Of note, since each array spot contains multiple cells, a more dominant clonal family could be represented by multiple cells or a lower number of cells (e.g., plasmablast) with high receptor expression; regardless, no follicle was completely dominated by one single IGH clonal family (Fig. 5B). The IGH clonal family number per follicle positively correlated with follicle size (fig. S24C). Most clonal families belonged to a single follicle, but some appeared in multiple follicles, with 150 clonal families co-occurring in >4 follicles (Fig. 5D). Of those clonal families, approximately one fifth presented primarily in one ‘dominant’ follicle, i.e., defined as >75% of UMIs belonging to a single follicle (Fig. 5E and F, and fig. S24D), whereas other clonal families spread more evenly across multiple follicles (Fig. 5E and G). These findings suggest that a substantial number of IGH clonal families expand and presumably undergo affinity hypermutation in multiple GCs simultaneously, in line with laser microdissection studies in humans (*31*) and photoactivation studies in mice (*32*). These multi-follicular clonal families may be derived from recently activated GC B cells that migrated from one follicle to another during clonal expansion or from re-activated clonally related memory B cell clones that entered separate follicles simultaneously. Interpreting these results with caution, since the analysis is of a single individual, we found high IGH clonal diversity across human GCs, without any single clonal family taking over an entire GC, and both restricted and cross-follicular clonal expansion.

**Figure 5:**
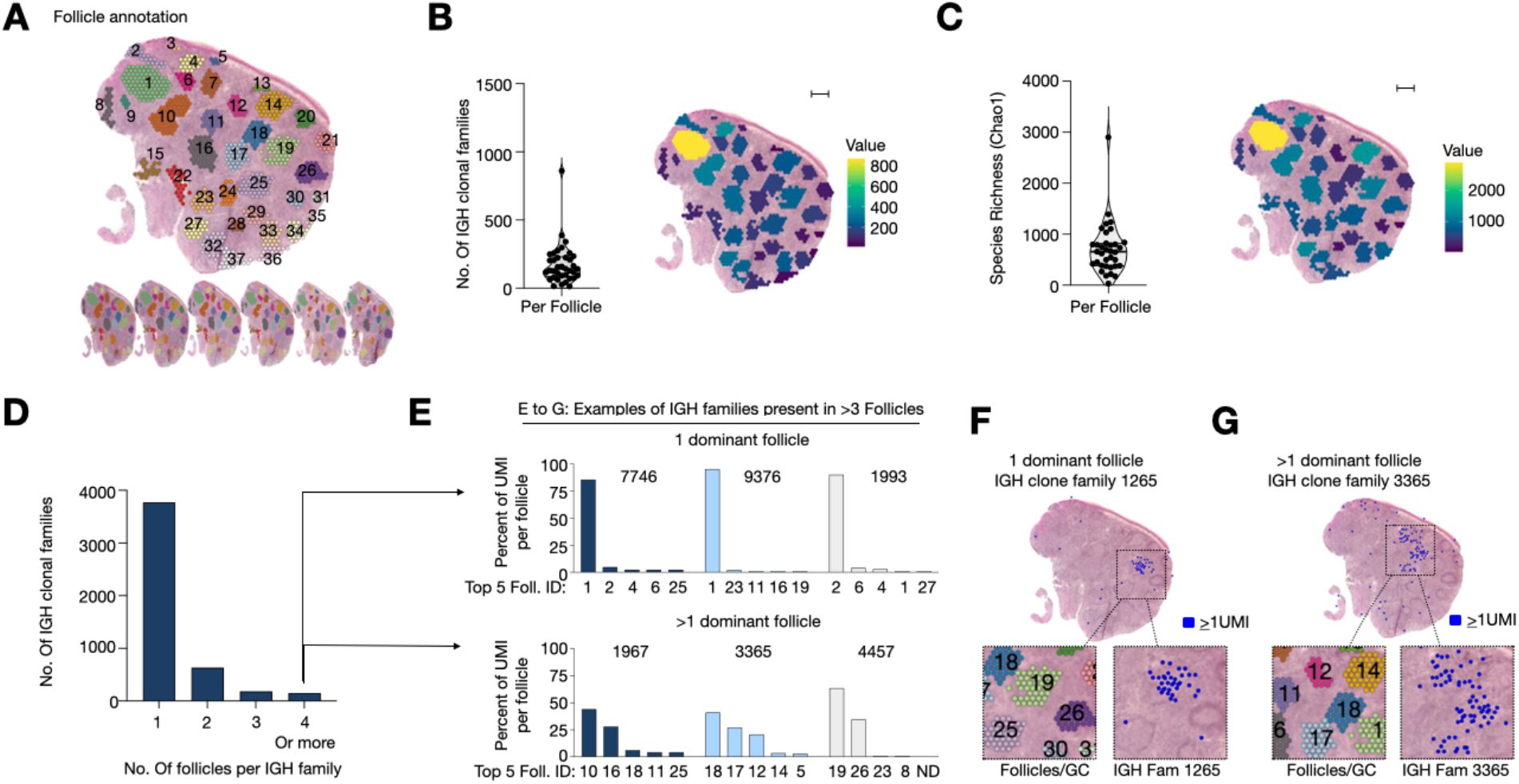
Spatial VDJ delineates IG clonal diversity in B cell follicles. (**A**) B cell follicle/GC annotation across human tonsil sections. (**B** and **C**) The IGH clonal families (B) total counts and (C) species richness (*Chao1*) (left) and spatial distribution (right) per follicle. (**D**) The distribution of each follicular IGH clonal family across follicles. (**E**) The percentage UMI distribution across the top 5 follicles for six representative IGH clonal families present in >4 follicles. ND: Not Detected. (**F** and **G**) Spatial distribution for a clonal family with (F) one or (G) multiple dominant follicles. Visualization on section 17. For all panels: data from 6 replicate sections are pooled.

B cell follicles contain molecularly and functionally distinct compartments, including the GC dark and light zone, enriched for somatic hypermutation and clonal expansion versus clonal selection, respectively (*7*). These regions were well represented in the Spatial GEX dataset and varied across follicles (fig. S25A to C); we annotated cluster c4 as dark zone, clusters c5/c1 as light zone, and cluster c7 as the mantle zone based on enriched genes and single cell deconvolution results (fig. S7F and fig. S25C); all but one follicle had evidence of an active GC (c1, 5, and/or 4). The total IGH clonal family count (i.e. diversity) was the highest in part of the light zone (c5) (fig. S25D); possibly reflecting on-going clonal selection in this area. Combined, Spatial GEX enables high-resolution demarcation of clonal composition across biologically relevant anatomical regions within lymphoid tissue.

Antibodies further diversify their functions by class switch recombination (CSR), i.e., replacing their constant region, which through its FC receptor interactions confers diverse effector functions. CSR was thought to mainly occur in GCs, but recent evidence indicates that CSR can occur outside GCs (*7, 13, 33*). Where CSR occurs in human lymphoid tissues remains debated. A cell undergoing or having recently undergone CSR should appear in our dataset as two (or more) different isotypes within one IGH clonal family present in the same array spot. Furthermore, if these transcripts were unmutated with respect to each other (i.e., had the same V gene sequence), they may have recently undergone CSR, here referred to as putative class switching events (pCSR). We detected 810 IGH clonal families containing multiple isotypes; of these, we detected 137 cases where identical V gene sequence alignments were matched with multiple isotypes, of which 17 occurred in the same spatial barcode (here referred to as pCSR events) (Fig. 6A), e.g., clonal family 2357 (fig. S26A to C). These 17 pCSR events all occurred extrafollicularly (Fig. 6A) and were mainly switches from IGHG1 to IGHG2 (Fig. 6B). One such event (in clone family 12298) occurred in an array spot with concurrent *AICDA* and *UNG* expression, the genes encoding critical enzymes mediating class switch recombination, providing further evidence that this represents a true class switching event (Fig. 6C). Combined, we find that Spatial VDJ detects putative class switch recombination events in human tissue and that these occur extra-follicularly.

**Figure 6:**
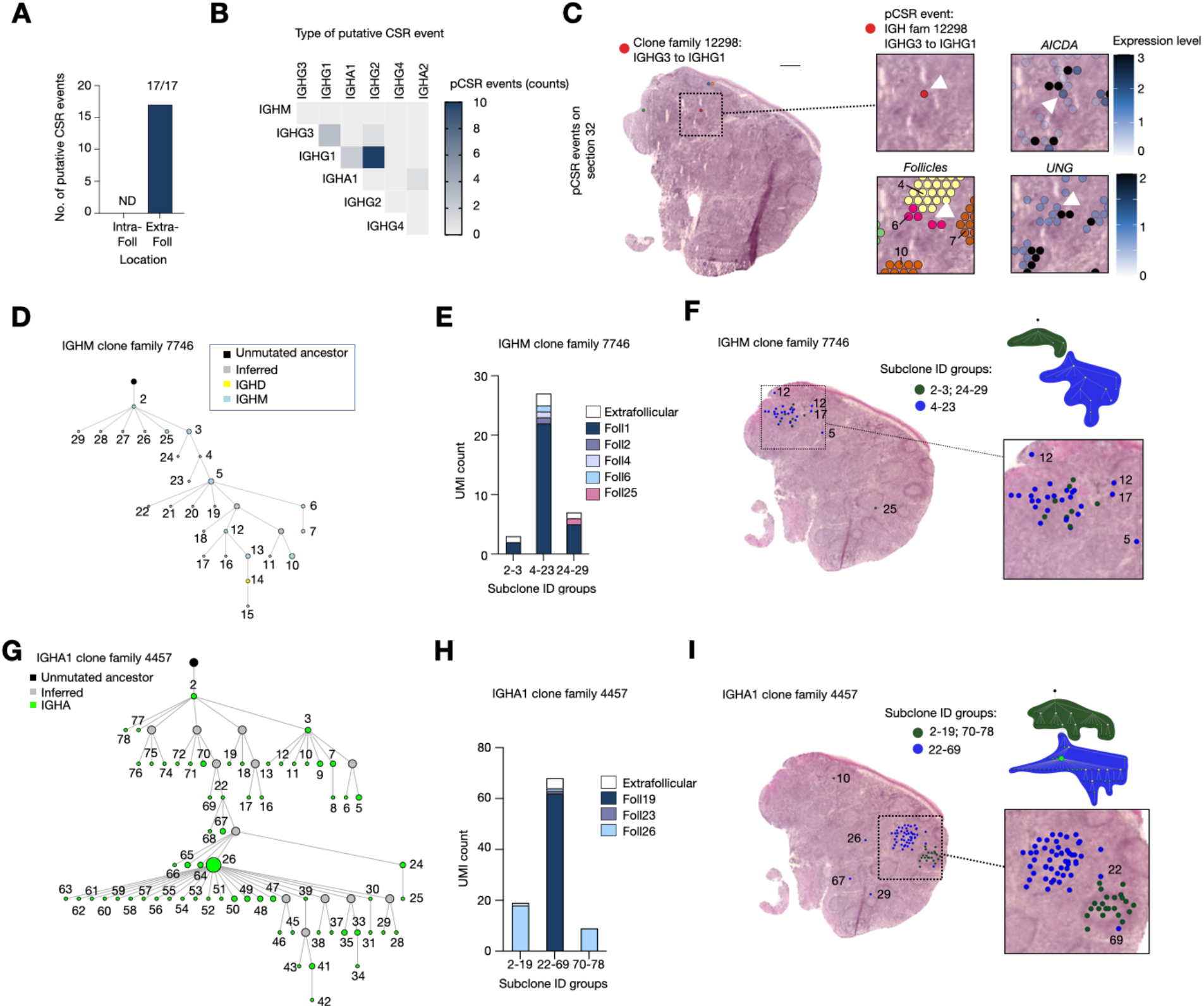
Spatial VDJ uncovers IG spatial clonal evolution in human lymphoid tissue. (**A**) The number of putative CSR (pCSR; n=17) occurring within (+) or outside (-) follicle/GC areas. (**B**) Type of pCSR events. (**C**) An individual pCSR event is visualized on section 32 (top, left zoom-in), its position relative to follicles (bottom, left), *AICDA* (top, right) and *UNG* (bottom, right) gene expression. White arrow denotes pCSR event in IGH family 12298. (**D**) Lineage tree of IGHM clone family 7746. Each node of the tree (‘sub-clone’) represents a unique V sequence and the size of the node is proportional to the number of unique reads in that node. All sub-clones are of the IGHM isotype (blue color). Inferred clones (not present in the data) are gray and lack a sub-clone ID. (**E**) Intra- and extra-follicular UMI distribution for IGHM clone family 7746 sub-clones. (**F**) Distribution of IGHM clone family 7746 on tonsil tissue. (**G**) Lineage tree of IGHA1 clone family 4457, as in (D). (**H**) Intra- and extra-follicular UMI distribution for IGHA1 clone family 4457 sub-clones. (**I**) Spatial distribution of IGHA1 clone family 4457. For all panels except (C): data from 6 replicate sections is pooled. For F and I: data from 6 sections is visualized on section 17. For (C): data is from section 32.

To better understand how IGH clonal families spatially segregate while undergoing somatic hypermutation, we generated phylogenetic trees for a subset of expanded IGH clonal families (Fig. 6D to I and fig. S26F to G). First, we explored clonal families that appeared primarily in one follicle, e.g., clone family 7746 (Fig. 6D to F) and the most prominent clone family 806 (fig. S26D to F). These clonal families were of the IGHM/D isotype, almost exclusively follicular, and had minor spreading to other follicles/extra-follicular areas by individual subclones (Fig. 6E and F, fig. S26D and G). Given their considerable clonal size, exclusive and concurrent *IGHM* and *IGHMD* expression, and follicular location, these clones are likely recently activated naive B cells undergoing somatic hypermutation. In contrast, clone family 4457 was an expanded, seemingly actively mutating family spread primarily across two follicles (Fig 6G to I). A branch of subclones (IDs 22-69) was almost exclusively found in Follicle 19, whereas the remaining subclones (IDs 2-19; 70-78) were only found in Follicle 26. These findings suggest that a subclone 22 (or a closely related, not detected subclone), likely a GC B cell, migrated from Follicle 19 to seed Follicle 26. Taken together, we provide evidence of clonal ‘escape’ to another GC during ongoing somatic hypermutation and map IG clonal evolution in human tissue.

## Discussion

Spatial VDJ allows high-resolution antigen receptor mapping within human lymphoid and tumor tissues. We validated clonal sequences captured by Spatial VDJ using orthogonal methods and developed a computational framework (*repair*) that both confirmed known and paired *de novo* IG receptor pairs. Furthermore, we presented two alternative methods (optimized for long- versus short-read sequencing) to capture TR clones that yielded comparable results. Distinct IG receptors spatially segregated across different tumor-associated areas within the same tumor, opening up the possibility to use Spatial VDJ as a first screen for tumor-reactive therapeutic antibodies. Spatial VDJ also uncovered B and plasma cell clonal dynamics across GCs in human tonsil tissue, located putative class switching events, and mapped clonal family trajectories with respect to tissue elements. Our approach yields a comprehensive high-resolution map of full-length human IG receptor evolution within their anatomical niches.

We foresee that Spatial VDJ will be applied across research fields to answer urgent clinically relevant questions. First, by delineating antigen receptors within their tissue context, Spatial VDJ could identify critical molecular and cellular elements that are required for lymphocyte clonal expansion, which are relevant to generate productive tumor-associated or vaccine-induced responses. Second, Spatial VDJ determines B cell clonal dynamics at an unprecedented scale and resolution in human secondary lymphoid tissue, which is an evolving organ, with germinal centers appearing, persisting, and disappearing dynamically during an immune response. Most of what we know about GCs as competitive niches are from elegant transgenic mouse models permitting lineage tracing of B cell clones, but these are not applicable to human tissue. We show that Spatial VDJ can appreciate the specialization of many distinct individual GCs in parallel and could be extended to understand how antibody responses are orchestrated in response to infections, vaccines, and autoimmune disorders. Finally, linking an antigen receptor sequence to its antigen reactivity can be highly challenging; by spatially co-localizing T or B cell clones with the transcriptional landscape (both host and non-host), Spatial VDJ could identify candidate antigen-specific clones, e.g., in autoimmunity or cancer. Identifying clones co-localizing with a variety of antigens, including tumor, viral, or self-antigens, could help identify T or B antigen receptor sequences for engineered T cells or antibody-based therapeutics.

## Acknowledgments

We acknowledge Marlon Stoeckius, Caroline Gallant and Katie Pfeiffer at 10x Genomics, Inc., for resources and technical support, Mattias Karlén’s graphical design of the experimental outlines, Helena Lönnqvist, Annelie Mollbrink, Sarantis Giatrellis, and Ludvig Larsson for expert assistance, and Ghamdan Al-Eryani, Frisén, Lundeberg, and Sandberg lab members for critical feedback on the study.

## Author contributions

CE, KT, and JMo initiated the study and performed the laboratory experiments relating to Spatial VDJ. CE and KT coordinated the study, analyzed the data, generated the figures, performed Spatial GEX, and wrote the manuscript with input from JMo, AA, QL, and HT. KT performed Spatial GEX and Spatial VDJ data processing and visualizations. CE and JM performed the single cell GEX/VDJ experiments. QL designed and implemented the code for the following analyses: SR-Spatial VDJ, Bulk SS3 VDJ, single cell GEX/VDJ, enrichment/depletion, lineage trajectories, and clonal overlap, and analyzed data, together with CE. AA performed the single cell deconvolution analysis, developed repair, and guided statistical/computational analysis. HT performed long read Spatial VDJ clonal analysis and scRNA-seq analysis. ES performed the RNAscope analysis. GM helped with laboratory experiments. SS helped with data analysis. JMo developed Bulk SS3 VDJ with MHJ. JMi coordinated tonsil sample acquisition. XC coordinated and performed initial processing of breast tumor samples under JH supervision. JH, JLa, Jmo, JLu, and JF were involved in study conceptualization and provided resources. JMo, JLu, and JF supervised the study. All authors read and approved the manuscript.

## Data and materials availability

All data, code, and images needed to reproduce the figures in the manuscript are deposited on zenodo (DOI: 10.5281/zenodo.7326539). References to data and code per figure panel are annotated in table S2. Raw sequencing data will be deposited on the Swedish EGA node and available upon request due to privacy concerns (GDPR compliance).

## Supplementary Materials

Materials and Methods

Figs. S1-S26

Table S1: Primer and probe sequences

Table S2: Figure panels linked to scripts and underlying datasets

