## SupplementaryMaterial for "Spatial transcriptomics of T and B cell receptors uncovers lymphocyte clonal dynamics in human tissue"

**This PDF file includes:**

Materials and Methods
Figs. S1 to S26

**Other Supplementary Materials for this manuscript include the following:**

Tables S1 to S2

### Materials and Methods

*Patient samples*

The tonsil samples were obtained from anonymous adult patients undergoing tonsillectomy due to obstructive sleep apnea syndrome. Permission to collect these unidentified tissues was obtained from the Ethical Review Authority in Stockholm, Sweden (2006/646-31/4, amendments 2015/1083-2, and 2017/1659-32). All patients gave informed consent prior to collecting tissue. From each tonsil a smaller piece (≈0.5 $cm^3$) was cut out, immediately embedded in OCT, placed on dry ice, and stored at -80°C.

*Breast tumor samples*

Breast cancer samples were obtained by Dr. Johan Hartman from the Department of Clinical Pathology and Cancer Diagnostics at Karolinska University Hospital, Stockholm, Sweden. Experimental procedures and protocols of the study were previously approved by the regional ethics review board (Etikprövningsnämnden) in Stockholm (2016/957-31, amendment 2017/742-32, 2021-00795, and 2022-05245-02). Breast tumor samples were collected from two different patients with untreated invasive ductal carcinomas during surgery. Histological evaluations of patient tumors were performed by pathologists for diagnostic purposes: tumor characteristics, including hormone receptor, HER2, and Ki67 expression are presented in a supplemental figure (fig. S13A). Both tumors were HER2 positive. For each tumor biopsy, different regions (*n=4-5*) were selected by the pathologists depending on the size of the tumor. From each region, tissue was isolated for immediate embedding in OCT and for gene expression analysis with spatial transcriptomics and antigen receptor analysis. We prepared Spatial GEX and VDJ libraries for three regions for patient no. 1 (P1) (fig. S13 and 14), and two regions for patient no. 2 (P2) (fig. S13 and S15). The remaining material for each tumor region was used for single-cell RNA sequencing analysis. Samples for spatial transcriptomics were immediately frozen and stored at -80°C until further analysis. The tissues obtained for single-cell analysis were processed directly.

*Preparation and sequencing of Spatial Gene Expression Libraries (Spatial GEX)*

Sections of fresh-frozen breast tumor and tonsil tissue were cut at 10 µm thickness and mounted onto slides from the Visium Spatial Gene Expression Slide & Reagent kit (10X Genomics). For the tonsil, six sections were analyzed covering 200 µm in the Z direction (orthogonal to the tissue plane) of the sample (fig. S3A). Sequencing libraries were prepared following the manufacturer's protocol (Document number CG000239 Rev A, 10x Genomics). Prior to imaging, coverslips were mounted on the slides according to the protocol's optional step *Coverslip Application & Removal*. Tissue images were taken at 20x magnification using the Metafer Slide Scanning platform (MetaSystems) and raw images were stitched with VSlide software (MetaSystems). Adaptations of the protocol were made in that the Hematoxylin staining time was reduced to 4 minutes and tissue permeabilization was performed for 12 minutes. Final libraries were sequenced on NextSeq2000 (Illumina) or NovaSeq6000 (Illumina).

*Preparation and sequencing of PCR amplification of T and B cell receptors (lacking the spatial barcode) from Spatial transcriptomics Gene Expression Libraries (Spatial GEX)*

Starting with Spatial GEX full-length cDNA libraries from two consecutive tonsil sections (Tonsil #1A and B) prepared as described above, 0.5 ng of each cDNA library was used as PCR input material. Triplicates of each sample and PCR reaction (TRB, IGHG, and IGHM) were performed. The primers in the PCRs targeted: a) the constant region of TRB, IGHG, and

IGHM (Table S1), and b) the V segments for TRB (*34*) and IGH (*35*). The constant primers were selected based on their proximity to the CDR3 region and the testing of various primers for each target in PCR optimization experiments. Both the forward and reverse primers were tagged with partial P5 and P7 handles that allowed subsequent Truseq indexing for Illumina sequencing. The PCR was performed using the KAPA HiFI Hotstart ready mix according to manufacturer's instructions with 30 amplification cycles. The most optimal annealing temperature had previously been established during optimization experiments.

After the initial PCR, we bead-purified the PCR product before carrying it forward into the PCR indexing reaction. Using Quantit to measure the DNA concentration of each sample, we also normalized the input material to the PCR indexing reaction (1.25ng/reaction). The indexing PCR was performed as previously described (*4*). Twenty-four unique indexes were used, with each cDNA library receiving a unique index (the TRB, IGHG, and IGHM products from the same cDNA library received the same indexing since the TCR and BCR clonotypes can be distinguished from each other bioinformatically using the constant primer sequence). After bead-purification of all reactions, all PCR reactions were pooled into one sample. This pooled library was run on a gel and a large band at around 500bp (expected product size) was gel-purified and sequenced (NovaSeq, 2x 150bp). The resulting data were de-multiplexed and the fastq files were analyzed using MIXCR (version 3.0.7) with the following command (*36*);

```
mixcr align --species hsa --report -p rna-seq
-OvParameters.geneFeatureToAlign=VTranscriptWithP
-OvParameters.parameters.floatingLeftBound=true
-OjParameters.parameters.floatingRightBound=false
-OcParameters.parameters.floatingRightBound=true
<R1Fastq>  <R2Fastq>  <SampleName>


mixcr assemble --report MIXCR/*.report --write-alignments
-OassemblingFeatures="[CDR3]"
-OseparateByV=false
-OseparateByJ=true
-OseparateByC=false <SampleName>.vdjca <SampleName>.clna
```

The number of clonotypes with TRB, IGHG, or IGHM constant gene call with read count >1 were calculated using Microsoft Excel and plotted using GraphPad Prism.

*Target enrichment of spatial cDNA with hybridization capture (LR/SR-Spatial VDJ)*
For target enrichment we used IDT xGen Hybridization and Wash Kit (#1080584) with one Discovery pool (IDT) each for B cell receptor and T cell receptor transcripts (IG and TCR pool) (table S1). The B cell receptor (BCR) consists of two heavy chains (encoded by the *IGH* gene loci) and two light chains (*IGK* or *IGL*) (*37*). The T cell receptor (TCR) is either made up of an α (*TRA*) and β chain (*TRB*), or a γ (*TRG*) and δ (*TRD*) chain (*38*), with T cells expressing TCRs encoded by TRA and TRB (αβTcells) being more abundant. Here, we focus our spatial analysis on αβT cells; i.e. probes targeting *TRD* and *TRG* were not included. Custom blocking oligos (IDT) (Table S1) were designed to hybridize to universal adaptor sequences of the cDNA library, to prevent off-target fragments from binding to BCR/TCR transcripts and contaminating the

enriched library. To ensure sufficient product for the hybridization capture protocol, we co-captured the TR and IG transcripts in the same reaction. The IG and TCR discovery pools were mixed at ratios 1:3 or 1:12.
We followed the protocol "xGen hybridization capture of DNA libraries", version 4 (IDT), with an input of 10 µl Visium cDNA per reaction, corresponding to between 45-130 ng, and with the hybridization reaction carried out overnight.
The enriched and purified libraries were amplified twice with an AMPure bead wash after each PCR, using 25 µl 2xKAPA mix, 7.5 µl cDNA primers (10X Genomics) and 17.5 µl sample in MQ water. The following settings were used for the PCRs:

1. 98°C: 3 min
2. 98°C: 15sec
3. 63°C: 30sec
4. 72°C: 2min
5. Repeat steps 2-5 6x for a total of 7 cycles ($1^{st}$ PCR) and 4x for a total of 5 cycles ($2^{nd}$ PCR)
6. 72°C: 1min

*Enrichment test*
To evaluate the performance of target enrichment with hybridization capture, we used 10 µl from Visium Gene Expression cDNA libraries (n=2) and performed enrichment as described above. The resulting samples were prepared for sequencing using the Visium Gene Expression protocol and, for comparison, we also prepared matched libraries from 10 µl of cDNA without enrichment (n=2). All four libraries were sequenced on the Illumina Nextseq2000 platform with a 28+150 setup and downsampled to 9M reads per sample. The data was analyzed with the Spaceranger pipeline, as described below.

*Long read library preparation and sequencing (LR-Spatial VDJ)*
The product from the hybridization capture was used as input into the SMRTbell library preparation (PacBio). We concentrated the DNA by AMPure Bead Purification (0.8x), eluting in 6 µl of EB buffer, using 1 µl for Qubit measurements. We used at least 1 µg of input for each library and multiplexed 8 samples in one sequencing run. We used PacBio Barcoded Overhand Adapters for multiplexing and followed the manufacturer's instructions for the library preparations. This protocol indexed the enriched libraries by ligation to avoid unnecessary PCR cycles, which can introduce errors and chimeric reads that are particularly problematic for long-read sequencing. The pooled library had more than the minimum amount required for sequencing. A SMRT Enzyme clean-up kit was used to remove linear and single-stranded DNA. The final libraries were sequenced on a Sequel II at the National Genomics Infrastructure (NGI)/Uppsala Genome Center.

*Semi-nested PCR (SR-Spatial VDJ)*
After hybridization capture and post-capture PCR amplification (14 cycles), further TRA and TRB enrichment was performed by PCR reactions with the following primers: V primers targeting either the TRAV or TRBV genes, 5' of the CDR3 region (i.e. 'Outer' TRAV or TRBV primers) and a primer ('partRead1') targeting the universal partial read 1 sequence present on all poly-dT captured transcripts in Visium Gene Expression cDNA

libraries (see table S1 for primer sequences). PartRead1 is also compatible with TruSeq indexes to allow sample multiplexing for sequencing. The Outer V primer PCR input was 2,5-5ng of hybridization captured cDNA from tonsil Visium Gene Expression libraries and the reaction was run with KAPA HiFi HotStart ReadyMix (2X) (KAPA Biosystems). Four replicate reactions were prepared per sample and pooled at the final indexing step. All primers were diluted 40x for a final concentration of 2.5 µM from 100 µM stocks (Integrated DNA Technologies). The PCR was run for 12 cycles for TRB and 14 cycles for TRA, under the following conditions:

1. 98°C: 5 min
2. 98°C: 20sec
3. 65°C: 30sec
4. 72°C: 1:30min
5. Repeat steps 2-5 11 or 13x for a total of 12 or 14 cycles
6. 72°C: 7min

Quantitative real-time PCR (qPCR) was performed to determine the appropriate number of cycles (to avoid exponential amplification). The Outer V primer PCR product was purified using a magnetic bead clean-up with AMPure beads (0.6x), followed by two 80% EtOH washes. The Outer V primer PCR product was eluted in EB buffer after incubation at 15min at 37°C. The cleaned PCR product was quantified using Qubit and BioAnalyzer (Agilent). 5 ng of each PCR product was used as input to the subsequent Inner V primer PCR.

The Inner V primer PCR was performed with the following primers: V primers targeting either the TRAV or TRBV genes, close/adjacent to the CDR3 region (i.e. 'Inner' V primers) and the same universal partial read 1 primer as described for the Outer V primer PCR ('partRead1') (see table S1 for primer sequences). These Inner V primers have a handle compatible with TruSeq indexing (table S1). The primer concentrations and reagents were as described for the OUTER V primer PCR. Quantitative real-time PCR (qPCR) was performed again to determine the appropriate number of cycles (to avoid exponential amplification). The following conditions were used for the PCR reaction:

1. 98°C: 5 min
2. 98°C: 20sec
3. 72°C: 30sec
4. 72°C: 1:30min
5. Repeat steps 2-5 7x for a total of 8 cycles
6. 72°C: 7min

The PCR product was cleaned using AMPure magnetic bead-clean up and ethanol washes as described above. The final eluted PCR products were quantified using Qubit and BioAnalyzer (Agilent). For the indexing step, 10 ng of input material was used from each replicate. All replicates for each individual sample were pooled at this stage and received the same index. The samples were PCR indexed using TruSeq Indexes, according to the conditions listed below and as described previously (*4*).

1. 98°C: 3 min
2. 98°C: 20sec
3. 60°C: 30sec
4. 72°C: 1:30min
5. Repeat steps 2-5 7x for a total of 8 cycles
6. 72°C: 5min

The PCR product was cleaned using AMPure magnetic bead-clean up and ethanol washes as described above. The final eluted PCR products were quantified using Qubit and BioAnalyzer (Agilent) and pooled for sequencing. , The samples were sequenced on a Novaseq6000 SP flowcell using a short read 1 and a longer read 2 to capture the entire CDR3 region and part of the constant region from the 5' end (35- 6- 259 setup).

*RNAscope*
Fresh frozen tissue sections were fixed for one hour in pre-chilled 4% PFA, the rest of the procedure was done according to the RNAscope® Multiplex Fluorescent Reagent Kit v2 Assay User Manual (Document Number 323100-USM). For tonsil section #25 (fig. S3A), probes targeting SDC1 (Cat#: 416961-C2; lot#: 22076C) and MS4A1 (Cat#: 426771-C3; lot#: 22076C) were used. For tonsil section #26 (fig. S3A), probes targeting CD3E (Cat#: 553971-C2; lot# 22076D) and MS4A1 were used. Control tonsil sections were incubated with either the RNAscope® 3-plex Positive Control Probe_Hs (320861; lot# 21258A) or the RNAscope® 3-plex Negative Control Probe_Hs (320871; lot# 21285A).

The TSA Vivid Fluorophore kit 570 (7526/1 KIT; batch 1A) was assigned to channel 2 with a concentration of 1:1500 for tonsil section #25 and concentration 1:750 for tonsil section #26. The TSA Vivid Fluorophore kit 650 (7527/1 KIT; batch 1A) was assigned to channel 3 with a concentration of 1:750 for both tonsil sections. For the control tonsil sections the TSA Vivid Fluorophores were assigned to the channels according to the RNAscope® Multiplex Fluorescent Reagent Kit v2 Assay User Manual. The following reagents were also used:
RNAscope® multiplex fluorescent detection reagents v2 (Cat#: 323110; lot# 2014906),
RNAscope® hydrogen peroxide (Cat#322335; lot#2015184), and
RNAscope® protease IV (Cat#322336; lot#: 2015288).

Images were acquired with Zeiss LSM700 confocal microscope with Zen2012 software and tile scan stitching was performed with Zen 2012 software and image processing was performed with Image J/Fiji software (version 1.53 for Mac, Java 1.8.0_172).

*Bulk SS3-based VDJ library preparation (Bulk SS3 VDJ)*
We modified the recently described Smart-seq3 (SS3) method to accommodate the use of low concentrations (10-100 ng) of purified RNA as the input and the use of many replicates to reduce amplification biases. This method will amplify UMI-barcoded IG and TR cDNA using primers directed to constant regions and a universal sequence 5' of the UMI region, similar to the 10x Genomics VDJ 5' protocol. We first isolated total RNA from fresh-frozen tonsil tissue. A section each of 10 µm thickness was placed in a Lysing Matrix D (MP Biomedicals) tube and lysed in a FastPrep (MP Biomedicals). Total RNA was extracted using the RNeasy Plus Mini Kit (Qiagen) following the manufacturer's protocol and the concentration of each sample was determined

before storing at -80C until needed for downstream cDNA generation and amplification. For cDNA generation, we added 1.2 µl of RNA (15-40ng/sample) to 1.6 µl of lysis buffer mixture (0.4 µl Poly-ethylene Glycol 8000 (50% solution), 1 µl H2O, 0.1ul dNTP (10mM, Thermofisher), 0.04 µl oligo dT30VN (100uM solution, IDT), 0.03 µl Triton X-100 (10% solution) and 0.03 µl RNAse Inhibitor (TaKaRa) in 96 well V-bottom plate (ThermFisher) placed on a cool rack kept on ice to prevent RNA degradation. The samples were then incubated at 72$^{o}$C for 10 minutes on a thermocycler and quickly placed back on the cool rack. We next added 1 ul of reverse transcription buffer mix to each sample (0.1 µl Tris-HCl pH 8.3 (1M), 0.12 µl NaCl (1M), 0.1 µl MgCl2 (100mM), 0.04 µl GTP (100mM), 0.32 µl DTT (100mM), 0.05 µl RNAse Inhibitor (40 U/µl RRI, TaKaRa), 0.08 µl TSO oligo (100 µM, IDT - /5Biosg/AGAGACAGATTGCGCAATGNNNNNNNNrGrGrG), 0.04 µl Maxima H-minus RT enzyme (200 U/µl), 0.15 µl H2O) and ran them on a thermocycler with the following conditions:

1. 42°C: 90 min
2. 50°C: 2 min
3. 42°C: 2 min
4. Repeat steps 2-3 9x for a total of 10 cycles
5. 85°C: 5 min
6. 4°C: hold

After reverse transcription, 6 µl of a PCR master mix (2 µl Kapa HiFi Hotstart buffer (5x, Roche), dNTPs 0.12 µl (10 mM, Thermo Fisher), MgCl2 (100 mM), 0.05 µl SS3 standard Forward Primer (100 µM, IDT - 5'TCGTCGGCAGCGTCAGATGTGTATAAGAGACAGATTGCGCAA*T*G3'), 0.01 µl Rev Primer (100uM, IDT – 5'ACGAGCATCAGCAGCATAC*G*A), 0.2 µl DNA Polymerase (1 U/µl, Roche), 3.57 µl $H_2O$) was added to each well and the samples were run on a PCR machine with the following conditions:

1. 98°C: 3 min
2. 98°C: 20 sec
3. 65°C: 30 sec
4. 72°C: 4 min
5. Repeat steps 2-4 11x for a total of 12 cycles
6. 72°C: 5 min
7. 4°C: hold

The amplified cDNA was purified using a magnetic bead clean-up with AMPure beads (0.6x), followed by two 80% EtOH washes to remove primers. Samples were resuspended in 15 µl of nuclease-free H2O and cDNA concentration was determined by qBit (DNA High Sensitivity kit) and checked for high-quality cDNA by Bioanalyzer (DNA High Sensitivity kit, Agilent).

To amplify TCR or BCR amplicons the samples were next split into two reactions as described below:

1) for TCR amplification we took 3 µl of cDNA and amplified it with PCR mix (12.5 µl KAPA 2x HiFi master mix (Roche), 1 µl Forward primer (10 µM), 1 µl Reverse primer (10 µM), 7.5 µl $H_2O$). The forward primer (SS3 short forward primer: TCGTCGGCAGCGTCAGATGTGTATAAGAGACAG) targets the universal sequence 5' of UMIs added in the Smart-seq3 reverse transcription step and an equimolar reverse primer mix containing TCRa or TCRb-specific primers targeting the constant region of either TCR transcript 200-300bp downstream of the exon start sites (table S1 for the primer sequences).
2) For BCR amplification the same process was done but instead of TCR constant primers, we used an equimolar mix of 7 primers targeting the different heavy (IGHA, IHGD, IGHE, IGHG, IGHM) and light (IGLC, IGKC) chains constant regions 200-300bp downstream of exon start sites (table S1 for the primer sequences).

PCR conditions for both reactions were as follows:

1. 98°C: 3 min
2. 98°C: 20 sec
3. 68°C: 1 min 15 sec
4. Repeat steps 2-3 11x for a total of 12 cycles
5. 4°C: hold

Amplified TCR and BCR cDNA libraries were again cleaned with AMPure XP beads as before to remove primers and resuspended in 10 µl of nuclease-free H2O. Sample concentrations were determined by QuBit (ThermoFischer Scientific).

A second PCR was next performed for both TCR and BCR amplicons to shorten the 3' end of each amplicon for illumina sequencing of the internal V-D-J recombinant sequence (CDR3).

1) For TCR amplicons 3 µl of amplified cDNA from PCR1 was added to a new PCR mixture (12.5 µl KAPA 2x HiFi master mix (Roche), 1 µl Forward primer (10 µM), 1ul Reverse primer (10 µM), 7.5 µl $H_2O$) containing a longer universal forward primer (SS3 Forward primer: TCGTCGGCAGCGTCAGATGTGTATAAGAGACAGATTGCGCAA*T*G) and an equimolar reverse primer mix containing TCRa or TCRb-specific primers targeting the constant region of either TCR transcript downstream of the exon start sites (table S1 for the primer sequences). Both the forward and reverse primers contained sequences to allow for direct ligation of i5 (5') or i7 (3') Nextera XT indexes for Illumina sequencing.
2) For BCR amplicons the same process was performed with the use of BCR-specific constant primers instead of TCR primers for the reverse primer set (table S1 for the primer sequences).

PCR conditions are listed below:

1. 98°C: 3 min
2. 98°C: 20 sec
3. 68°C: 1 min 15 sec

4a. Repeat steps 2-3 x19 for a total of 20 cycles for TCR amplicons
4b. Repeat steps 2-3 x11 for a total of 12 cycles for BCR amplicons
5. 72°C: 5 min
6. 4°C: hold

Amplified TCR and BCR cDNA was size-selected by a 2-step bead cleaning (SPRI beads) to remove primers and any larger sequences that appeared in the reactions and resuspended in 20 µl of nuclease-free H2O. Sample concentrations were determined by QuBit and purity of amplicons was assessed by bioanalyzer (DNA High Sensitivity, Agilent).

For sequencing individual reactions were indexed using Nextera XT primers (Illumina, 96 primer kit) which were added by PCR using the following PCR mix (2 µl cDNA (approx. 100 ng), 5 µl KAPA 5x buffer (KAPA HiFi Hotstart, Roche), 1.5 µl dNTPs (10 mM, Roche), 1 µl Taq Polymerase (KAPA HiFi Hotstart kit, Roche), 1 µl i7 primer (10 µM, Nextera XT, Illumina), 1 µl i5 primer (10 µM Nextera XT, Illumina), 11.5 µl $H_2O$).

PCR conditions are listed below:

1. 72°C: 3mins
2. 95°C: 30 sec
3. 95°C: 10 sec
4. 55°C: 30 sec
5. 72°C: 1 min
6. Repeat steps 3-5 x7 for a total of 8 cycles
7. 72°C: 5 min
8. 4°C: forever

Final libraries were cleaned with AMPure XP beads and resuspended in $H_2O$. Individual samples were measured and mixed in equimolar ratios for sequencing on the Illumina NovaSeq 6000 platform (2x 150 set-up) by the National Genomics Infrastructure, SciLifeLab (Solna).

*Cell processing for single-cell RNA sequencing*
Single-cell suspensions from separate breast tumor regions (P1: n=5; P2: n=4) were prepared by enzymatic tissue dissociation using the human Tumor Dissociation Kit (Miltenyi Biotec, 130-095-929) and gentleMACS dissociator (Miltenyi Biotec). Cell suspensions were stained with the Zombie Aqua Fixable viability dye (Biolegend, 423101) at room temperature for 20 minutes, then washed with Phosphate Buffered Saline (PBS). The cells were incubated with Human TruStain Fc block (Biolegend, 422302) for 10 minutes to limit nonspecific antibody binding, then stained for 20 minutes with anti-CD45 (1:40, Biolegend, 304021), anti-EPCAM (1:40, Biolegend, 324206) and cell hashing TotalSeq-C antibodies (Biolegend, 394661, 394663, 394665, 394667, 394669) in Fluorescence-activated cell sorting (FACS) buffer (PBS + 0.5% Bovine Serum Albumin). Each tumor region was labeled with a unique TotalSeq-C antibody to

enable pooling all tumor regions into a single 10x sample. The cells were subsequently washed and resuspended in FACS buffer. FACS was performed using an influx flow cytometer (BD Biosciences) to sort live $EPCAM^{-}CD45^{+}$ single cells for 10x Genomics Chromium Single Cell gene expression analysis. At least 25-30,000 cells were sorted per tumor region and all tumor regions for each patient were sorted into the same tube. Single stain controls (cells and beads) and fluorescence minus one controls (FMO), containing all the fluorochromes in the panel except the one being measured, were used to set voltages and to define the proper gating strategy.

*10x Genomics Chromium Single Cell library preparation and sequencing*

Single cell gene expression (GEX), VDJ clonotype, and feature barcoding libraries were generated from $EPCAM^{-}CD45^{+}$ cells using the 10x Genomics Chromium Single Cell 5′ assay, following the manufacturer's instructions. Libraries were profiled and quantified using a Bioanalyzer High Sensitivity DNA kit (Agilent Technologies) and Qubit High sensitivity kit (ThermoFischer Scientific). Final single cell gene expression libraries were sequenced (30,000 reads/cell for GEX; 4-6000 reads/cell for VDJ clonotype, and 4-5000 reads/cell for feature barcode libraries) on a NovaSeq 6000 SP flowcell (Illumina, 150-8-8-150 read set-up) by the National Genomics Infrastructure, SciLifeLab.

*Data processing of Spatial Gene Expression Libraries (Spatial GEX)*

Following demultiplexing of bcl files, read 2 fastq files were trimmed using Cutadapt (*39*) to remove full-length or truncated template switch oligo (TSO) sequences from the 5' end (beginning of read 2) and polyA homopolymers from the 3' end (end of read 2). The TSO sequence (AAGCAGTGGTATCAACGCAGAGTACATGGG) was used as a non-internal 5' adapter with a minimum overlap of 5, meaning that partial matches (up to 5 base pairs) or intact TSO sequences were removed from the 5' end. The error tolerance was set to 0.1 for the TSO trimming to allow for a maximum of 3 errors. For the 3' end homopolymer trimming, a sequence of 10 As was used as a regular 3' adapter to remove potential polyA tail products regardless of its position in the read, also with a minimum overlap of 5 base pairs. The trimmed data were processed with the spaceranger pipeline (10X Genomics), version 1.2.1 (tonsil) and version 1.0.0 (BC) and mapped to the GRCh38 v93 genome assembly.

*Long read sequencing analysis (LR-Spatial VDJ)*

The input to the analysis was demultiplexed consensus reads from the long-read Pacbio sequencing (github: HoseinT/long-read-processing-STAR-paper). The analysis was done using Python programming language. First, the fastq files were parsed into a dataframe with readID, sequence, and quality columns. Second, we searched for the Truseq adapter and the TSO sequence to anchor the ends of each read and discarded the reads that lacked these sequences. Third, we searched for the pattern 'CGACGCTCTTCCGATCT' which is part of the Truseq adapter starting in the first seven bases of either the read or its reverse complement. If any of the positions matched the sequence with hamming distance 1 or less we marked it. We did the same for the TSO sequence ('TCTGCGTTGATACCACT'). We reverse complemented the reads as needed so that all the reads have the Truseq adapter at the beginning and the TSO at the end. We identified the spatial barcode and the UMI by obtaining the first 16 bases following the Truseq adapter for the spatial barcode and the 12 bases following that as the UMI. We made sure that the following 4 bases were all T's and filtered out the reads that had any other bases in that interval. We noticed that the most common UMI was a polydT, which was considered an artifact, and those reads were removed. We also defined the end of polyT region as the first matching position for the pattern '[^T]T{0,2}[^T]T{0,2}[^T]' after 28 bases after the end of the Truseq adapter location.

*LR-Spatial VDJ clonotype analysis*
To run MIXCR (version 3.0.3), we trimmed the polyT and TSO and wrote the reads to a new fastq file. We used MIXCR with this command:

'mixcr analyze shotgun -s hsa --align -OsaveOriginalReads=true --starting-material rna <TrimmedFastq> <SampleName>'

then ran another MIXCR command to report alignments for each read:

'mixcr exportAlignments -f -cloneIdWithMappingType -cloneId -readIds -descrsR1 <SampleName>.clna <ReportFile>'

Then, we used the resulting tabular file to assign the reads to the clonotypes in MIXCR output. To use UMIs, we filtered out reads that did not map to any clone (cloneID==-1), then grouped the reads table by the Visium barcode and UMI and counted how many reads they have and how many clones were associated with each UMI. We filtered out UMIs that have been assigned to more than one clonotype, since they were likely due to PCR or sequencing errors. A count matrix (Clone ID x spatial barcode) was generated and merged to the clone list exported by MiXCR. After removing spots outside the tissue section (as is standard for the spaceranger pipeline), we imported the gene expression cluster and follicle annotations and calculated UMI, spatial barcode and overlaps among different clusters, follicles, and sections.

*SR-Spatial VDJ clonotype analysis*
To preprocess short-read sequences, we modified the example workflow of "UMI Barcoded Illumina MiSeq 325+275 paired-end 5'RACE BCR mRNA" from pRESTO (version 0.6.2, Immcantation group)(*40*). Reads with less than 20 Phred quality scores were regarded as low-quality sequences and were removed. Then, we removed reads that lacked a valid spatial barcode by comparing the reads to the Visium barcode list. We only kept the reads that included the 'inner V' primer sequences used in the second step of the TRA and TRB semi-nested PCR enrichment steps. Input fastq files that include more than 13 millions will be downsampled to 13 million reads by SplitSeq.py samplepair, due to the platfrom limitation (downsample rate: 0.76-0.94). To have a more accurate UMI collapsing result, we regarded both the UMI and spatial barcode sequence as the "UMI" region in the pipeline. We selected UMIs with at least 3 reads per group in tonsil for repertoire analysis.

We merged all reads from different tonsil sections and used MiXCR (version 3.0.13) with the following commands:

‘mixcr analyze amplicon --library repseqio.v1.5 --species hsa --starting-material rna --5-end no-v-primers --3-end c-primers --adapters adapters-present --align "-OsaveOriginalReads=true" --assemble "--write-alignments" <R1Fastq> <R2Fastq> <SampleName>’

To report the alignments for each read, we further ran the MIXCR command:

‘mixcr exportAlignments -cloneId -descrsR2 -targetSequences -nFeature CDR3 -jGene <SampleName.clns> <SampleName.tsv>’

Then, we used customized R scripts to create the clonal UMI count matrix for each tonsil section. The exported UMI collapsed reads that lacked a valid CDR3 call from MiXCR output, did not map to any clone (cloneIdID == -1), or had fewer read counts than 3 reads per UMI were removed. We observed some cases of index hopping, i.e. where individual UMI collapsed reads from different samples contained identical nucleotide CDR3 sequence, J gene, spatial barcode, and UMI sequence. The clone that contained the most read count in its UMI group was considered the true UMI collapsed reads and the rest discarded. At these settings, we still observed a small number (1-4 per section) of TR clones with an unexpectedly high spatial barcode number per clone (that was not reproduced in the LR-Spatial VDJ data), suggestive of PCR over-amplification and artifacts in spatial barcode region. Therefore, we removed those clones from further analysis. Section overlap was calculated the same way as we did for LR reads.

**Visualization and Analysis of Spatial data**

*Gene expression data of tonsil #1 and #2*

Two sets of tissue images, spatial coordinates, and count matrices generated through the Spaceranger (10x Genomics) pipeline were imported into R (R Core Team, 2017) using the InputFromTable() function of the STutility package (*41*). For fig. S1C, when applicable, features were collapsed based on antigen receptor chain identity and visualized using the FeatureOverlay() function.

*Enrichment test*

Spaceranger output data were imported into R as described for tonsil #1 and #2. Four datasets were loaded; one enriched and one non-enriched per original cDNA library (tissue section). A Seurat object (*42*) was created to contain all four datasets, using the InputFromTable() function of the STutility package (*41*). No filtering of genes was performed, but spots outside the tissue were removed. Gene groups were aggregated into supergroups for visualization in bar plots and on tissue (fig. S2), including only genes belonging to constant segments. For example, counts of IGHA1 and IGHA2 were summed in the supergroup IGHA, and TRBC1 and TRBC2 into TRBC. For statistical analysis fig. S2, we collapsed all counts from any constant chain of IGH, IGK, and IGL into the supergroup IG and all TRA and TRB into TR. Next, we performed a statistical test using the compare_means() function of the ggpubr package, method = wilcox.test. In these tests, we compared all data points of the two enriched samples to those of the two non-enriched for aggregated IG and TR counts, respectively.

*Spatial GEX and LR/SR-Spatial VDJ tonsil and tumor data*
Spaceranger output data were imported into R as described for tonsil #1 and #2. One Seurat object (*42*) per sample type or patient (tonsil, P1 and P2) was created using the STutility package. For each dataset, genes were filtered for a minimum of 100 counts across the entire dataset and a minimum of 5 spots. Spots were also filtered based on the number of counts; spots with at least 300 counts (P1) or 500 (tonsil and P2) were kept.

In order to add clonotype data to the Seurat objects described above, the clonotype count matrices were modified to contain the same barcodes (bcs) as the gene expression count matrices. Accordingly, bcs absent in the gene expression count matrix (after filtering described above) were removed, and bcs only present in the gene expression count matrix were added to the clonotype count matrix and filled with zeros for each clone. Finally, clonotype names were edited to include the first four letters from the "C hit" column of the MiXCR output, followed by "clone" and the number assigned by MiXCR. Each modified clonotype count matrix was then loaded as a new assay into its respective Seurat object (tonsil, P1, and P2), whereafter genes and clonotypes were visualized on the tissue images using built-in functions of the STUtility package.

*Normalization and Clustering of GEX data*
The gene expression data of tonsil, P1, and P2, respectively, were normalized across spots with Seurat SCTransform() function. Dimensionality reduction was performed with non-negative matrix factorization, using STUtility RunNMF() function with 24 factors. These factors were subsequently used for clustering using Seurat FindNeighbours() function followed by Seurat FindClusters() at resolution 0.8.

For Tonsil #2, this strategy rendered 13 clusters, which were annotated and grouped based on shared gene expression and single cell deconvolution results from *stereoscope* (method described below). The cluster groups were consistent with known anatomical structures observed in H&E stains of each section. For follicle annotation on tonsil sections, each follicle/GC had to contain at least two spots belonging to at least one of the four clusters annotated as B cell follicle/GC cluster group (c1, 4, 5, or 7) and be found in at least two independent sections.
For P1 and P2 breast tumor samples, the strategy described above generated 18 and 12 clusters, respectively. Tumor areas were annotated based on gene expression cluster and single-cell deconvolution results per Spatial GEX cluster from *stereoscope* (method described below). All tumor clusters had enriched tumor-associated gene expression and cancer epithelial cell signatures. Cluster 13 (P1) exhibited a heterogeneous epithelial signal, with cancer and normal epithelial cell enrichment varying across sample regions. Due to the presence of cancer epithelial signature in at least one sampled region, it was denoted as 'Tumor', but likely contains normal breast epithelial tissue as well. Based on the breast cancer spatial cluster annotation and border detection with STUtility RegionNeighbours() function, we adapted Seurat FindMarkers() function for calculating differentially expressed LR- Spatial VDJ IG clones across regions. Specifically, we used the Poisson test, with a minimum of 0.2-fold difference and 0.01 as the minimum percentage of spots in an area expressing a clonotype. The latter parameter is drastically reduced from the default of 0.1, which is optimized for scRNA-seq analysis. We reasoned that spatial VDJ data should require a lower value to reveal clonotypes that are region-specific but only expressed in a minority of spots in that region.

*Bulk SS3 Clonotype analysis*
To preprocess bulk SS3 sequences, we modified the same example workflow used in short-read sequence analysis. We trimmed the first 11 bases in R1 to remove the ATTGCGCAATG sequence pattern by cutadapt (*39*). Reads with less than 20 Phred quality scores were removed due to low-quality. We corrected insufficient UMI diversity and removed those reads that lacked the primer sequences used in the bulk SS3 VDJ enrichment steps. We collapsed reads into UMI groups, which were used for repertoire analysis.

We merged all reads from different tonsil sections and ran MiXCR (version 3.0.13) on both TCR and BCR reads with these commands:

mixcr analyze amplicon --library repseqio.v1.5 --species hsa --starting-material rna --5-end no-v-primers --3-end c-primers --adapters adapters-present --align "-OsaveOriginalReads=true" --assemble "--write-alignments" <R1Fastq> <R2Fastq> <SampleName>

To report alignments and clones, we further ran the MIXCR command:
mixcr exportAlignments $<SampleName>.clns
Tonsil_$<SampleName>_alignment_customized_hit_des.tsv -readIds -descrsR1 -descrsR2 -cloneId -vHit -dHit -jHit -cHit -vAlignment -dAlignment -jAlignment -cAlignment -targetSequences -targetQualities -chains -targets -defaultAnchorPoints

Post MiXCR analysis was finished in R with customized code. Section overlap was calculated the same way as we did for LR reads.

*Single cell deconvolution (Stereoscope) Analysis*
We used the software *stereoscope* (v.0.3) to decompose the spatial gene expression profiles into contributions from specific cell types, a.k.a. single cell mapping (*12*). *stereoscope* uses a probabilistic framework that models both single cell and spatial transcriptomics data with a negative binomial distribution. In short, *stereoscope* first learns cell type and gene specific parameters from the single cell data (where no mixing of cells occur), to then use these parameters in a guided decomposition of the mixed gene expression profiles in the spatial transcriptomics data.

The input to *stereoscope* is the raw (unnormalized) single cell UMI count data, cell type annotations, and raw (unnormalized) spatial transcriptomics UMI count data. The output from *stereoscope* is a [spot]x[cell type] matrix, where each element gives the proportion of cells at a given spot that belongs to a specific cell type. For more details we refer to the original *stereoscope* publication. Below, we outline the two analyses that *stereoscope* was used for.

**Tonsil analysis**
In the tonsil analysis, we used the tonsil single cell data (*13*) as well as the tonsil Visium spatial transcriptomics data presented in this paper. The tonsil data had two different annotation levels, Lineage and Cellsubset, we analyzed both these tiers in the exact same manner, which is described below.

For computational efficiency, we subsampled the single cell data according to the following strategy: if a cell type had fewer than 3 cells, exclude the cell type; if a cell type had more than 3 cells but fewer than or exactly 1000 cells, use all cells; if a cell type had more than 1000 cells, randomly pick (without replacement) 1000 cells from the population. We did not use all of the genes, but instead relied on a set of highly variable genes (based on the single cell data), which were extracted according to the following recipe (using the *scanpy* suite, v.1.8.2) (*43*):

```
sc.pp.normalize_total(...,1e4)
sc.pp.log1p(...)
sc.pp.highly_variable_genes(...,n_top_genes=5000)
```

where the first argument represents the object holding the single cell data. Using the set of highly variable genes we ran *stereoscope* with 50000 epochs and a batch size of 2048, all other parameters were set to the default - the number of epochs and batch size were identical for both steps in *stereoscope*.

**Breast cancer analysis**
In our breast cancer analysis we used an external single cell data set by Wu et al. (*30*), while the Visium spatial transcriptomics data set was produced in-house (presented in this paper). The single cell data set contains three different tiers of "granularity", i.e., major, minor, and subset, listed from least to most fine annotation level. For this study, we focused on the minor and subset tiers. These were both analyzed using the exact same approach, which is given below. For P1 Region D, only samples D1 and D2 were included in the analysis.

We employed the same subsampling strategy as described for the tonsil data (see previous paragraph). A select set of genes was used in this analysis as well. This gene set was taken from a previous study where the same single cell data set was mapped to a different spatial transcriptomics data set; more precisely, it is the gene set given in Supplementary Data 13 in Andersson et al (*18*). We refer to the cited reference for more details about this gene set but, in brief, this gene set consists of the most expressed genes (in the single cell data) together with certain marker genes associated with the cell types present in the single cell data set. As in the tonsil analysis, 50000 epochs and a batch size of 2048 were used in the analysis, with all other parameters set to their default values.

*Single-cell gene expression and VDJ data processing*
Sequencing outputs were processed by Cell Ranger (version 5.0, 10X Genomics) (*44*). Gene-barcode count matrices were analyzed with the Seurat package (version 4.0, Satija Lab). Two steps of filtering were introduced here. First, raw gene expression matrices were subset by the barcode list in VDJ output, including T cell subsets and B cell subsets. Based on the UMI count, gene count, and mitochondrial percentage of raw gene expression matrices and their subsets, each threshold was selected to keep the maximum count of high-quality cells and avoid losing T and B cells which have VDJ sequencing outputs. Second, doublets in each sample were detected and filtered out by HTODemux() function in Seurat. All samples were integrated and scaled into one count matrix by Seurat. Dimension reduction, UMAP generation, and clustering, were performed on the merged dataset by Seurat. The merged dataset was clustered by a gradient of the resolution, from 0.2 to 2. We chose 0.8 as the final resolution by comparing the top-listed differentially

expressed genes in each cluster. Cell types were annotated by differentially expressed genes and their marker genes expression level. Our scRNAseq dataset also lacked neutrophils and eosinophils, which is a common occurrence in single cell RNAseq analysis and may be due to cell loss during tissue digestion (of note, neutrophils typically have a lower per cell UMI count, but we did not observe neutrophils even when lowering the UMI threshold during analysis). To further account for doublets after the first round of annotation, we removed T/NK cells that had B cell receptors, B cells that had T cell receptors, and cells from non-lymphocyte clusters that had either T or B cell receptors in the single-cell dataset. We also removed potential doublets from T and B cell VDJ output by excluding cells with more than two pairs of chain. After quality filtering, only cells that met either one of these criteria were kept: only one TRB chain, one TRA and one TRB chain, or one TRB chain and two TRA chains. Then, we reanalyzed the single-cell dataset and created new UMAP coordinates, differential gene expression list, and cluster/subcluster annotation. We used the following parameters for generaing breast cancer single-cell UMAP and cluster: runUMAP: dims = 1:15, n.neighbors = 25,min.dist = 0.2; findNeighbors: dims = 1:30; FindClusters: resolution = 0.8. For T cell subclustering, we used following parameters: runUMAP: dims = 1:10, n.neighbors = 30,min.dist = 0.2; FindNeighbors: dims = 1:10; FindClusters: resolution = 0.3. For additional VDJ clonotype quality filtering and to calculate the clonal overlap with the Spatial VDJ output (defined by MiXCR for ), cells that belonged to different clones from cellranger VDJ output but with identical nucleotide CDR3 sequence and J gene in both receptors (i.e. TRA/TRB in T cells, IG light chain/IG heavy chain in B cell, etc.) would be merged and assigned with a renamed clone ID by a customized R script, all cells within the new clone were assigned the minimum clone ID within the group. The T cells that had a single TRB sequence, which CellRanger had identified as separate clones, were merged to other clones expressing the matched TRB nucleotide CDR3 and J gene. We kept unmatched single chain TRB clones in the clonal list. In addition, we removed doublet "TGCTACCGTTCAGGCC-4" by comparing its receptor chain to other cells in the same clone. All the dimension reduction and annotation results, along with the VDJ output files were imported into Loupe Browser (version 5.0, 10X Genomics) and Loupe VDJ Browser (version 4.0, 10X Genomics) for interactive analysis.

*Clonotype overlap analysis*

To compare clonal lists from the MiXCR analysis result, including LR-Spatial, SR-Spatial and Bulk SS3 VDJ, we used customized R script and marked clones with identical nucleotide CDR3 sequence and J gene as the overlapped clones, which was consistent with the default components that MiXCR used for collapsing reads into clone. If multiple clones had identical nucleotide CDR3 sequence and J gene, we grouped them together as one overlapped case. To compare clonal lists from Cell Ranger VDJ (single-cell) and MiXCR (LR- and SR- Spatial VDJ) outputs, we compared the single-cell clone list with MiXCR output by matched nucleotide CDR3 sequence and J gene. We then calculated the percentage of LR-Spatial VDJ clones that overlapped with other analysis methods. Clone fraction was calculated based on dividing each clonotype's UMI count with the total TR (*TRA, TRB*) or IG (*IGH, IGK, IGL*) UMI count from either LR- Spatial VDJ, SR- Spatial VDJ, or bulk datasets.

*Clonal evolution analysis*

**Pre-processing steps**

For this analysis, we collapsed reads within the same UMI group to generate a consensus read. Individual reads from the same sample with identical UMI and spatial barcode were grouped by their full-length sequence. One or more read groups with the most abundant read count were selected. Then, we calculated the average sequencing quality score for each read group. First, we kept the read groups with the highest quality score. Second, if multiple groups were present, we selected the group with the longest sequence. Third, for the remaining read groups with identical length, average quality score, and read count, we masked the different bases between the groups as N and generated the final sequence for that UMI group. Then, we extracted the section information from the R2 header (descrsR2 column) from the reads alignment output, grouped reads by sections or replicates, and converted them to a count matrix (clone ID =row name; section number = column ID). Post filtered LR- Spatial VDJ reads were split into different segments including V, D, J, etc., by using anchor points ("refpoints" column) per sequence from the MiXCR alignment output. We referred to the anchor point definition and the script named "mixcr2imgt.py" from the Immcantation group to split LR- Spatial VDJ reads (*40*, *45*). Then, we loaded the read list into R and converted the column name to the Adaptive Immune Receptor Repertoire (AIRR) compatible format (*46*). We restricted our analyses to the heavy chain (IGH), since it should provide the richest clonal structure (*47*). IGH clones defined by MIXCR have unique CDR3 sequences, which separates closely related IG clones that have undergone somatic hypermutation. To group the MIXCR-defined clones together to define 'IGH clonal families', we adapted the instruction named "Clustering sequences into clonal groups" from the Immcantation group. We calculated the minimum normalized hamming distance between the nucleotide junction sequence and selected 0.1, 0.1, and 0.08 as the threshold according to their distribution for tonsil, breast cancer P1, and breast cancer P2 LR IGH clonal families, respectively. Of note, the CDR3 sequence contains the conserved cysteine and tryptophan/phenylalanine residues on its 5' and 3' side according to the AIRR standards documentation. However, the MiXCR results already included those two residues in the CDR3 sequence. Therefore, we simply renamed the nucleotide MIXCR CDR3 sequence to "junction". Finally, we grouped the IGH clones into IGH clonal families by "DefineClones.py" script from SHazaM (version 1.1.1, Immcantation group) (*45*). We extracted the IGH reads, removed the ones without a valid C region annotation, and excluded one read that had an IGLC alignment. The final read list was used for class-switching and lineage analysis.

**IGH clone family analysis**

We imported the tonsil LR- Spatial VDJ IGH clone family read list and the STUtility object into the R environment, extracted gene expression cluster and follicle annotation and merged them into the read list. For follicle analysis, we determined the follicle count per clone family by counting unique follicle ID per clone. Then, we converted all follicles as "in" to define intra-follicular IGH clonal families. We grouped reads by follicle to calculate follicle stats including UMI count, spatial barcode count, clone family count and their ratio. To describe the expansion level within each follicle, we ranked the follicles for each clonal family based on the max UMI percentage, and extracted the value per follicle. We calculated species richness based on the *Chao1* estimator (*48*):

$$Chao1 = s + \frac{n_1^2}{2n_2}$$

Where $s$ was the total clone family count within the follicle, $n_1$ was the UMI count of top expanded clone family within the follicle, and $n_2$ denoted for the UMI count of the second top expanded clone family.

**Class-switching analysis**

We removed IGHGP and IGHD reads from the tonsil LR- Spatial VDJ IGH clone family read list since IGHGP is a pseudogene and IGHD was regarded as an alternative splicing result from IGHM. B cells that exclusively express IGHD as a result of a "true" class switching event are most often rare, but can occur in mucosal tissues, including the tonsil (*8*, *13*, *49*); however, we detected very few clones with IGHD as the dominant isotype. Therefore, we concluded that this individual had few, if any, true IGHD class-switched clones, and these were not investigated further for the purpose of class switching. We also removed IGH clonal families with less than two reads since they would not contain class-switching events. We grouped reads within each clone family by follicle, cluster, or spatial barcode, and extracted class-switched reads within the samples. Putative class switch recombination events (pCSR) were defined as two sequences with identical CDR3 V gene sequences (i.e. hamming distance = 0) with two different IGH constant chains/isotypes co-localizing in the same spatial barcode.

**Lineage relationship analysis**

To create lineage trees using SHazaM (Immcantation group) (*45*), we imported the tonsil LR-Spatial VDJ IGH clone family read list and removed the reads that lacked a valid CDR3 sequence or contained indel(s) in the V sequence compared to the reference. We recovered the reference sequence based on the mutation patterns from the "bestVAlignment" column and inserted IMGT gaps. Then, we removed the reads that did not have a valid junction length (not divisible by 3, which potentially represented a frame-shift mutation). To create the lineage tree and assign unique subclone IDs to each node, we followed the vignettes of "Lineage reconstruction" in Alakazam (version 1.2.0, Immcantation group) (*45*) with phylip (version 3.697, Joe Felsenstein) for creating lineage trees (*50*). Each node of the tree (hereafter referred to as 'subclone') represents a unique V sequence versus the others and the size of the node is proportional to the number of unique reads belonging to that node. We traversed the lineage tree through depth-first search to assign adjacent ID to closely related subclones. Then, we plotted lineage trees and exported count matrices for visualizing nodes within the tissue section. We also matched the spatial barcodes to their follicle and cluster identity. Finally, we generated all possible trees and created the clone family subclones vs spatial barcode count matrix for statistical analysis and spatially visualization.

*Visualizations of IGH clonal families and subclones on tissue*

Count matrices of IGH clonal families and subclones were imported into R and added to the tonsil object as new assays. To generate aligned x and y coordinates of all six tonsil sections, and enable plotting of all data jointly, we used the functions MaskImages() and AlignImages() of the STUtiliy package. Next, all non-zero observations of an IGH clonal family or subclone were plotted and colored by tissue section on a transparent background. The same IGH clonal family or subclone was then visualized on one of the sections using the STUtility FeatureOverlay() function. The joint data image was superimposed on the tissue image and aligned by matching shared spots in the two image types using Adobe Illustrator.

*Pairing IG receptor chains with repair*

For pairing receptor chains, we filtered the LR-Spatial VDJ count matrix to include all IGL and IGK clonotypes, but to simplify the pairing, only one IGH clonotype per IGH clonal family (the one with highest counts), since multiple IGH chains could pair with a single light chain due to somatic hypermutation. The resulting count matrix was used as input in *repair* (github: almaan/star-repair) – a method we developed for pairing of receptor chains – and different values of spot and pairing score cut-offs were used to find the threshold that would generate the highest number of correct pairs and minimum number of incorrect pairs. A predicted pair was assigned as correct only if the two chains were a verified pair in the ground truth scVDJ dataset (which contained only pairs where both chains overlapped in both the spatial and single cell VDJ datasets), and incorrect if any of the chains were paired with another chain in the ground truth dataset.

*The repair method*
As mentioned above we developed a method for unsupervised pairing of the receptor chains; we refer to this method as *repair* (github: almaan/star-repair). In this method, we consider the task of pairing the different chains as an optimization problem. Below we elaborate on the details of this problem:

If $A \in R^{NxC_a}$ is the normalized expression matrix for the chains of type $a$, and $B \in R^{NxC_b}$ is the equivalent matrix for chains of type $b$, then we want to find a matrix $M \in {R_+}^{C_a x C_b}$ and scalar $g \in R_+$ such that $A = gBM^T$. Here, $N$ represents the number of observations, the number of $a$ type chains and $b$ type chains are given by $C_a$ and $C_b$ respectively.

$M$ can be considered as a 'mapping matrix' between chains of type $b$ to chains of type $a$. In layman's terms, element *(i,j)* in $M$ tells us how much of chain $b_j$ contributes to chain $a_i$*'s* expression, once we've accounted for the difference in abundance between the different chain types (taken care of by $g$). The scaling factor is taken as a scalar, shared across all chains, an assumption that equates to assuming that the difference in abundance between chains of type $a$ and $b$ is not influenced by the chains' identities.

During optimization we do not immediately learn the parameters $M$ and $g$ but rather the matrix $Q$ and scalar $p$ which relates to the former accordingly:

$$M_{ij} = \frac{pow(c,Q_{ij})}{\sum_k \; pow(c,Q_{ik})} \text{ and } g = exp(p)$$

Where $pow(c,x) = c^x$.Hence, $M$ is the row-wise generalized softmax (if c = $e$, then this equals the standard softmax) of Q, meaning that $M_{ij} \in (0,1)$. There are two benefits to using the (generalized) softmax function: i) it allows us to interpret the element $M_{ij}$ as the *probability* that chain $b_j$ is paired with chain $a_i$, ii) the larger the value of the base $c$, the more focused the probability mass will be on a single value. The second feature is considered as a benefit since it favors sparse mapping from chains of type $b$ to chains of type $a$, i.e., one $b$ chain dominates the explanation of one $a$ chain's expression; notably, the same $b$ chain can map to multiple $a$ chains.

Thus, the final optimization problem becomes: $min_{g,M} \; || A - gBM^T||^2$, with $g$ and $M$ defined as above. To solve this we use stochastic optimization (Adams optimizer), implemented using the *jax* python package (v.0.3.1).

Finally, once an optimal solution to *M* is found, as a pruning step, we formulate an unbalanced linear assignment problem (LAP) where we negate this *M* and use the result as our cost matrix. This generates a bijective (one-to-one) pairing between a subset of all chain types of *a* and *b* (if $C_a > C_b$ each *b* chain will be paired to an *a* chain, but not every *a* chain will be paired with a *b* chain, and vice versa for $C_a < C_b$. If $C_a = C_b$, then each *a* chain and *b* chain will be assigned a partner).

*Statistical analysis and figure presentation*

Data was processed using R, Python, and Microsoft Excel. Statistical analysis was performed and figures were prepared using R, GraphPadPrism (version 9.4.01), and Keynote (version 11.2). The experimental outlines in Fig. 1A, fig. S3A, and Fig. 3A were prepared by graphic designer Mattias Karlén. Other illustrations were prepared in Keynote or Adobe Illustrator by the authors.

### fig. S1

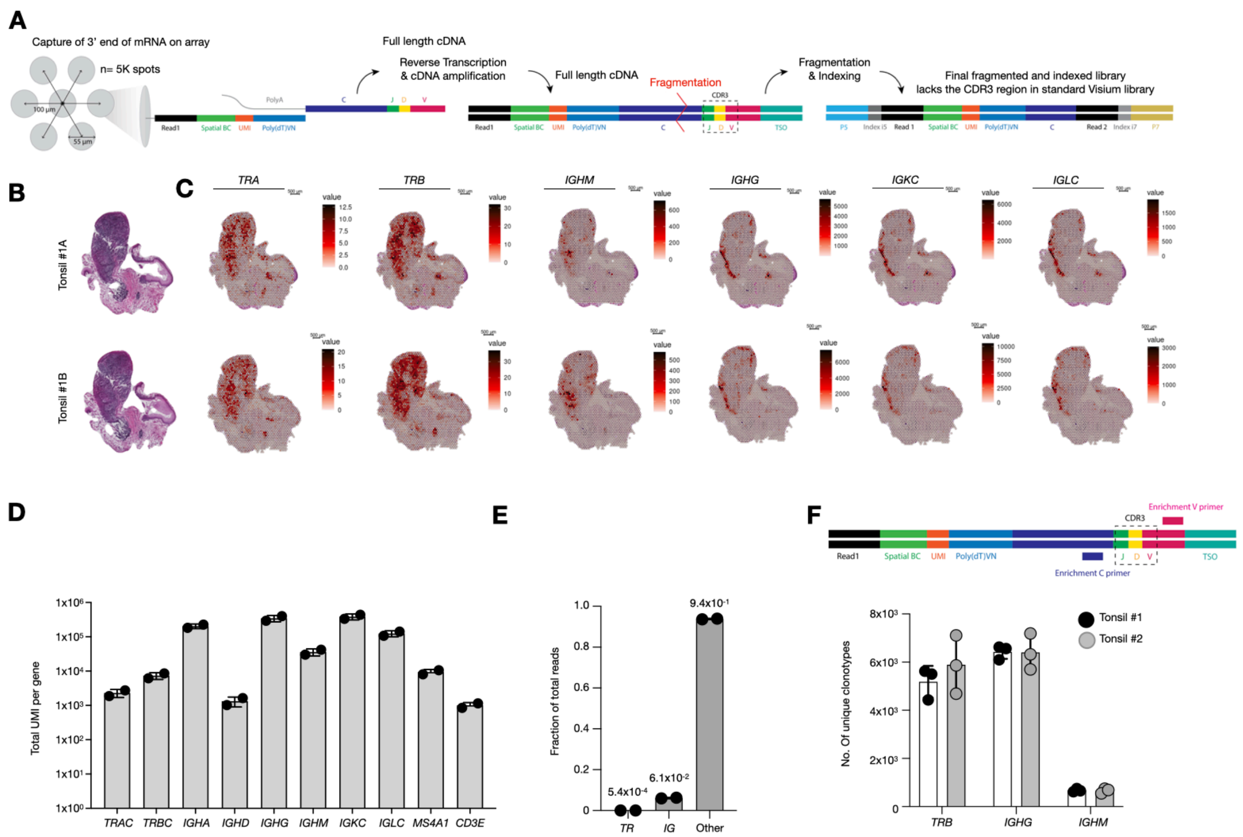


**fig. S1. TR and IG constant gene expression and clonal identity can be obtained from tonsil Spatial GEX libraries.**

(A) Diagram of how the CDR3 sequence is lost during the fragmentation step in standard Visium (Spatial GEX) libraries.

(B) Hematoxylin and eosin staining of two consecutive tonsil sections (Tonsil #1) used for Spatial GEX libraries.

(C) *TRA, TRB, IGHG, IGHM, IGKC, and IGLC* constant gene expression in tonsil Spatial GEX libraries. *IGHE*, which is a low-abundance IGH isotype in non-allergic individuals, was not detected (*12*, *24–26*).

(D) Total UMI count per gene for the constant genes presented in (C). B and T cell marker genes, *MS4A1* (encoding CD20), and *CD3E*, respectively, are included for reference.

(E) Fraction of total reads for TR and IG transcripts in tonsil Spatial GEX libraries. Mean value denoted above the bars.

(F) Outline of the PCR approach targeting the V and C genes of each receptor chain (*TRB*, *IGHG*, or *IGHM*) from full length Spatial GEX cDNA libraries (top). Here, the CDR3 region, but not the spatial barcode, is retained in the PCR product. The number of unique clonotypes amplified from Spatial GEX tonsil libraries (n=2 tissue sections; n=3 replicate PCR reactions) (bottom).

fig. S2

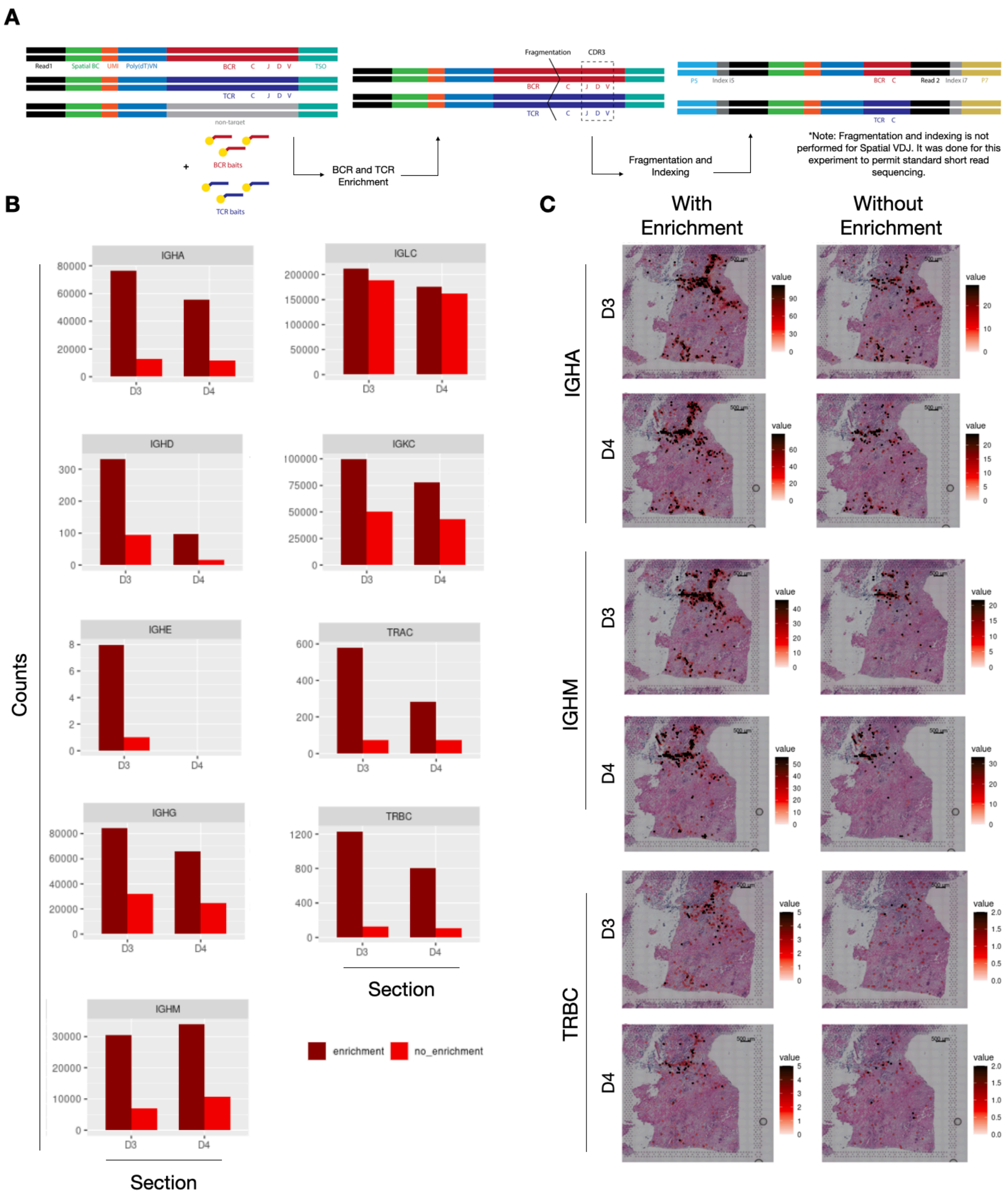


**fig. S2: Hybridization target capture of IG and TR constant regions increases the IG and TR gene counts in spatial transcriptomics libraries.**

(A) Outline of hybridization targeted capture of TCR (TR) and BCR (IG) transcripts.

(B) IG and TR constant gene counts with (dark red, left) or without (light red, right) enrichment

for the same breast cancer Spatial GEX library (section TumD3 and D4), Statistical significance between enriched versus non-enriched was p= 3.0x10-202 and p= 1.7x10-146 for TR and IG, respectively, calculated by Wilcoxon signed-rank test.
(C) Visualization of IGHA, IGHM, and TRB spatial gene expression in Spatial GEX libraries with (left) or without (right) target enrichment.

### fig. S3

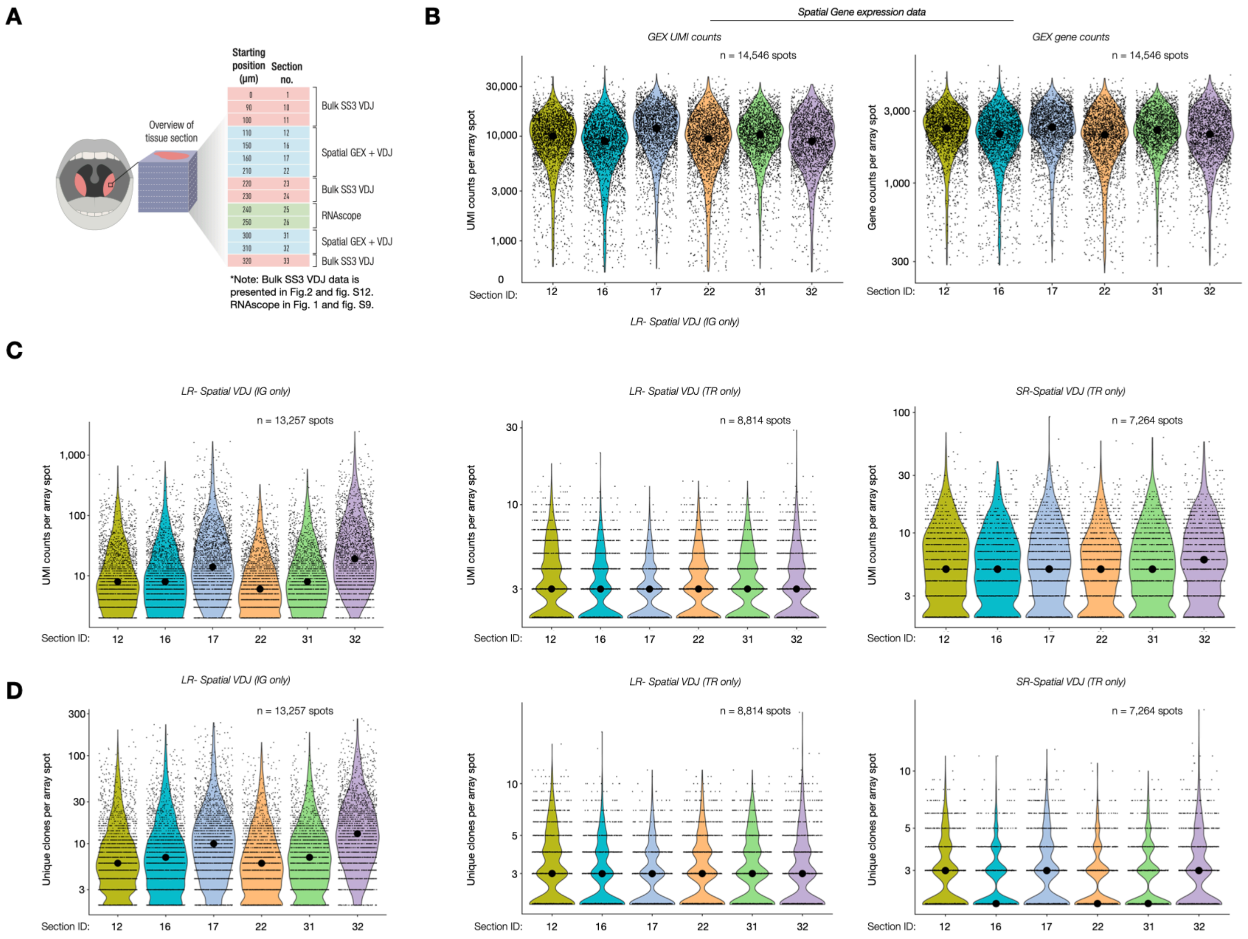


**fig. S3. Spatial GEX- and VDJ libraries capture consistent numbers of unique features (genes or clones) and total UMI counts per feature across replicate tonsil sections.**
(A) Experimental outline and sectioning strategy of samples analyzed in (A to D).
(B) Number of total UMI counts (left) and unique genes (right) per spot in Spatial GEX tonsil libraries.
(C) Number of total UMI counts per spot for LR-Spatial VDJ IG (left), LR-Spatial VDJ TR (middle) and SR-Spatial VDJ TR (right) tonsil libraries (same as in A).
(D) Number of unique clonal counts per spot for LR-Spatial VDJ IG (left), LR-Spatial VDJ TR (middle) and SR-Spatial VDJ TR (right) tonsil libraries (same as in A). In (B to D), only spots with non-zero values are shown and black dots indicate the median values per section.

**fig. S4**

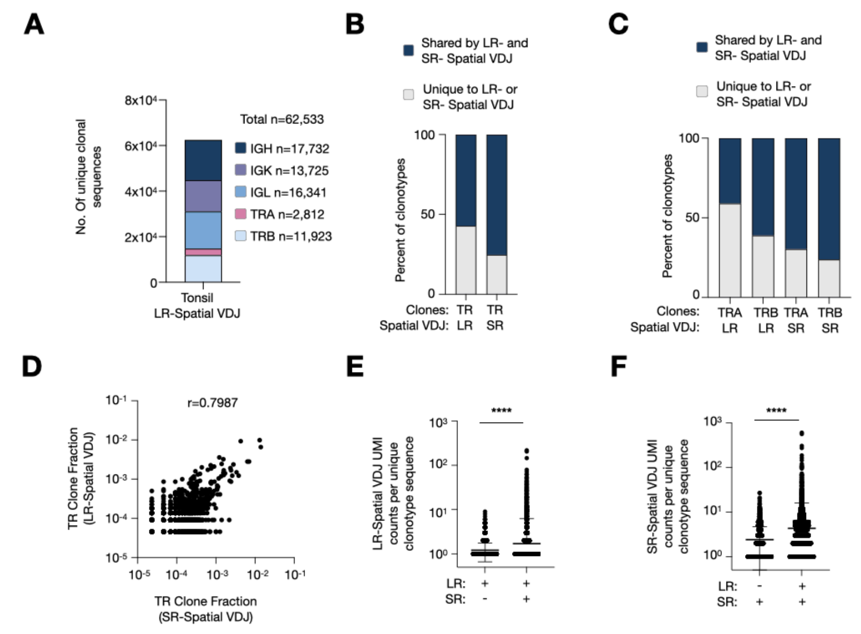


**fig. S4. Spatial VDJ captures a substantial number of TR and IG clonotypes in human tonsil with comparable results across LR- and SR- Spatial VDJ.**

(A) Total number of unique clonotypes across all six tonsil replicate sections split by antigen receptor chain.

(B) Percent shared and not shared TR clonotype sequences between LR- and SR- Spatial VDJ from the same samples.

(C) Percent shared and not shared TRA or TRB clonotype sequences between LR- and SR- Spatial VDJ from the same samples.

(D) TR Clone fraction of clonal sequences shared between LR- and SR- Spatial VDJ datasets. Pearson correlation coefficient is denoted on graph (r).

(E) LR-Spatial VDJ UMI count for each TRB clonotype present in only LR- Spatial VDJ or both LR- and SR-Spatial VDJ datasets n=6 replicate sections. Statistical significance was calculated using the Mann-Whitney test.

(F) SR-Spatial VDJ UMI count for each TRB clonotype present in only SR- Spatial VDJ or both LR- and SR-Spatial VDJ datasets. Statistical significance was calculated using the Mann-Whitney test.

For all panels: n=6 replicate sections from one human tonsil.

fig. S5

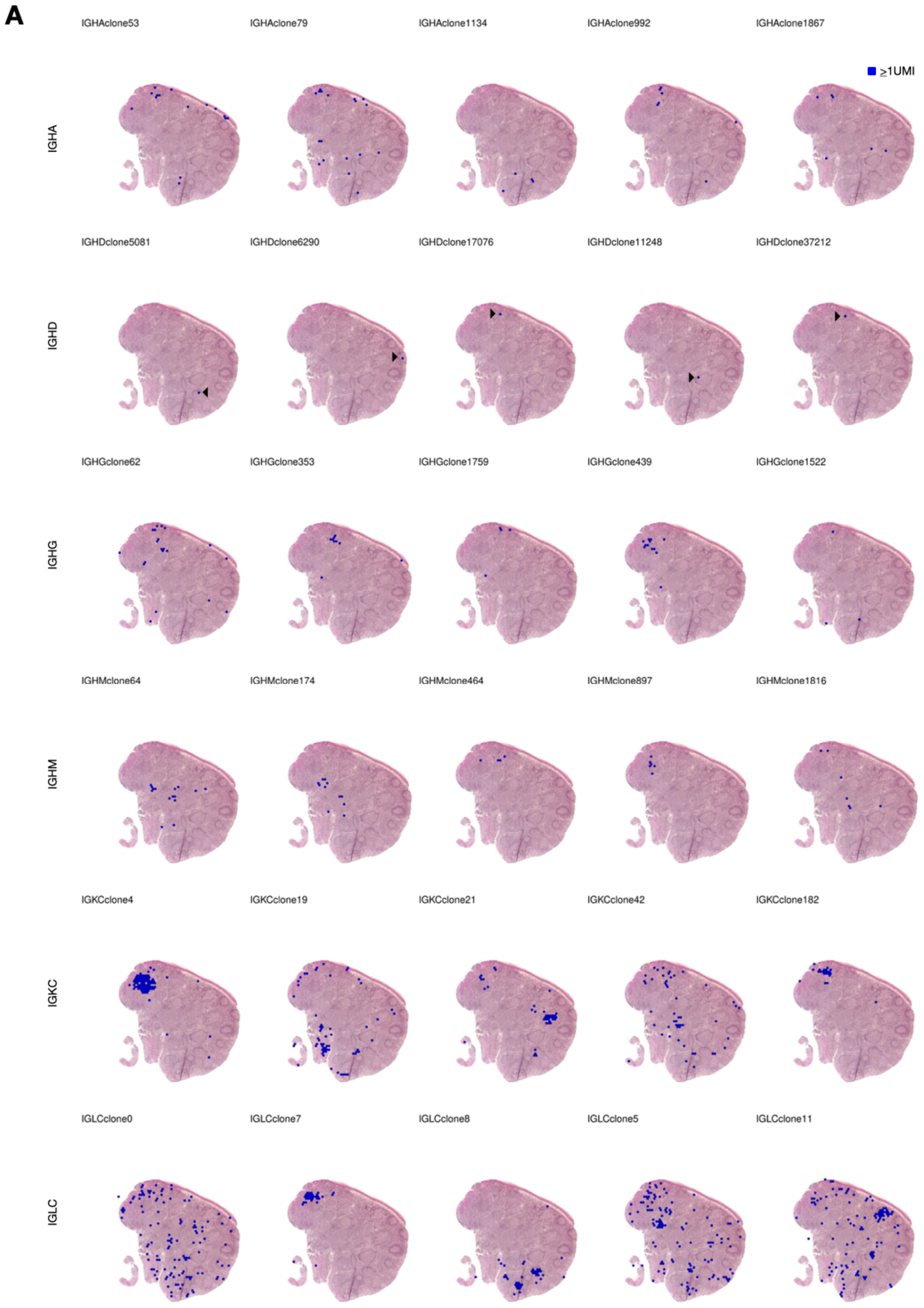


**fig. S5. Individual IG clonotypes have distinct spatial distribution.**
(A) Spatial distribution of selected individual clonal sequences for each IG constant chain visualized on one representative human tonsil section (section no. 17).

### fig. S6

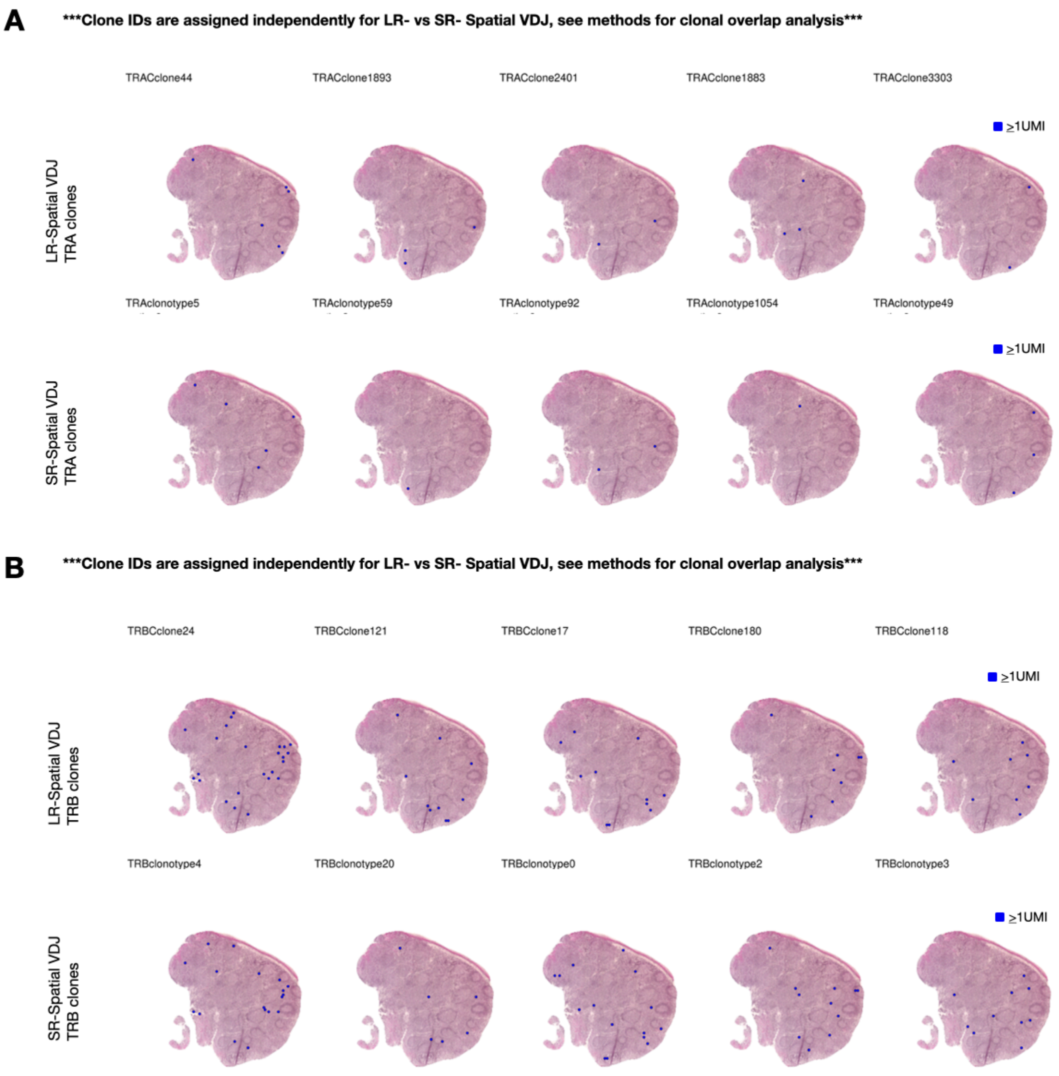


**fig. S6. Individual TRB clonotypes show distinct spatial distributions that match between LR- and SR-Spatial VDJ.**

(A and B) Spatial distribution of selected individual clones for (A) TRA and (B) TRB clones visualized on one representative human tonsil section (section no. 17). For each LR-Spatial VDJ clonotype (top row), the matched clonotype (identical nucleotide CDR3 and J gene call) is shown for SR- Spatial VDJ (bottom row).

### fig. S7

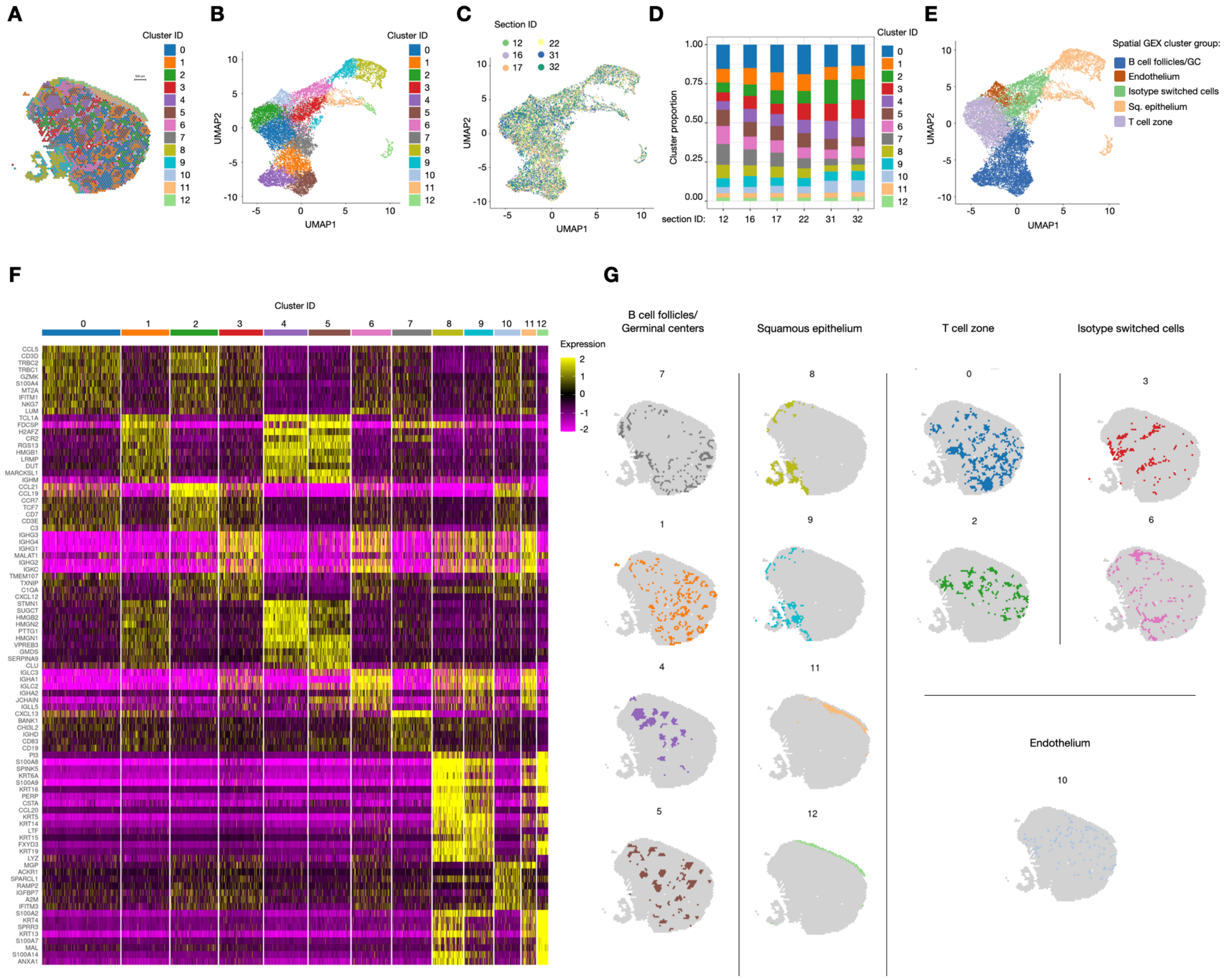


**fig. S7. Spatial transcriptomics of human tonsil identifies canonical features of lymphoid tissue anatomy.**

(A) Tonsil spatial gene expression (Spatial GEX) clusters plotted on one human tonsil section (representative section no. 17). Scale bar = 500 µm.

(B) UMAP of Spatial GEX data showing each individual cluster.

(C) UMAP of Spatial GEX data showing the contribution from each replicate tonsil section (n=6).

(D) The proportion of each cluster across replicate tonsil sections.

(E) UMAP of Spatial GEX cluster groups (annotated based on shared gene enrichment profiles). Note that the individual clusters in each group were also proximal in the UMAP embedding, supporting their shared expression patterns.

(F) Heatmap showing enriched genes for each cluster.

(G) Individual Spatial GEX clusters visualized on tonsil section 17 organized according to the cluster groups.

### fig. S8

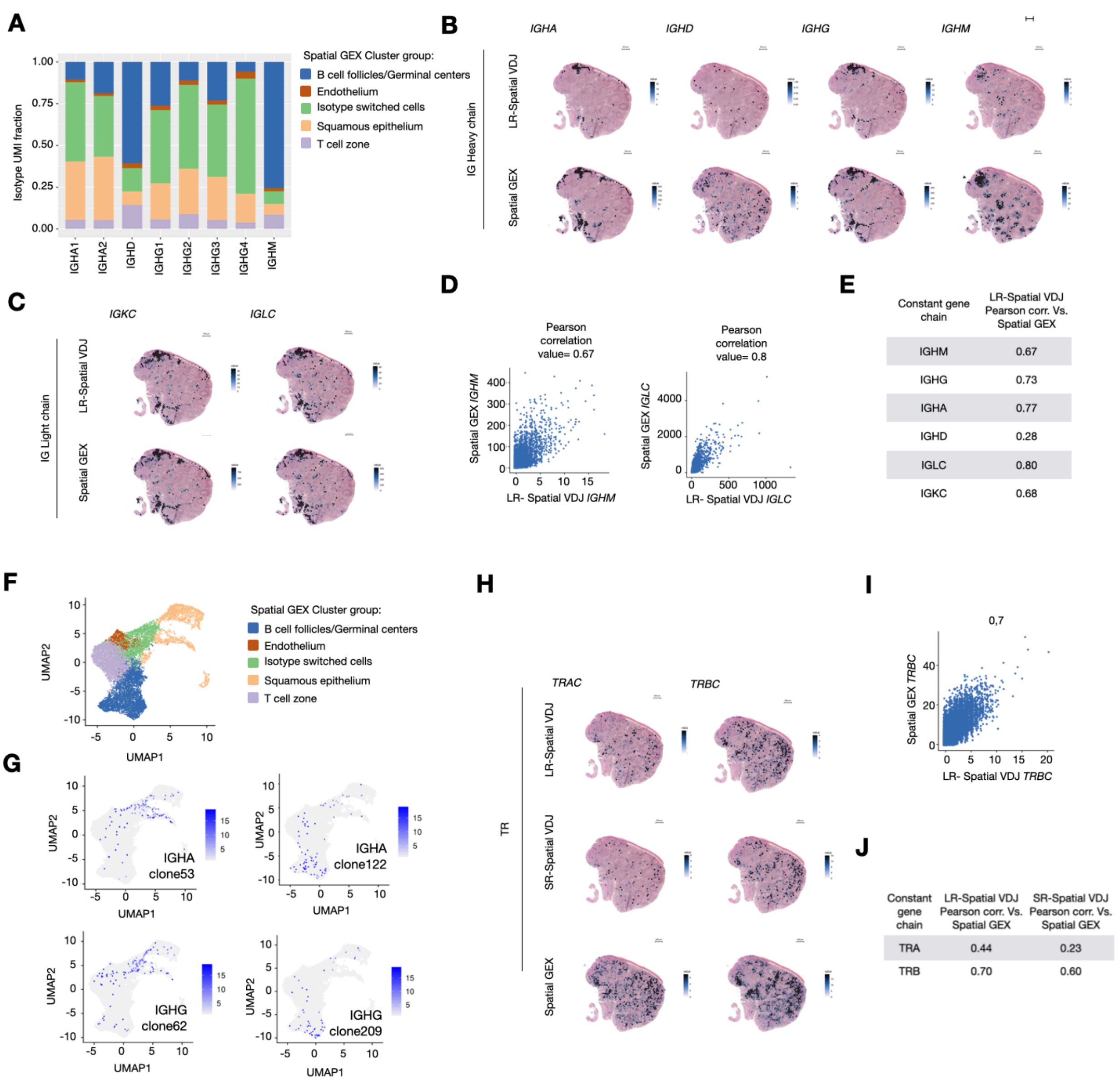


E

| Constant gene chain | LR-Spatial VDJ Pearson corr. Vs. Spatial GEX |
|---|---|
| IGHM | 0.67 |
| IGHG | 0.73 |
| IGHA | 0.77 |
| IGHD | 0.28 |
| IGLC | 0.80 |
| IGKC | 0.68 |

J

| Constant gene chain | LR-Spatial VDJ Pearson corr. Vs. Spatial GEX | SR-Spatial VDJ Pearson corr. Vs. Spatial GEX |
|---|---|---|
| TRA | 0.44 | 0.23 |
| TRB | 0.70 | 0.60 |

**fig. S8: Spatial VDJ captures IG and TR clonal spatial patterns that match Spatial GEX constant gene expression.**

(A) IGH isotype fraction of LR-Spatial VDJ reads across spatial GEX clusters. *IGHE* expression was not detected, in agreement with tonsillar B cell single cell (*12*) and Spatial GEX of another tonsil sample (fig. S1).

(B-C) Spatial distribution of (B) IGH or (C) IGK/L constant gene expression in LR- Spatial VDJ (top) and Spatial GEX libraries (bottom) in tonsil tissue (representative section no. 17). Scale bar = 500 µm.

(D) Dotplot showing *IGHM,* and *IGLC* counts per array spot for Spatial- VDJ versus GEX

datasets in tonsil tissue.
(E) Pearson correlation coefficient (r) for LR-Spatial VDJ- versus Spatial GEX data for each IG constant gene.
(F) Spatial GEX cluster groups visualized on the UMAP embedding. Also in fig. S7E.
(G) Representative IGHA (top) and IGHG (bottom) clones plotted on the Spatial GEX UMAP space.
(H) Spatial distribution of TR constant gene expression in LR- Spatial VDJ (top), SR-Spatial VDJ (middle) and Spatial GEX libraries (bottom) in tonsil tissue. Scale bar represents 500µm.
(I) Dotplot showing *TRBC* counts for LR-Spatial VDJ- versus GEX datasets in tonsil tissue.
(J) Pearson correlation coefficient (r) between LR-Spatial VDJ- versus GEX data for each TR constant gene
All panels: when applicable, the data for IG isotypes IGHA1-2 and IGHG1-4 are merged into 'IGHA' and 'IGHG', respectively. Similarly, TRBC1 and TRBC2 data are merged to 'TRBC'.

**fig. S9**

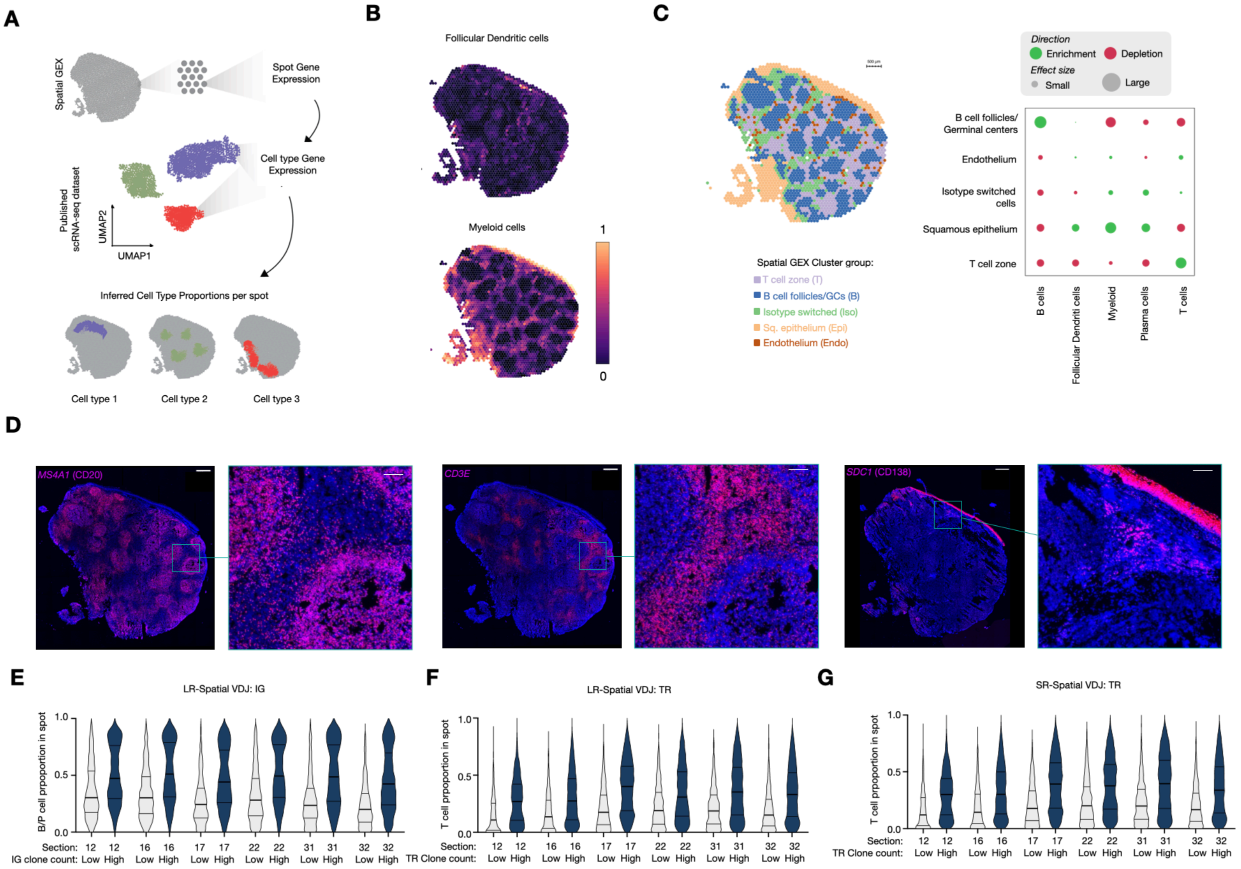


**fig. S9. Single cell deconvolution using stereoscope delineates spatial organization of cell types across human tonsil tissue that matches Spatial VDJ clonal capture.**

(A) Outline of cell deconvolution of Spatial GEX data using stereoscope, guided by published tonsil scRNA-seq dataset (*12*).

(B) Visualization of major cell type distribution within tonsil defined by single cell deconvolution. See also Fig. 1I for T, B, and plasma cell distribution.

(C) Groups of Spatial GEX cluster groups plotted on tonsil tissue (left) and major cell type enrichment and depletion across Spatial GEX cluster groups (right). Data from section 17 is shown.

(D) Zoom-in of RNAscope staining of *MS4A1* (B cell marker), *CD3E* (T cell marker), and *SDC1* (Plasma cell marker) in tonsil tissue sections shown in Fig. 1 (the entire tissue is reproduced here for clarity). Of note, SDC1 is also highly expressed by tonsil squamous epithelial cells, as expected (*27*). Scale bar = 500μm. See fig. S3A for experimental outline.

(E to G) Cell type proportion for spots containing high or low clonotype counts (E: LR-Spatial VDJ IG; F: LR-Spatial VDJ TR; G: SR- Spatial VDJ TR) split across all replicate tissue sections (from the same human tonsil sample).

### fig. S10

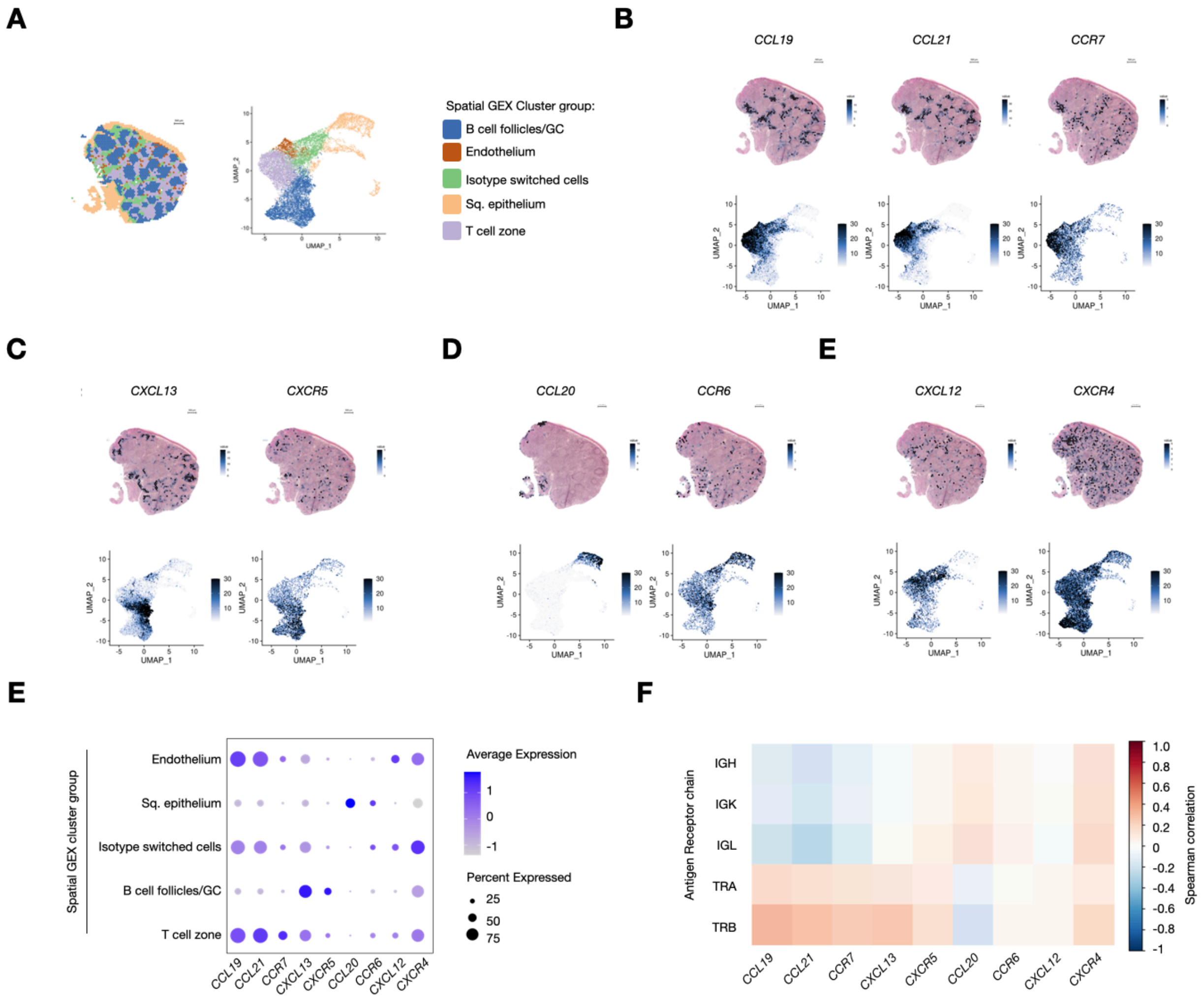


**fig. S10. Spatial distribution of chemokines and chemokine receptors with known roles in lymphoid tissue organization.**

(A) Spatial GEX cluster groups presented on tissue section and the UMAP embedding for reference (also in Fig. 1 and fig. S7-9).

(B to E) Distribution of individual chemokine ligand/receptor pairs shown on the tonsil tissue and UMAP embedding.

(E) Dotplot showing the fold change average expression of the chemokine ligands/receptors shown in B to E across the Spatial GEX cluster groups (i.e. the anatomical regions detected by Spatial GEX).

(F) Spearman correlation between LR-Spatial VDJ antigen receptor chains and chemokine ligands/receptors spatial gene expression levels. Red color indicates positive correlation values and blue indicates negative correlation values.

**fig. S11**

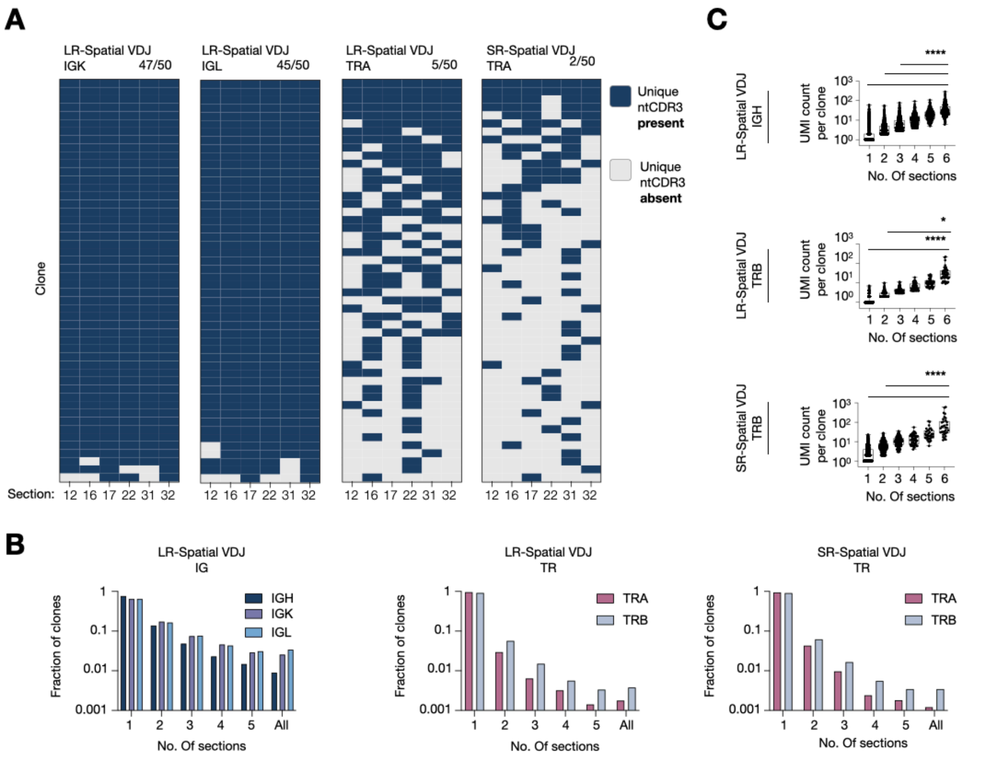


**fig. S11. Spatial VDJ reproducibly captures expanded clones across human tonsil sections.**
(A) Presence or absence of the top 50 IGK, IGL, or TRA clonotypes (based on UMI count and sorted on number of sections) across replicate tonsil sections (n=6).
(B) Fraction of clonotypes detected in 1, 2, 3, 4, 5, or all (n=6) replicate tonsil sections for the receptor chains listed.
(C) UMI count for each clone present in 1, 2, 3, 4, 5, or all (n=6) replicate tonsil sections for each receptor chain and dataset listed. Statistical significance between for UMI counts per clone found in all replicate sections versus those found in 1-5 sections is shown here and was calculated using the Kruskal Wallis test followed by Dunn's multiple comparison's test.
For the entire figure: when applicable, the data for TRBC1 and TRBC2 clones are merged to 'TRB'.
*, **** denotes $p<0.05$ and $p<0.0001$, respectively.

### fig. S12

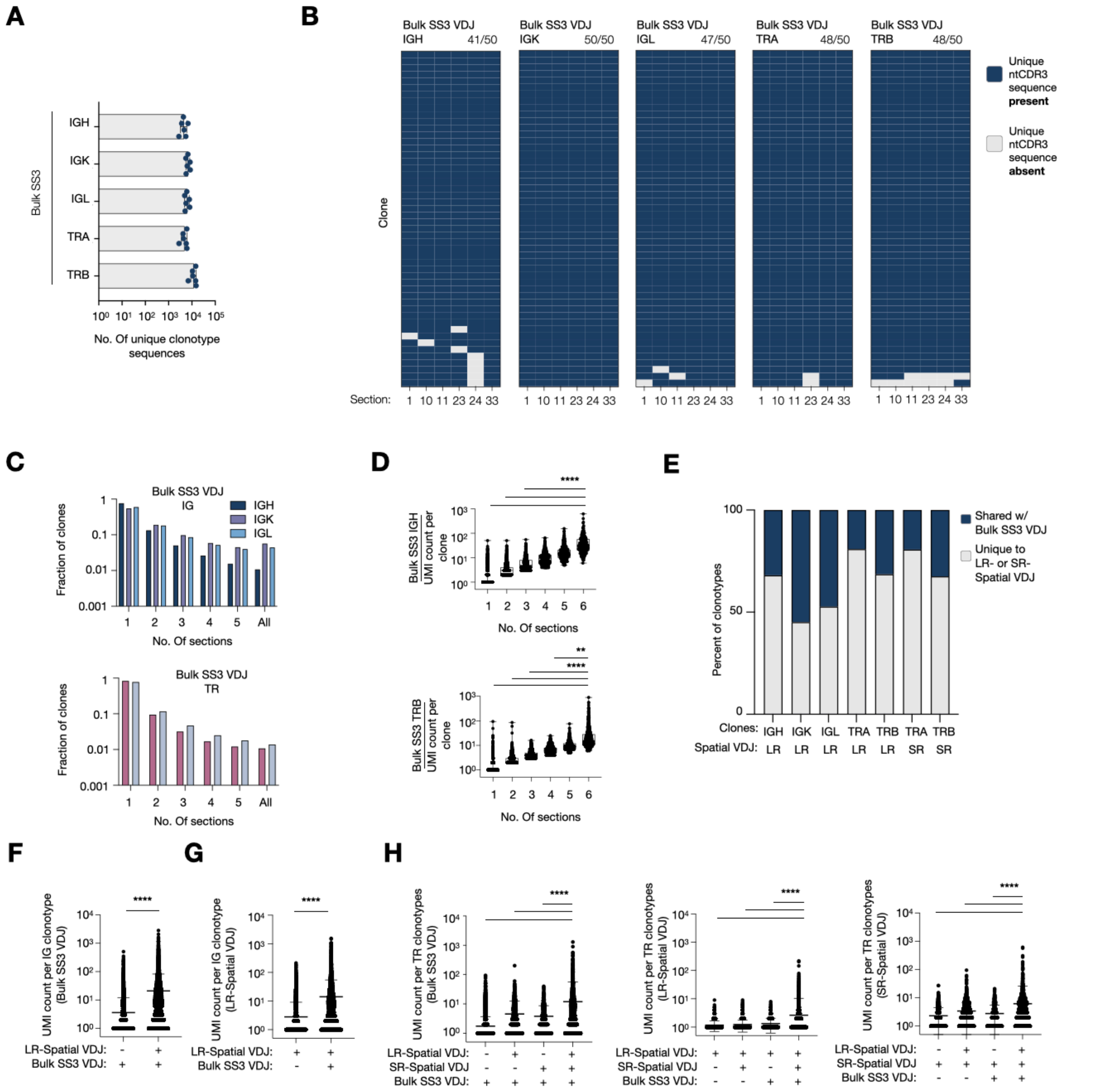


**fig. S12. TR and IG clones are validated by bulk SS3 VDJ analysis from matched tonsil sections.**

(A) Unique clonotype sequences amplified from tonsil section RNA (n=6 replicates) using an SS3-based protocol adapted for bulk VDJ analysis. These sections were from the same tonsil sample analyzed by LR- and SR- Spatial VDJ. Experimental outline shown in fig. S3A.

(B) Presence or absence of the top 50 IGH, IGK, IGL, TRA or TRB clonotypes (based on UMI count and sorted on number of sections) across replicate tonsil sections from bulk SS3 VDJ analysis.

(C) Fraction of IG (top) or TR (bottom) clonotypes detected by Bulk SS3 VDJ in 1, 2, 3, 4, 5, or all (n=6) replicate tonsil sections for the receptor chains listed.

(D) UMI count for each IGH (top) or TRB (bottom) clone present in 1, 2, 3, 4, 5, or all replicate tonsil sections for each receptor chain listed for Bulk SS3 VDJ. Statistical significance between for UMI counts per clone found in all 6 replicate sections versus those found in 1-5 sections is shown here and was calculated using the Kruskal Wallis test followed by Dunn's multiple comparison's test.
(E) Percent of clonotypes split by constant receptor chain that were uniquely found in or shared between LR- or SR- Spatial VDJ and bulk SS3 VDJ.
(F) LR-Spatial VDJ UMI count for each IG clonotype present in only LR- Spatial VDJ or both LR- Spatial VDJ and Bulk SS3 VDJ datasets. Statistical significance was calculated using a Mann-Whitney test.
(G) Bulk SS3 VDJ UMI count for each IG clonotype present in only Bulk SS3 VDJ or both LR-Spatial VDJ and Bulk SS3 VDJ datasets. Statistical significance was calculated using a Mann-Whitney test.
(H) UMI count (LR-Spatial VDJ, SR-Spatial VDJ, or Bulk SS3 VDJ) for each TR clonotype present in each single or combination of indicated dataset. Statistical significance between UMI counts for clones shared between all three datasets versus all other combinations is shown here and was calculated using the Kruskal Wallis test followed by Dunn's multiple comparison's test.
**, **** denotes $p<0.01$ and $p<0.0001$, respectively.

### fig. S13

**A**

| Patient ID | Her2 Status | HER2 IHC score | ER (%) | PR (%) | Ki67 (%) | Biopsy ID | Spatial GEX | LR-Spatial VDJ | SR- Spatial VDJ | Single cell VDJ | Single cell GEX |
|---|---|---|---|---|---|---|---|---|---|---|---|
| P1 | Positive | 3+ | 30 | 0 | 79 | Region (Reg) A, B, C, D1-4, E | RegC, D1-4, E | RegC, D1-2, E | RegC and E | All regions | All regions |
| P2 | Positive | 3+ | 90 | 90 | 70 | Region (Reg) A, B, C, and D | RegA and B | RegA and B | RegA and B | All regions | All regions |

**B**

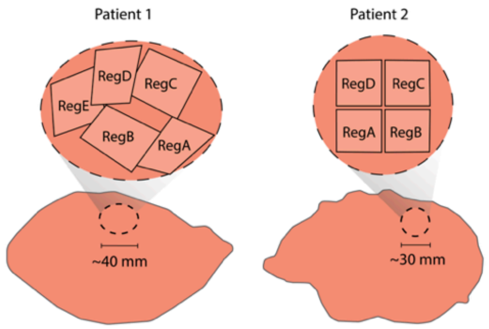


**fig. S13: Sample and analysis overview of human breast cancer samples.**
(A) Table containing sample IDs, patient clinical information, and performed analyses.
(B) Overview of regional biopsies from each patient sample.

**fig. S14**

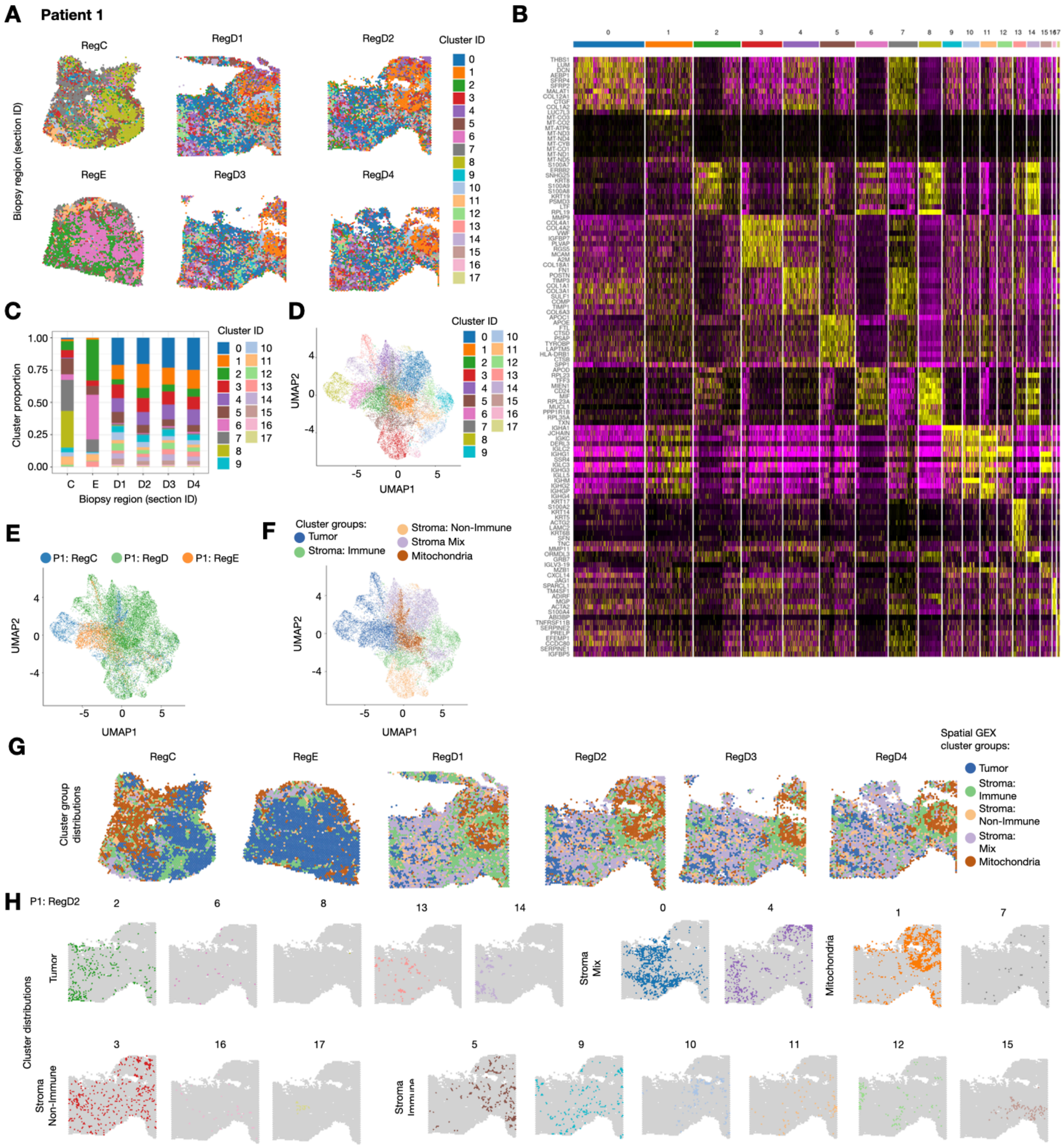


**fig. S14. Spatial transcriptomics of human breast cancer (Patient 1) maps tumor and stromal gene expression patterns.**

(A) Spatial gene expression clusters (n=18) plotted on breast cancer samples (Patient 1, regions: C, D, and E). Four replicate sections were performed for region D (1-4), where D1/D2 and D3/D4 represent pairs of adjacent sections.

(B) Heatmap showing enriched genes for each cluster.

(C) The proportion of each cluster across sampled regions. Samples D1-D4 are replicate sections

from the same biopsy region.
(D) UMAP of Spatial GEX data showing each individual cluster.
(E) UMAP of Spatial GEX data showing the contribution from each sampled region (C, D, and E). Replicates from TumD are merged here.
(F) UMAP of Spatial GEX data showing the cluster groups (n=5, annotated based on shared gene enrichment profiles). The individual Spatial GEX clusters were categorized based on shared features resulting in the cluster groups: ‘Tumor’, ‘Stroma: Non-immune’, ‘Stroma: Immune’, ‘Stroma: Mix’ (enriched for both non-immune and immune-associated stromal features), and ‘Mitochondria’ (enriched for mitochondrial genes).
(G) Cluster groups visualized on Patient1 samples.
(H) All Individual clusters visualized on RegD2 listed according to the cluster groups.

### fig. S15

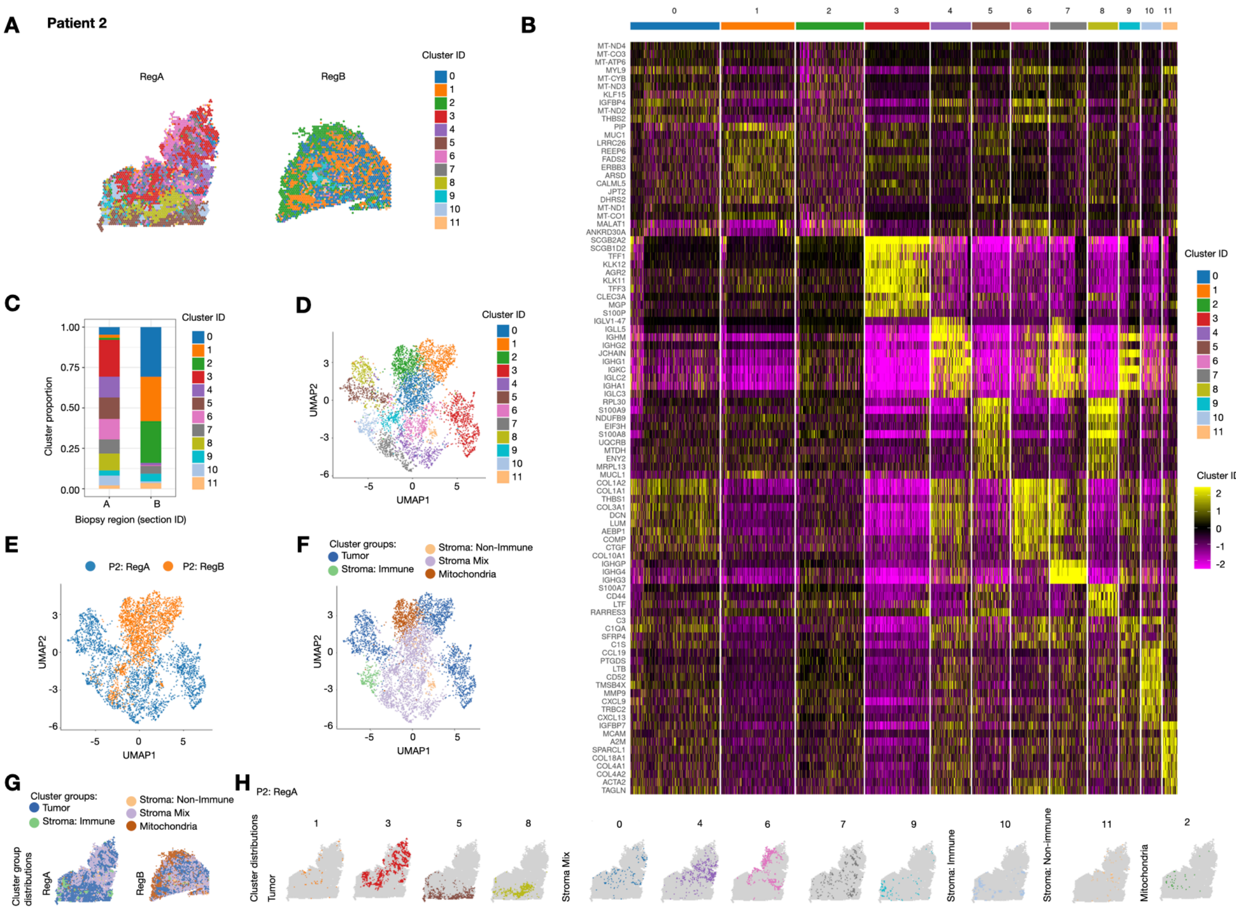


**fig. S15. Spatial transcriptomics of human breast cancer (Patient 2) maps tumor and stromal gene expression patterns.**

(A) Spatial gene expression clusters (n=12) plotted on breast cancer samples (patient 2, regions: A and B).

(B) Heatmap showing enriched genes for each cluster.

(C) The proportion of each cluster across sampled regions.

(D) UMAP of Spatial GEX showing each individual cluster.

(E) UMAP of Spatial GEX showing the contribution from each sampled region (A and B).

(F) UMAP of Spatial GEX data showing the cluster groups (n=5, annotated based on shared gene enrichment profiles). The individual Spatial GEX clusters were categorized based on shared features resulting in the cluster groups: ‘Tumor’, ‘Stroma: Non-immune’, ‘Stroma: Immune’, ‘Stroma: Mix’ (enriched for both non-immune and immune-associated stromal features), and ‘Mitochondria’ (enriched for mitochondrial genes).

(G) Spatial GEX cluster groups visualized on Patient2 samples.

(H) All Individual clusters visualized on RegA listed according to the cluster groups.

**fig. S16**

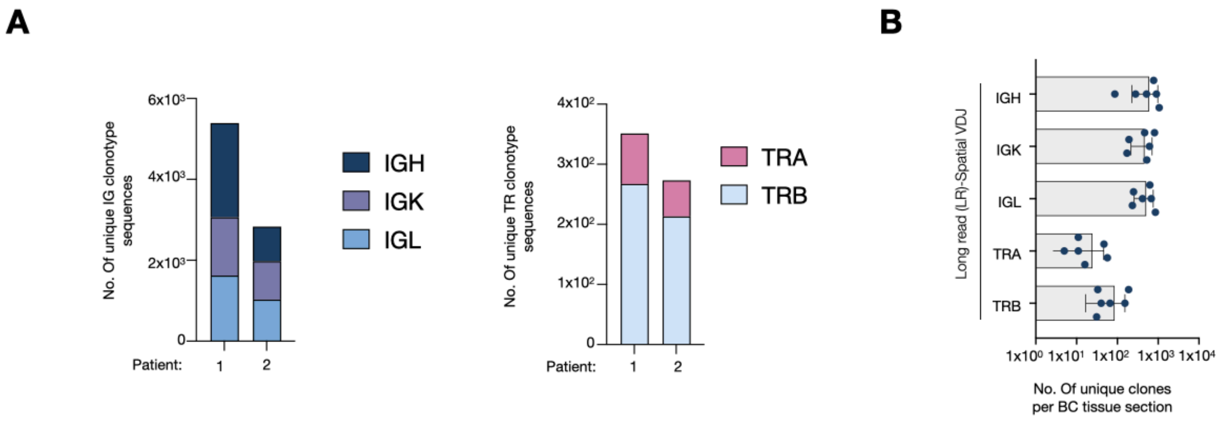


**fig. S16. Spatial VDJ permits high fidelity mapping of T and B cell clones in human breast tumor tissue.**

(A) Total number of unique clonotypes across all samples from each respective breast cancer sample split by antigen receptor chain.

(B) Unique clonal distribution pooled from LR-Spatial VDJ libraries from breast cancer samples: Patient1 (n= 3 sampled regions, with one section for RegC and E, and two replicate sections for RegD) and P2 (n= 2 sampled regions, one section prepared per region).

fig. S17

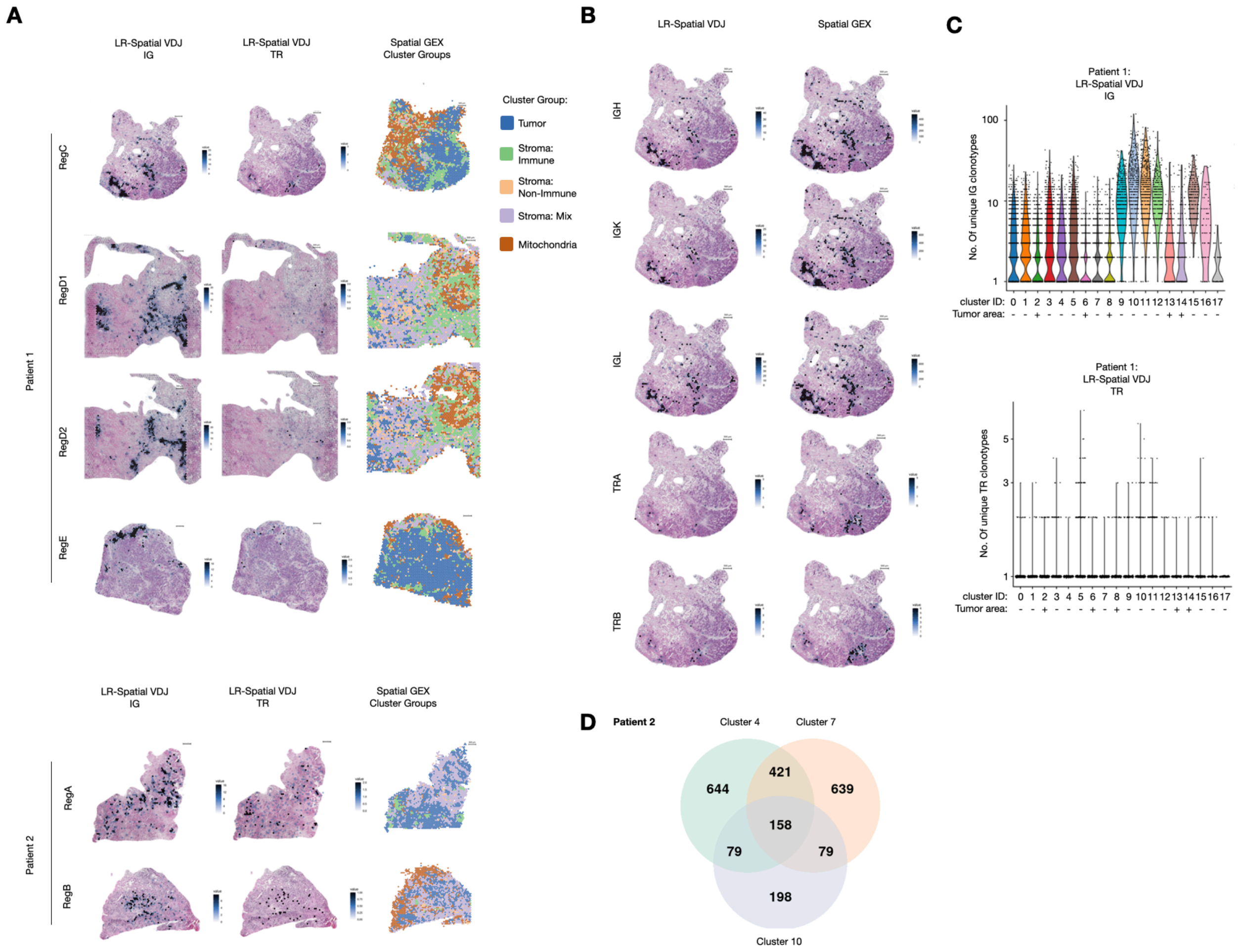


**fig. S17. Spatial VDJ-captured Clonal spatial distribution is conserved between adjacent tissue sections and consistent with Spatial GEX data.**

(A) Unique clonal distribution for LR- Spatial VDJ prepared from multiple regions from two individual patients (P1 and P2). Spatial GEX cluster groups are shown to the right for reference (also in fig. S14 and S15).

(B) Spatial distribution of IG constant gene expression in LR- Spatial VDJ (top) and Spatial GEX libraries (bottom) in Patient 1 RegC.

(C) No. of unique IG (left) and TR (right) clonotypes per spot in different Spatial GEX clusters ('+' denotes tumor cluster). Data from all regions (RegC, D, and E) from P1.

(D) Shared IG clonotypes between cluster 4, 7, and 10 for P2. Related to Fig. 3C.

Abbreviations: Tum - Tumor

**fig. S18**

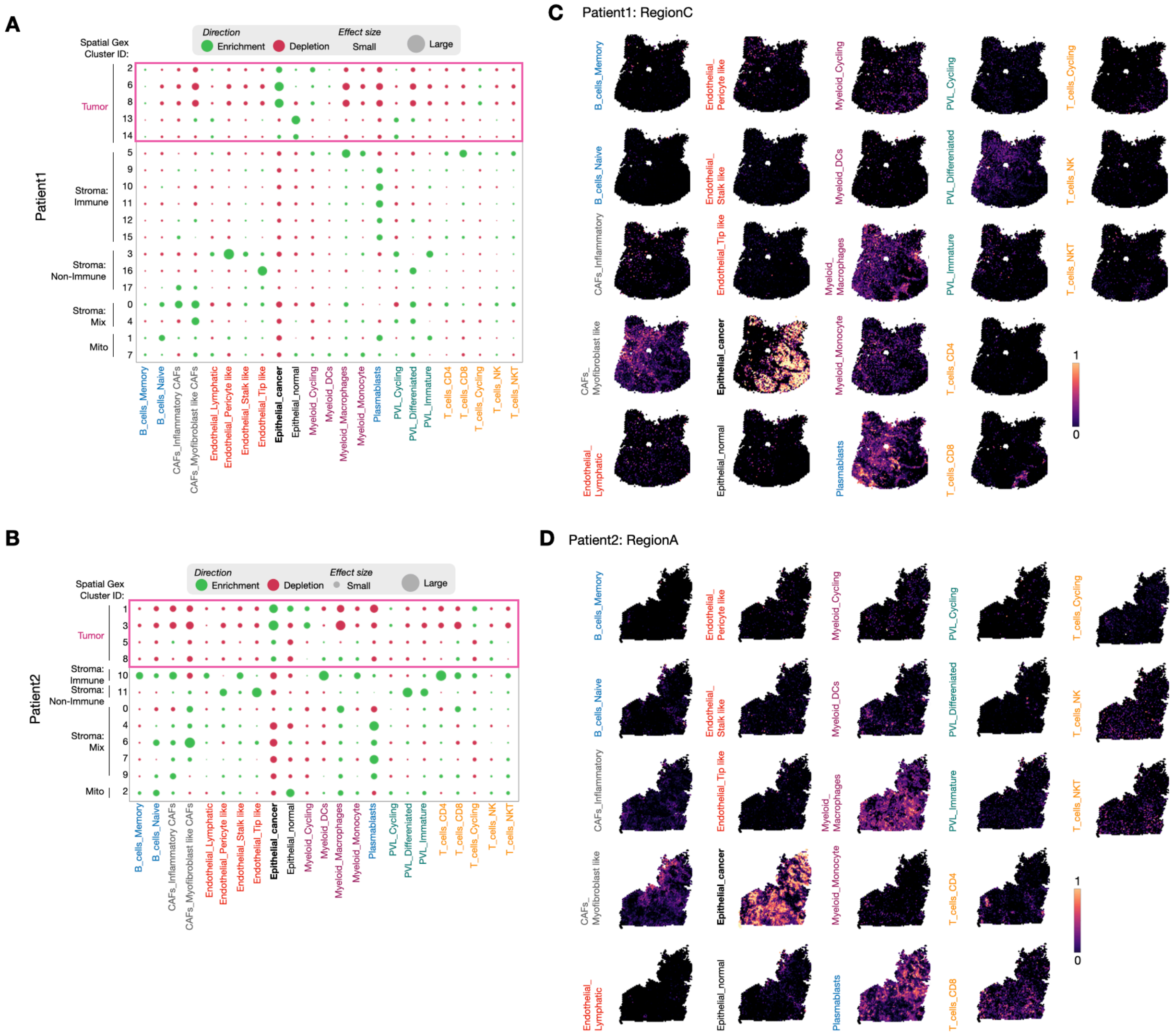


**fig. S18. Enrichment/depletion of cell types across all breast cancer Spatial GEX clusters.** (A) Cell type enrichment/depletion across the Spatial GEX individual clusters (sorted by cluster group). 'Minor' cell tier plotted as described in (Methods). Region C, D, and E datasets pooled from Patient1.

(B) Cell type (left) enrichment/depletion across the Spatial GEX individual clusters (sorted by cluster group). Region A and B datasets pooled from Patient2. 'Minor' cell tier plotted as described in (Methods).

(C and D) Visualization of cell type distribution within breast cancer tissue, (C) Patient1: RegC and (D) Patient2: RegA, based on single cell deconvolution analysis. Abbreviations: PVL - perivascular, DCs - dendritic cells, CAFs - cancer associated fibroblasts, NK - natural killer cells, NKT - Natural T-killer cell.

**fig. S19**

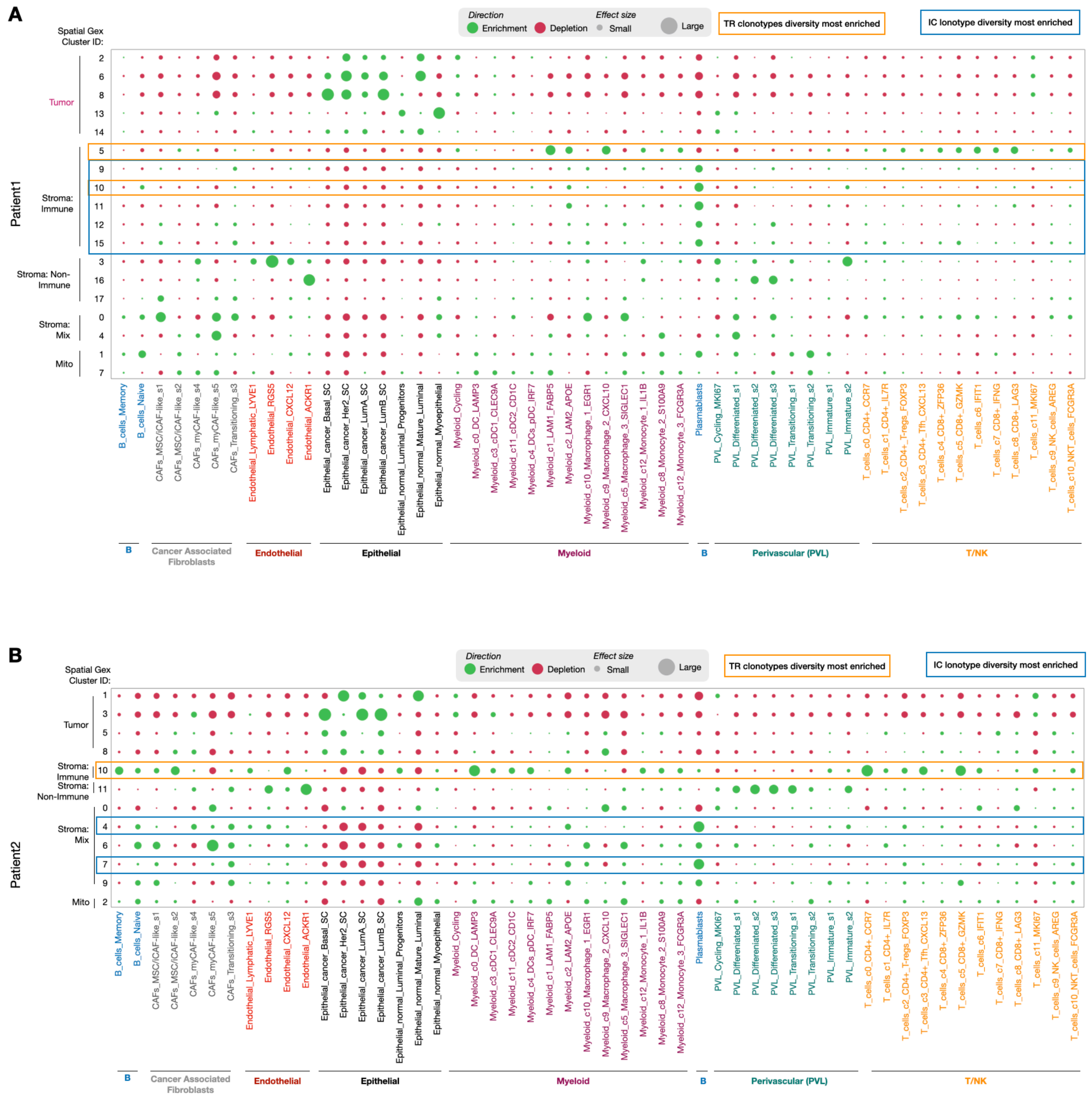


**fig. S19. Cell subset enrichment and depletion across all individual breast cancer Spatial GEX clusters.**

(A and B) Cell subset (B lineage cells, T cells, myeloid, and non-immune) enrichment and depletion enrichment/depletion across the Spatial GEX individual clusters (sorted by cluster group and cell type) based on single cell deconvolution data. Data pooled from multiple biopsy regions from (A) P1: Region C, D, and E, and (B) P2: Region A and B. Abbreviations: PVL - perivascular, DCs - dendritic cells (c = conventional, p = plasmacytoid), CAFs - cancer associated fibroblasts (my = myofibroblastic, i = inflammatory), NK - natural killer cells, NKT - Natural T-killer cell, MSC - mesenchymal stem cell.

fig. S20

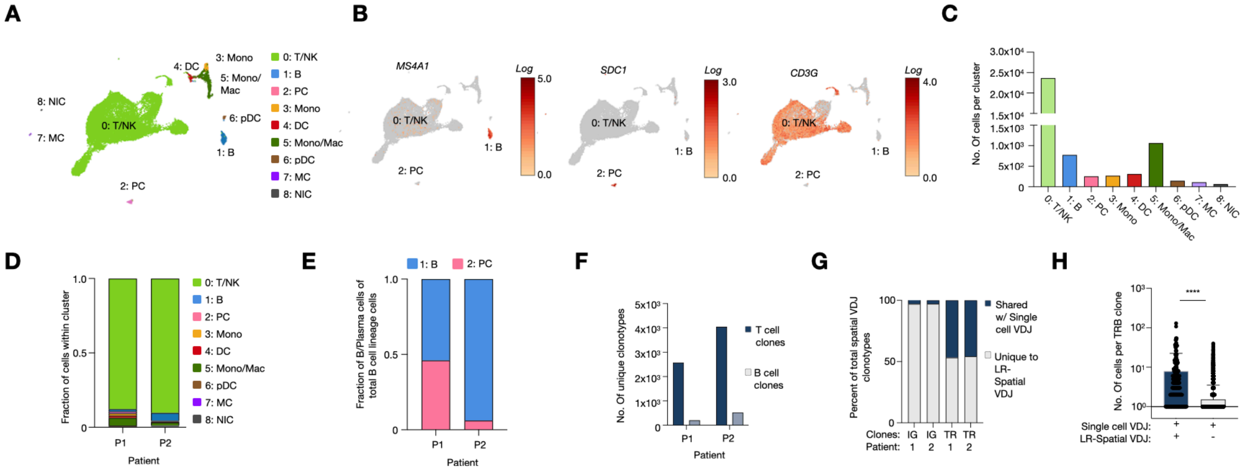


**fig. S20. Single cell immune gene expression analysis of breast tumor tissue.**

(A) UMAP of CD45$^{+}$ flowcytometry-sorted cells from two breast cancer samples analyzed with single cell RNA-seq.

(B) Expression of B cell (*MS4A1*), plasma cell (*SDC1*), and T cell (*CD3G*) marker genes visualized on the UMAP embedding.

(C) Number of cells per cell type cluster.

(D) The proportion of each cell type cluster across patients.

(E) The proportion of B versus Plasma cells per patient.

(F) The number of unique T or B cell clonotypes per patient.

(G) Percent shared and not shared IG or TR clonotype sequences between LR- Spatial VDJ and the single cell VDJ dataset.

(H) Number of cells per TRB clone between clones shared or not between LR-Spatial VDJ and the single cell VDJ datasets.

Abbreviations: T/NK - T or NK cells; B - B cells; Mono - Monocytes; Mac - Macrophages; DC - Dendritic Cells; PC - Plasma cells; pDC - plasmacytoid Dendritic cells; MC - Mast cells; NIC - Non-Immune Cells

fig. S21

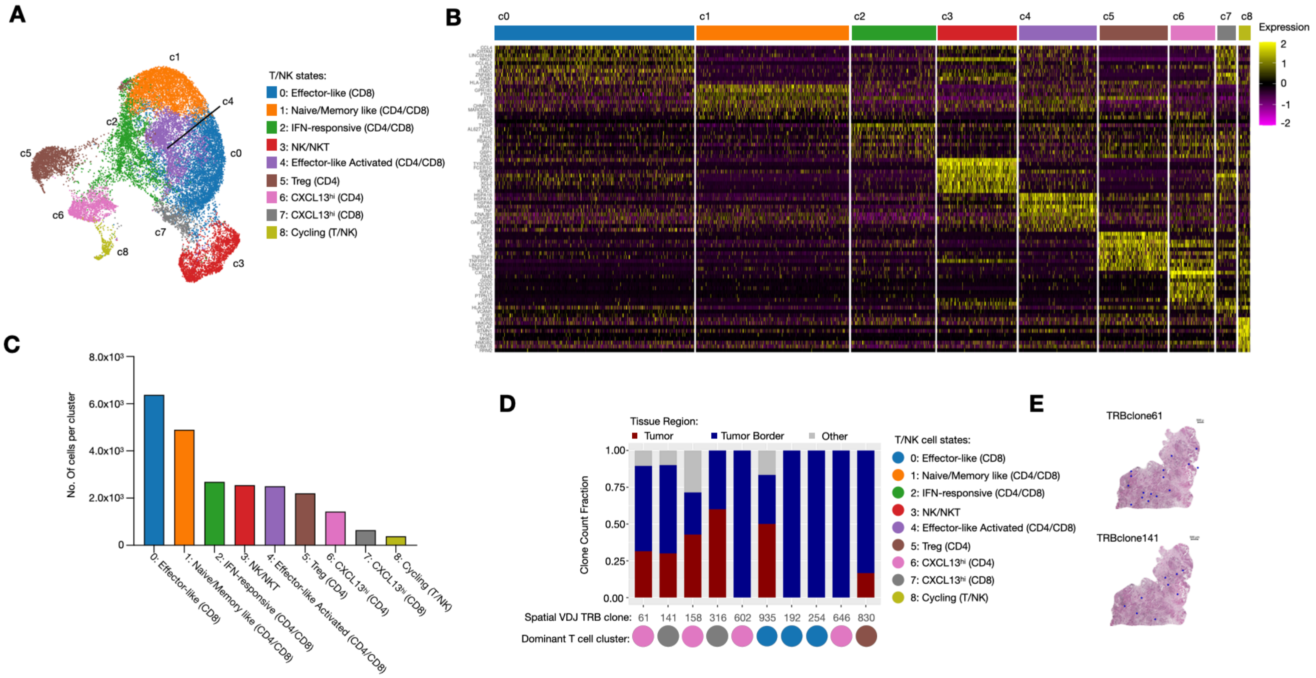


**fig. S21. Single cell T/NK cell state gene expression analysis of breast tumor tissue.**
(A) UMAP of T/NK cell clusters from CD45$^+$ flowcytometry-sorted cells from two breast tumors analyzed with single cell RNA-seq.
(B) Differentially expressed genes across T/NK cell clusters defined in (A).
(C) Number of cells per cluster.
(D) The UMI distribution across tissue location and the dominant T/NK cell state cluster distribution per the ten most expanded TRB clones in the Spatial VDJ dataset. The dominant T/NK cell cluster is defined as the cluster to which >50% of the cells belonged to for each clone.
(E) Spatial distribution on tumor tissue of two representative clones from (D).

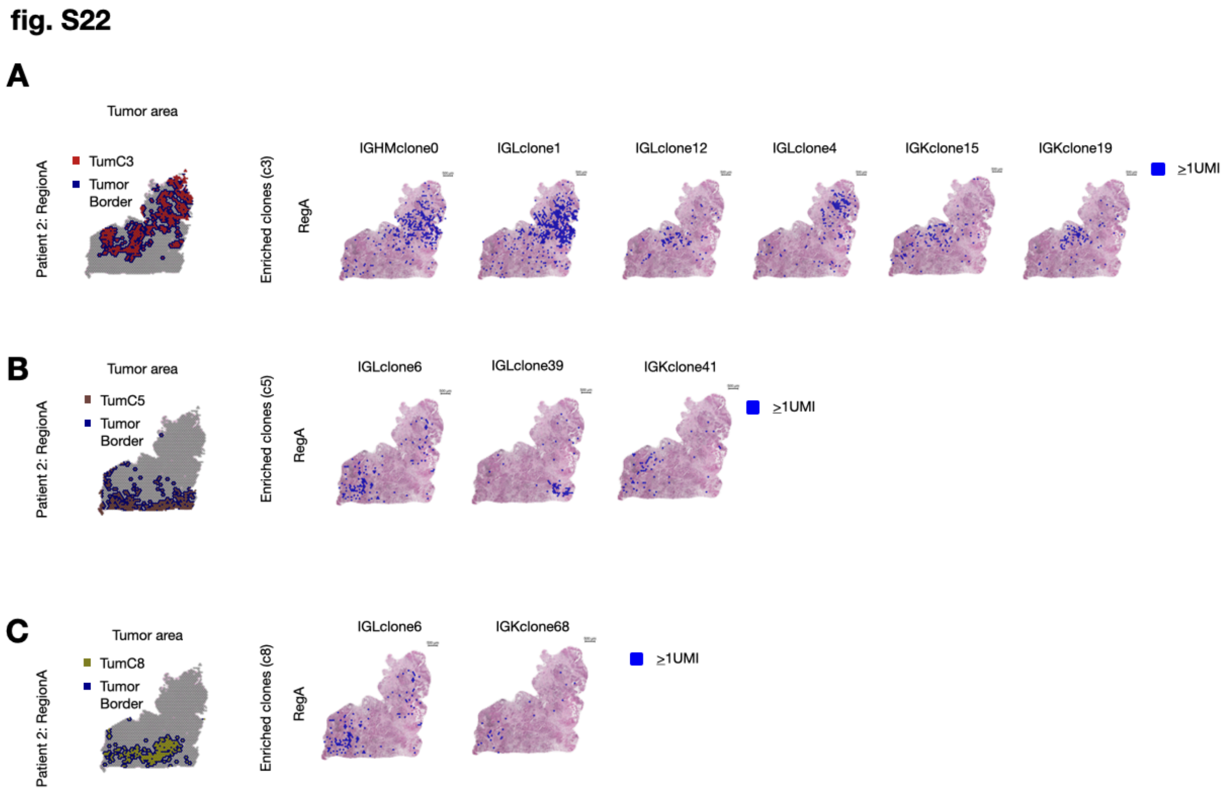


**fig. S22. Distinct spatial segregation of IG clones along different breast tumor areas.** (A to C) Tumor c3, 5, and 8 clusters and their surrounding borders visualized on Patient RegA (left). The spatial distribution (right) of all significantly enriched IG clonotypes per (A) Tum-c3, (B) Tum-c5, and (C) Tum-c8. Spatial distribution of representative IG clonotypes within each respective area (right). See Fig. 3 for additional data.

fig. S23

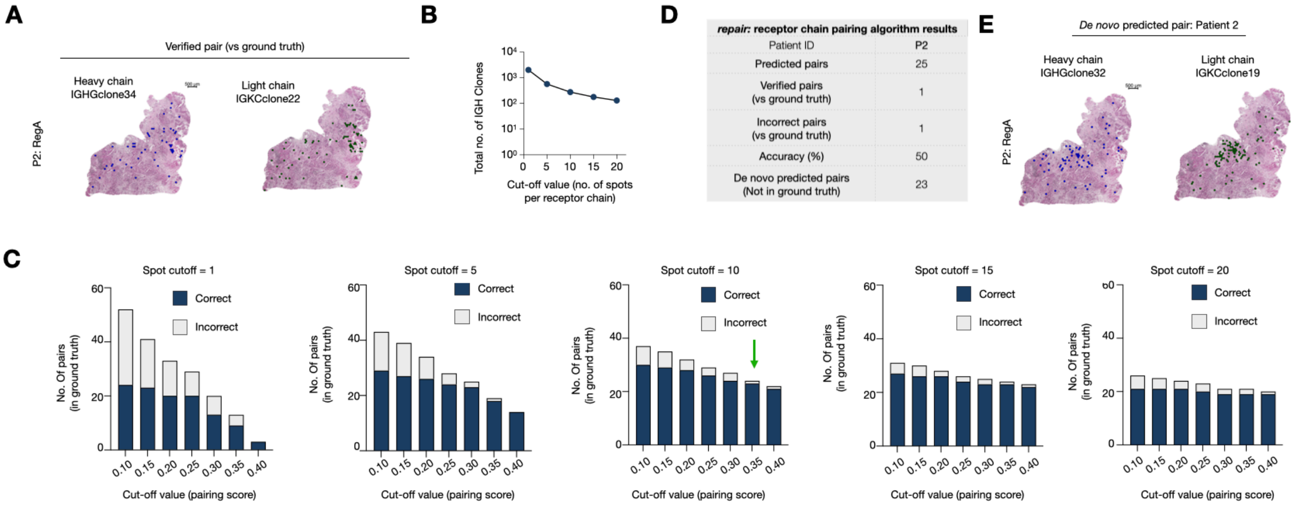


**fig. S23. Paired receptors show similar spatial distributions.**

(A) Spatial distribution of representative IG paired receptors from breast cancer Patient2 (RegA). Pairing verified by single cell VDJ analysis of cells from the same sample (both heavy and light chains in each pair were present in both datasets).

(B) Total number of IGH clones in Patient1 Spatial VDJ dataset at different minimal cut-offs of spots per clone. Of note, the most abundant IGH sequence in each IGH clonal family was selected to account for multiple *related* IGH sequences pairing with the identical light chain. Therefore, only the most abundant heavy chain sequence in each clonal family will be considered for pairing.

(C) The number of correct versus incorrect pairs for Patient1 predicted by *repair* at different cut-off values for the pairing score. Each panel represents data that is thresholded at different numbers of spots per IGH clone. A spot cut-off (n=10) and pairing score threshold (0.35) were selected for subsequent analyses.

(D) Table showing the results from *repair* for the Patient2 Spatial VDJ dataset (verified against the single cell VDJ data from the same patient).

(E) Representative example of a *de novo* predicted receptor pair for Patient2 (RegA). See Fig. 4.

**fig. S24**

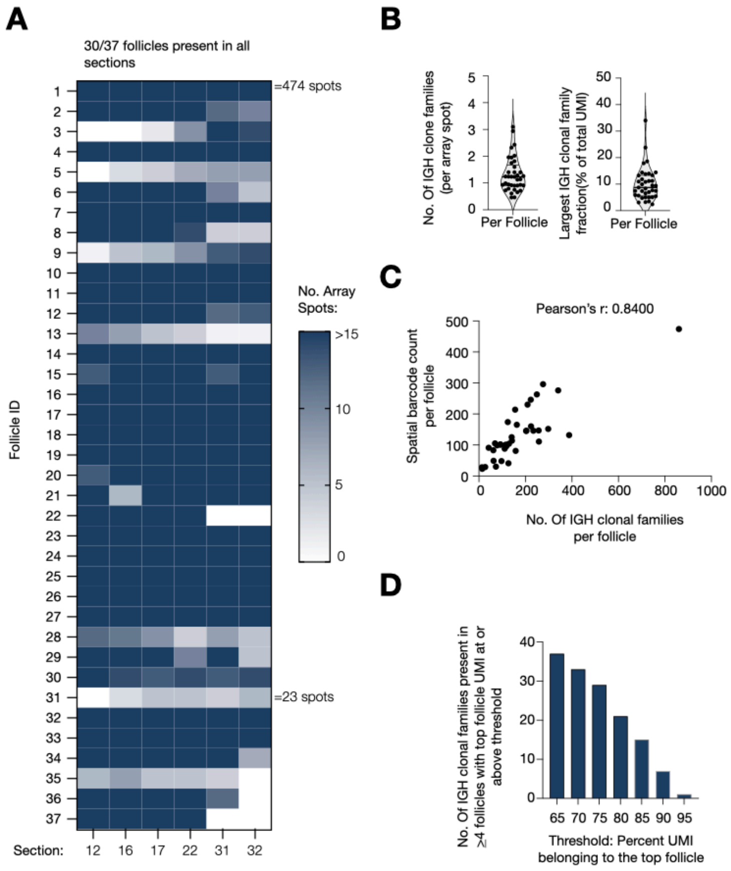


**fig. S24. Spatial VDJ defines follicular B cell clonal dynamics in human tonsil tissue.**
(A) Heatmap of follicle spot count per tonsil tissue section (values larger than 15 are marked as the same color, see scale). The largest (Foll1) and smallest (Foll31) follicles (and their spot counts) are annotated on the heatmap.
(B) Density (count/array spot) of IGH clonal families per follicle (left) and largest IGH clone family fraction per follicle (right).
(C) Follicle size (i.e. array spot count per follicle) versus the number of IGH clonal family count per follicle. Pearson's coefficient (r) is denoted on the graph.
(D) The number of IGH clonal families that have more than the indicated UMI threshold belonging to a *single* follicle. Only IGH clonal families present in four or more follicles are included in the analysis. See Fig. 5.

**fig. S25**

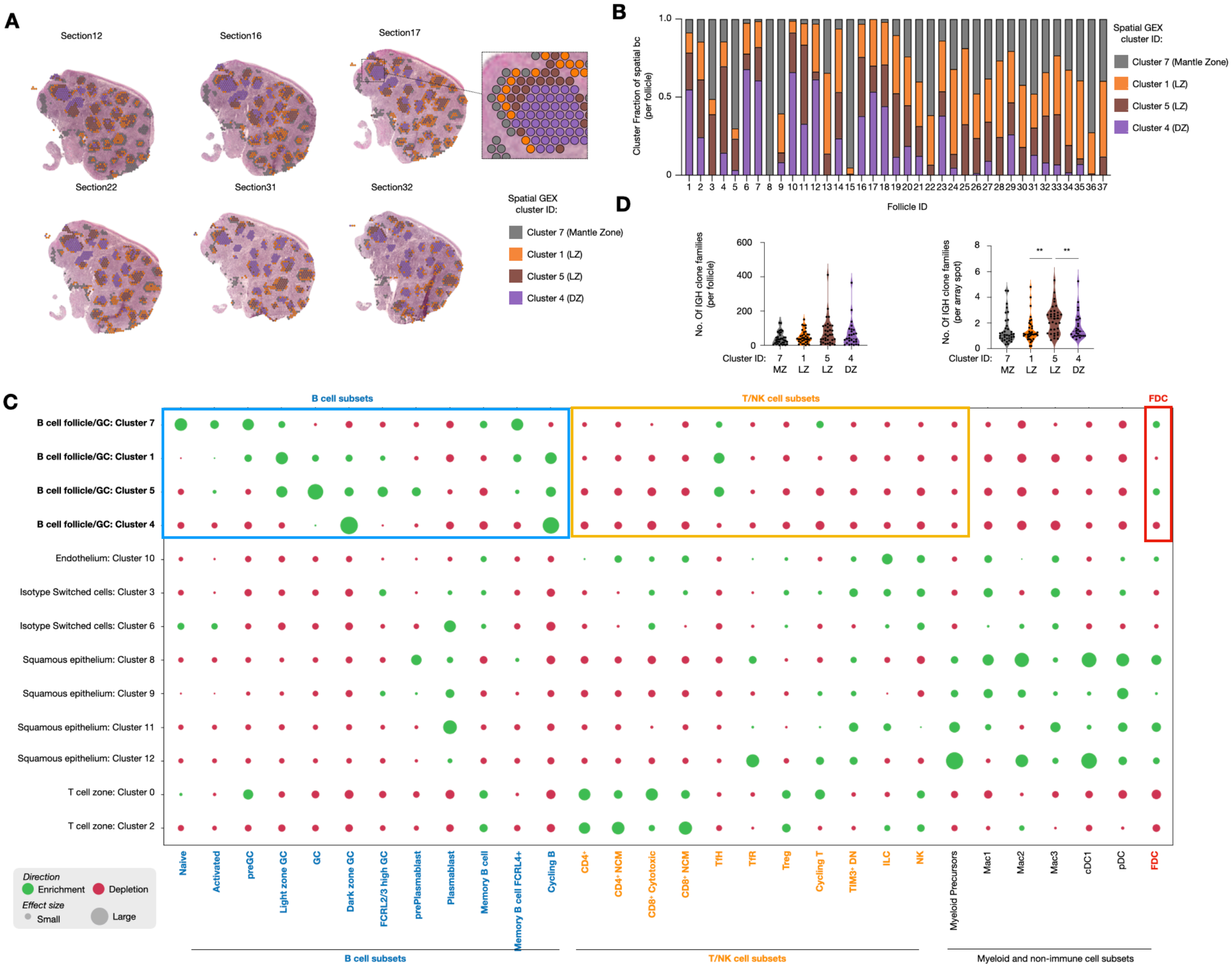


**fig. S25. Spatial VDJ analysis defines intra-follicular regions at high resolutions.**
(A) Spatial GEX follicular cluster distribution across follicles and tonsil tissue section. Predicted light zone (LZ) and dark zone (DZ) are denoted as cluster 1 and 5 versus 4, respectively, and mantle zone as cluster 7. See fig. S7 for differentially expressed genes.
(B) Spatial GEX cluster proportion across individual follicles.
(C) Single cell deconvolution (*stereoscope*) of tonsillar cell subsets across all individual Spatial GEX clusters. The clusters belonging to B cell follicles/GCs are bold. The B cell subsets are listed with respect to their differentiation status, with naive cells to the left. Note the unique enrichment of Tfh in the B cell follicle/GC clusters. Abbreviations: B - B cells; GC - Germinal Center; T/NK - T or NK cells; NCM - Non-Central Memory; TIM3$^{+}$ DN - Double Negative; Tfh - T follicular Helper cells; TfR - T follicular Regulatory cells; Treg - Regulatory T cells; ILC - Innate Lymphoid Cells; Mono - Monocytes; Mac - Macrophages; DC - Dendritic Cells; cDC - conventional DC; PC - Plasma cells; pDC - plasmacytoid Dendritic cells; FDC - Follicular Dendritic Cells.
For these panels: Cluster annotation was done based on several criteria: i) shared expression of

GC-associated genes, including *SUGCT, GMDS,* and *SERPINA9*, ii) enriched gene expression of light vs dark zone-associated genes in c1/c5 vs 4 (fig. S7F), and iii) single cell spatial decomposition data showing enrichment/depletion of light vs dark zone-associated B cell states as well as Tfh enrichment in the light zone. Cluster 7, which was defined as the mantle zone and the 'outermost' layer of the follicles/GCs, had enriched levels of *CXCL13* (expressed by FDC and Tfh), *IGHD, CD19,* and *BANK1* expression (fig. S7F), suggesting the enrichment of naive, unswitched mature B cells in this area. Single cell decomposition corroborated these findings; naive, activated, and preGC B cells were particularly enriched within c7. These observations are consistent with cluster 7 being an initial site of B cell activation, prior to GC entry.

fig. S26

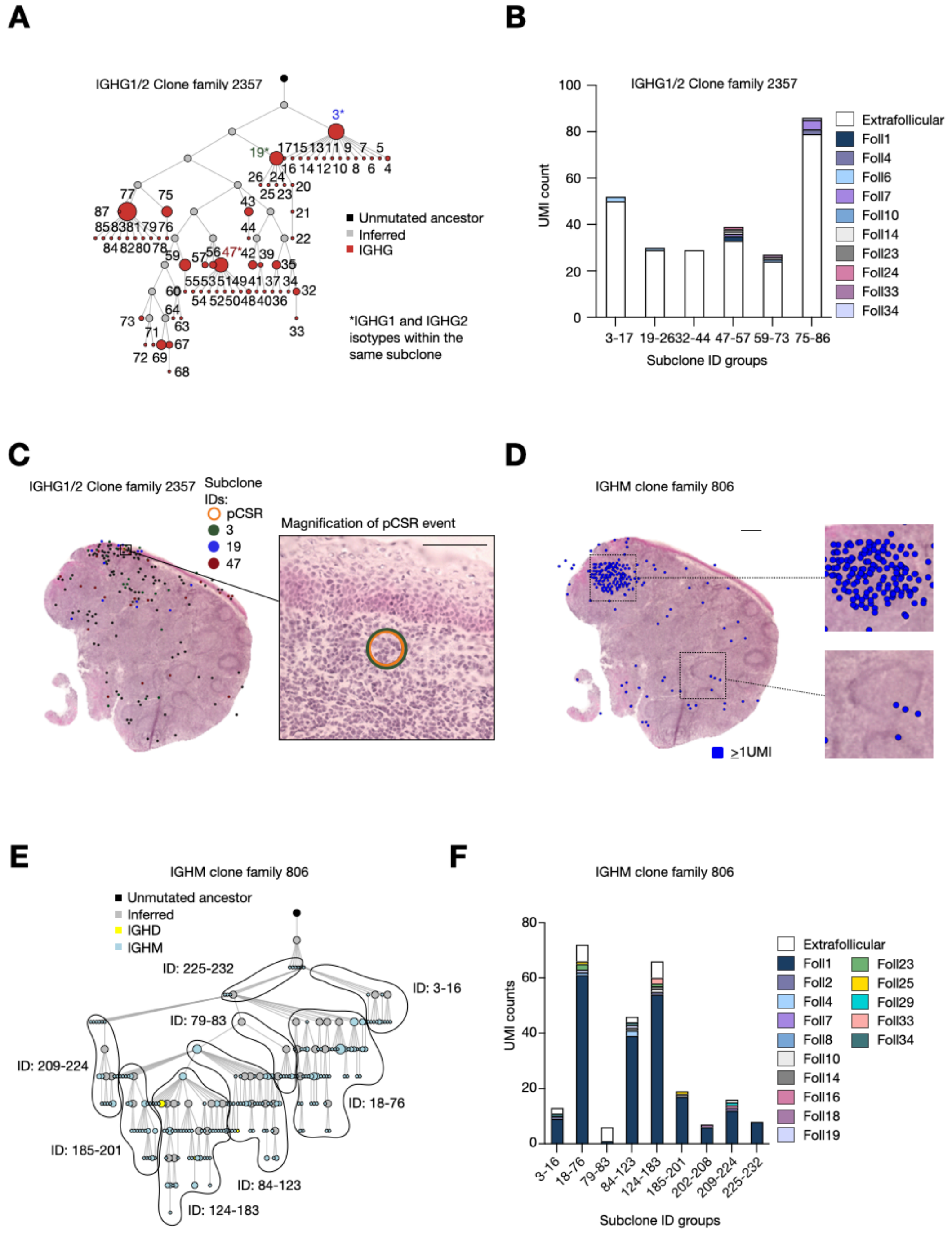


**fig. S26. Spatial VDJ delineates spatial IG clonal evolution.**

(A) Lineage tree of clone family 2357 containing putative Class Switching Recombination (pCSR) event.

(B) IGHG1/2 clonal family 2357 follicular distribution expressed as UMI count per subclone per location. Extra-follicular location is denoted in white.

(C) Spatial distribution of clone family 2357. pCSR event highlighted in orange.

(D) Spatial distribution of clonal family 806 with zoom-ins on two different follicles (the most expanded IGH clonal family based on UMI counts within the LR-Spatial VDJ data).
(E) Phylogenetic tree of IGHM clonal family 806
(F) IGHM clonal family 806 follicular distribution expressed as UMI count per subclone per location. Extra-follicular location is denoted in white.
